# Integrated transcriptome and lineage analyses reveal novel catecholaminergic cardiomyocytes contributing to the cardiac conduction system in murine heart

**DOI:** 10.1101/2022.11.04.515095

**Authors:** Tianyi Sun, Alexander Grassam-Rowe, Zhaoli Pu, Huiying Ren, Yanru An, Xinyu Guo, Wei Hu, Ying Liu, Yangpeng Li, Zhu Liu, Kun Kou, Xianhong Ou, Tangting Chen, Xuehui Fan, Yangyang Liu, Tu Shu, Yu He, Yue Ren, Ao Chen, Zhouchun Shang, Zhidao Xia, Lucile Miquerol, Nicola Smart, Henggui Zhang, Xiaoqiu Tan, Weinian Shou, Ming Lei

**Author notes:** Joint First Authors (Tianyi Sun, Alexander Grassam-Rowe, Zhaoli Pu). Lead Contact (M.L.). Correspondence (M.L.), (W.S.), (X.T.).

## Abstract

Cardiac conduction system (CCS) morphogenesis is essential for correct heart function yet is incompletely understood. Here we established the transcriptional landscape of cell types populating the developing heart by integrating single-cell RNA sequencing and spatial enhanced resolution omics-sequencing (Stereo-seq). Stereo-seq provided a spatiotemporal transcriptomic cell fate map of the murine heart with a panoramic field of view and in situ cellular resolution of the CCS. This led to the identification of a previously unrecognized cardiomyocyte population expressing dopamine beta-hydroxylase (*Dbh*^+^-CMs), which is closely associated with the CCS in transcriptomic analyses. To confirm this finding, genetic fate mapping by using *Dbh*^Cre^/Rosa26-tdTomato reporter mouse line was performed with Stereo-seq, RNAscope, and immunohistology. We revealed that *Dbh*^+^-derived CMs first emerged in the sinus venosus at E12.5, then populated the atrial and ventricular CCS components at E14.5, with increasing abundance towards perinatal stages. Further tracing by using *Dbh*^CFP^ reporter and *Dbh*^CreERT^/Rosa26-tdTomato inducible reporter, we confirmed that *Dbh*^+^-CMs are mostly abundant in the AVN and ventricular CCS and this persists in the adult heart. By using *Dbh*^Cre^/Rosa26-tdTomato/Cx40-eGFP compound reporter line, we validated a clear co-localization of tdTomato and eGFP signals in both left and right ventricular Purkinje fibre networks. Finally, electrophysiological optogenetic study using cell-type specific Channelrhodopsin2 (ChR2) expression further elucidated that *Dbh*^+^-derived CMs form a functional part of the ventricular CCS and display similar photostimulation-induced electrophysiological characteristics to Cx40^CreERT^/ChR2-tdTomato CCS components. Thus, by utilizing advanced transcriptomic, mouse genetic, and optogenetic functional analyses, our study provides new insights into mammalian CCS development and heterogeneity by revealing novel *Dbh*^+^-CMs.

**Highlights:**

- Stereo-seq provided a spatiotemporal transcriptomic cell fate map of the murine heart with a panoramic field of view and in situ cellular resolution of the CCS.
- Established the transcriptional landscape of cell types populating the developing murine heart.
- Revealed previously unreported catecholaminergic cardiomyocyte populations contributing to the developing and mature murine cardiac conduction system.

## Introduction

The cardiac conduction system (CCS) plays an essential role in cardiac physiological function by initiating and coordinating cardiac contraction. The CCS comprises a set of distinct and specialized components, including the pacemaker sinoatrial node (SAN), the coordinating atrioventricular node (AVN) and the rapid transmission ventricular His–Purkinje system. Failure in correct development of CCS components leads to several fast (e.g., Wolff-Parkinson-White syndrome) and slow dysrhythmic (atrioventricular block and sick sinus syndrome) diseases^1^. Gaining insight into the morphogenesis and maturation, and the physiological nature of cell-type heterogeneities of the CCS during the developmental process is critical for understanding the pathogenesis of these dysrhythmic diseases^1^. However, our understanding of when and how diverse cell types arise to form different components of the CCS remains limited^2^.

In murine heart, pacemaker activity can be detected as early as embryonic day (E)8.0^3^. However, the formation of the CCS is not completed until perinatal stages^2,4^. The origin of the cell, cellular differentiation, and the maturation of the various components of the CCS, in particular the ventricular CCS, are incompletely understood. In general, it has been widely accepted that the Purkinje fiber (PKJ) network arises from developing trabecular cardiomyocytes through endothelial-derived inductive signals^5,6^. Recently, it has also been suggested that a polyclonal PKJ network forms by progressive recruitment of conductive precursors to this scaffold from a pool of bipotent progenitors^7^.

On the other hand, catecholamines, the canonical sympathetic neurotransmitter, are known to be essential for cardiac development: mice deficient in catecholamine synthetic enzymes tyrosine hydroxylase (*Th*)^8^ and dopamine beta-hydroxylase *(Dbh)*^9,10^, died in utero with cardiac developmental defects. There is also mixed evidence around the role of catecholamines in establishing and maintaining normal cardiac rhythm^11–14^. However, the true importance and source of these catecholamines for the CCS development remain uncertain^14^.

Recent refinement of transcriptomic technologies including single-cell RNA sequencing (scRNA-Seq) and spatial enhanced resolution omics sequencing (Stereo-seq) ^15,16^ has facilitated the interrogation of cell populations with increasing classification power, revealing and characterizing novel cell types. Several studies have applied such approaches, particularly scRNA-Seq, to investigate cardiogenesis, focusing on subpopulations of cells isolated using predefined genes^17–20^. Given the complexity and incomplete understanding of CCS morphogenesis and maturation, it is important to establish a molecular approach that enables simultaneous analysis of global organ-wide spatial gene expression patterns without biasing against cellular heterogeneity. We thus co-opted Stereo-seq, a newly established high-resolution large-field spatially resolved transcriptomics technique^15,16^, with scRNA-Seq, to map the transcriptional landscape of CCS formation and maturation in the murine heart. This led to the identification of an unreported cardiomyocyte population expressing *Dbh*. Our subsequent lineage tracing analysis revealed that *Dbh*^+^-derived CMs first emerged in the sinus venosus at E12.5, subsequently contributed to both atrial and ventricular CCS components in the process of cardiogenesis and became more prominent in the CCS from perinatal stages to adults. Using optogenetic electrophysiological interrogation of *Dbh^Cre^*/ChR2-tdTomato mice, we further confirmed that *Dbh*^+^-derived CMs function as part of the ventricular CCS. Collectively, our data provide a transcriptomic landscape of the CCS in the murine heart which reveals *Dbh*-expressing cardiomyocytes contributing to the formation and function of CCS in the mammalian heart.

## Results

### 1) Integrated transcriptomic analyses identified a population of *Dbh+* cardiomyocytes in the murine heart

We first sought to characterize the cellular composition of the developing murine heart from middle-to-late embryonic (E8.5, E10.5, E12.5, E14.5, E16.5) and postnatal (P3) stages, by combining scRNA-Seq and Stereo-seq (Figure 1a). Our workflow combined current best practices with recent computational advances (Figure 1a and Figure S1).

**Figure 1.**
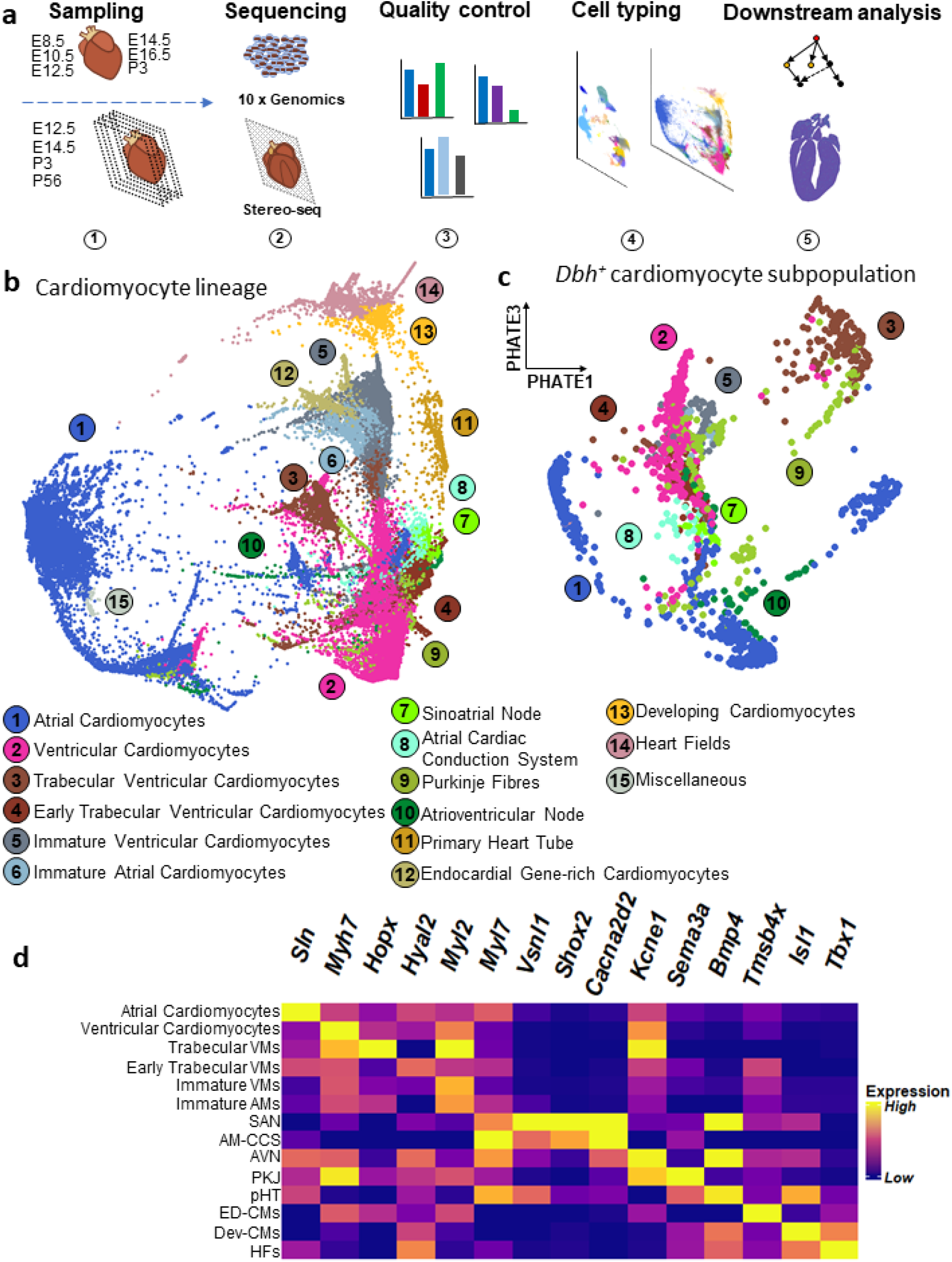
Single-cell RNA sequencing identified a *Dbh*^+^ subpopulation within the cellular landscape of the murine cardiomyocyte lineage. a. Summary of workflow: whole embryos (E8.5, E10.5) or whole hearts (E12.5, E14.5, E16.5, P3) were sampled for single cell RNA sequencing. Slices of whole heart from E12.5, E14.5, and P3 were sampled for Stereo-seq spatial transcriptomics. We used 10X Genomics and Stereo-seq for sequencing. We performed quality control before we clustered and labelled cells based on their gene expression. We performed a range of downstream analyses with both single-cell RNA sequencing and Stereo-seq data. b. PHATE plot of the cellular landscape of the cardiomyocyte lineage. Numbered labels indicate the cell type with the corresponding colour on the PHATE plot, with full names below. (See Supplementary Data 2 for interactive version). c. PHATE plot of the *Dbh^+^* cardiomyocyte subpopulation in the developing murine heart. Numbered labels indicate the cell type with the corresponding colour on the PHATE plot, with full names below and are common with panel b. (See Supplementary Data 3 for interactive version). d. Heatmap demonstrating selected marker genes for each population identified in the cardiomyocyte lineage. Yellow indicates high expression and purple indicates low expression. Gene expression values were normalised across heatmap rows, and then across columns. Abbreviations: VMs – ventricular cardiomyocytes; AMs – atrial cardiomyocytes; SAN – sinoatrial node; AM-CCS – atrial cardiac conduction system; AVN – atrioventricular node; PKJ – Purkinje fibres; pHT – primary heart tube; ED-CMs – endocardial gene-rich cardiomyocytes; Dev-CMs – developing cardiomyocytes; HFs – heart fields.

We established the major cell populations across the developing murine heart. With the use of scRNA-Seq, by 10 ×Genomics Illumina HiSeq PE150 (Figure 1a), following pre-processing and library quality control, our data had mean UMI and gene counts of 6417 and1482 respectively across 175237 cells in our scRNA-Seq (Figure S2). We identified a range of cell types across the developing heart and embryo (Figure S3). Initial analysis suggested that the scRNA-Seq identified 4 overarching populations – early embryonic tissue (e.g., endodermal *Afp*^hi^, mesodermal *Prrx1*^hi^, or neural *Sox2*^hi^), mixed non-myocyte tissue (*Col1a1*^hi^, *Fbln2*^hi^, *Emcn*^hi^, *Upk3b*^hi^, *Lmod1*^hi^), and ventricular (*Myl2*^hi^, *Myl7*^lo^, *Myh7*^hi^, *Kcne1^hi^*) and atrial (*Myl7*^hi^, *Myh6*^hi^, *Nppa*^hi^, *Sln*^hi^) cardiomyocyte populations across 26 cell types (Figure S3a, Figure S4, Figure S5a-d, Figure S6a, Data S1). Thus, our scRNA-Seq could sample the expected cellular composition of the developing heart at a gross level.

We could discriminate major populations and subpopulations of working and non-working cardiomyocytes across our scRNA-Seq dataset of the developing murine heart. To better understand cardiomyocyte populations in the developing heart, we undertook more extensive analysis of the cardiomyocyte lineage. To focus on identifying heterogeneity within and between differentiated and transitory cell states in the cardiomyocyte-lineage, we selected cell types mapping the expected developmental lineage of cardiomyocytes, from mesodermal cardiac progenitors through to neonatal cardiomyocytes to best discriminate transcriptional similarities and differences. We utilized a recently developed local and global structure-preserving dimensionality reduction technique (PHATE^21^ and “spectral-like” unsupervised clustering to identify 15 cardiomyocyte lineage cell types over n=107705 cells from earlier cardiac progenitors across to more mature cardiomyocyte cell types over the 3 dimensional PHATE space (Data S2), which can be taken as an approximation of pseudotime cell-state progression, given the manifold structure-preserving nature of the PHATE algorithm^21^. We observed the “development” from the mesodermal cardiogenic heart fields (*Hand1*^hi^, *Wnt2*^hi^, *Osr1*^hi^), through primary heart tube (*Acta2*^hi^, *Tagln*^hi^, *Pmp22*^hi^) formation, to immature ventricular (*Myl2*^+^, *Myl7*^lo^, *Actc1*^+^, *Tnni1*^hi^, *Tnni3*^lo^) and atrial (*Actc1*^+^, *Sln*^+^, *Myl7*^hi^) cardiomyocytes, and then through into more mature atrial (*Myl7^hi^*, *Sln^hi^*, *Nppa^hi^*, *Tnni3^hi^*) and ventricular (*Myh7^hi^*, *Myh6^lo^*, *Myl2^hi^*, *Kcne1^hi^*, *Tnni3^hi^*) cardiomyocytes (Figure 1b, Figure 1d, Figure S6b, Figure S7, Figure S8a-c, Data S2, Table S1). Interestingly, we saw that much of the CCS formed clusters distinct in 3D PHATE space from their respective most similar working cardiomyocyte populations (Figure 1b, Data S2). For example, Purkinje Fibres (PKJs) compared with the trabecular ventricular myocardium. With use of the “spectral-like” clustering algorithm^21^, and cluster labelling based on gene expression profiles (Figure 1d, Figure S6b, Figure S9, Data S2, Table S1) we could resolve the CCS from our mixed scRNA-Seq, whereas many prior scRNA-Seq studies^17–20^ did not discriminate separate CCS populations apart from those using CCS-biased sampling techniques^22,23^. Therefore, we used novel analytical approaches to better understand and reveal heterogeneity within the developing mouse cardiomyocyte lineage.

We identified previously unreported dopamine beta-hydroxylase (*Dbh*) expressing cardiomyocytes (*Dbh* ^+^-CMs) populations from within the cardiomyocyte lineage (Figure 1c). We observed *Dbh* expression across both working and non-working (i.e. pacemaker/conducting) cardiomyocyte populations (Figure 1c). The major *Dbh* ^+^-CM cell types were: atrial and ventricular cardiomyocytes; early and more mature ventricular trabecular cardiomyocytes; sinoatrial and atrioventricular nodal cardiomyocytes; and non-specific atrial cardiac conduction system and Purkinje fibres, although we did identify a small number of cells that belonged to earlier developing cardiomyocyte populations (Figure S10, Figure S11, Data S3). Overall, there were n=2591 *Dbh*^+^-CM, at 2.43% of all cardiomyocyte lineage cells from E8.5, through to P3. However, significant numbers of *Dbh* ^+^ cells only became apparent beyond E10.5 (Data S3). We confirmed expression of expected specific pan-cardiomyocyte markers such as *Tnnt2* (Figure S6ci), and CCS-specific markers (Figure S6cii), within the *Dbh*^+^ population. Strikingly, although *Dbh*^+^-CM represented only small percentages of the total working cardiomyocyte populations (AM: 2.13%, VM: 3.90%, VM-trab: 4.18%, and eVM-trab: 2.26%) (Figure S11), the CCS appeared to be over-represented in *Dbh*^+^ cardiomyocytes as percentages of their respective total cardiomyocyte lineage numbers (PKJ: 12.76%, AVN: 12.14%, SAN: 8.39%, and AM-CCS: 3.82%) (Figure S11). Pearson’s Chi-Squared test for independence confirms that cell type, within cardiomyocytes, and *Dbh* expression (positive or negative) are not independent (Χ^2^ = 1155.2, df = 9, p= 5.8e-243, n=103008). To summarize, we identified previously unreported transcriptomic expression of a catecholamine biosynthetic enzyme within developing murine cardiomyocytes, and its association with the CCS.

To facilitate future hypothesis generation, we also provide a publicly available web resource to visually explore our transcriptional landscapes in a cell-wise manner (Data S1-3), allowing examination of global and local cell distributions, as well as identifying a range of quality control metrics, in the developing murine heart.

### 2) Stereo-seq recapitulated our identification of Dbh-expressing cardiomyocytes across the developing and mature murine heart

To validate the identification of *Dbh*^+^-CMs, we used a new nanoscale resolution spatial transcriptomics technology Stereo-seq^15,16^, to analyze this *Dbh*^+^ cardiomyocyte population, including its spatial distribution, in the developing and mature murine hearts. We established a *Dbh*^+^ cell lineage tracing reporter line, *Dbh*^Cre^/Rosa26-tdTomato mice, for genetic fate mapping of *Dbh*^+^-CMs (Figure 2a). Following sampling, library preparation, sequencing and quality control, we had at least n = 3 sections for hearts from *Dbh*^Cre^/Rosa26-tdTomato mice at E12.5, E14.5, P3, and P56. DNA-labelled nanoballs were binned at 20 x 20 (~14 μm spot diameter) for E12.5, E14.5 and P3, and at 50 x 50 (~35 μm spot diameter) for P56 (Figure S12a).

**Figure 2.**
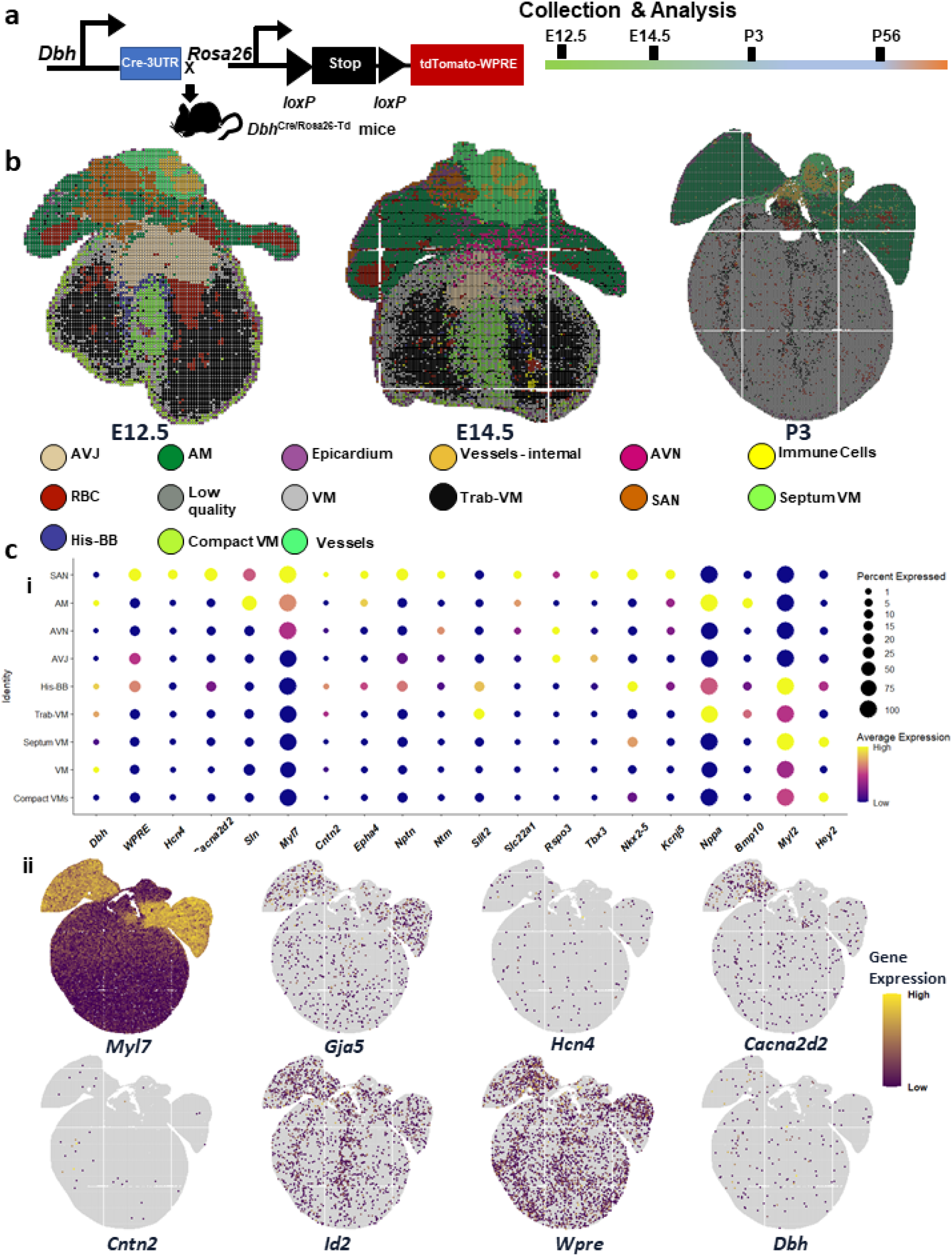
*Dbh* is expressed dynamically across the developing heart, as resolved with nanoscale-resolution Stereo-seq of a *Dbh*^Cre^/Rosa26-tdTomato reporter line. a. Schematic of *Dbh*^Cre^/Rosa26-tdTomato reporter line creation and analysis for spatially-resolved transcriptomics with Stereo-seq. b. Plot of spatially-resolved transcriptomic pixels (each pixel consists of 20×20 from spots of individual diameters between 500 or 715 nm) identified across hearts from E12.5, E14.5, and P3 mice, coloured by cell type identified through unsupervised clustering with corresponding cell types coloured below. c. i) Dot plot describing *Dbh* and *Wpre* (a tdTomato construct-derived transcript) expression, with other key marker genes, across major cardiomyocyte populations identified through unsupervised clustering of E12.5, E14.5, and P3 Stereo-seq data. Dots are sized by percentage of cells expressing each respective gene in each respective cell type. Dots are coloured by average expression of the respective gene in the respective cell type, with yellow indicating higher expression, and purple lower expression. ii) Spatially-resolved plots describing *Dbh* and *Wpre* (a tdTomato-derived transcript) expression, with other key marker genes, across a slice from a P3 mouse heart sequenced with Stereo-seq. Yellow indicates higher expression and purple lower expression. Abbreviations: VM – ventricular myocardium; Trab-VM – trabecular ventricular myocardium; SAN – sinoatrial node; AVN –atrioventricular node; AVJ – atrioventricular junction; AM – atrial myocardium; His-BB – His-bundle branch; AM-CCS – atrial cardiac conduction system; PKJ – Purkinje fibres; VM-CCS – ventricular cardiac conduction system.

Stereo-seq provided a spatiotemporal transcriptomic map of the murine heart with a panoramic field of view and in situ cellular resolution of the CCS. First, we sought to confirm Stereo-seq could faithfully recapitulate known biology in the specific context of the developing murine heart. We were able to resolve the expected changes in gene expression and spatial distribution across both the developing and adult mouse heart (Figure S13a-c). Confident in examining global changes in gene expression across our Stereo-seq dataset, we sought to ensure this matched with the appropriate cell type co-expression of genes. We identified a range of working and non-working cardiomyocyte populations in our analysis of stages E12.5, E14.5, and P3, alongside expected non-cardiomyocyte populations (Figure 2b-c, Figure S14). We omitted P56 from cell type classification to ensure comparability with our scRNA-seq data, which does not extend beyond P3. To minimize the potential of bias in our cell type classification; we also used additional supervised learning workflows to classify cells using our scRNA-Seq dataset as a reference for cell-type specific gene expression(Table S2)^24,25^. Overall, this supervised classification largely agreed with our cell types identified (Figure S15). However, we note that “AM-CCS”, mapped in a non-specific manner across the atrial myocardium (Figure S15), likely reflecting the weaker expression of CCS markers in AM-CCS, compared to the SAN/AVN, but with expression of more generic atrial markers (Figure S9). We believe the AM-CCS likely represents either some transitional sinus node population, or indeed inter-atrial conduction pathways (Figure S9). Those cell types with imperfect integration often appear to have mapped to cell types with expectedly similar transcriptomic signals; analysis of the robust cell type decomposition (RCTD) scores for certain populations suggests that these cell types may still be identified by this supervised learning technique, but that the final highest predictive score for each cell might be obscuring the complexity of similar or even overlapping cell types being captured in each bin (Figure S15). In sum, Stereo-seq could resolve cell types across the developing heart, down to a quasi-single cell resolution, within the context of a whole-organ sample.

Stereo-seq supported our identification of *Dbh* expression in cardiomyocytes across the developing heart. Following our cell type identification, we proceeded to investigate *Dbh* real-time expression and lineage tracing. We could readily observe *Dbh* expression across the myocardium at all stages observed (Figure 2ci-ii, Figure S13d). We recapitulated our earlier findings in our scRNA-Seq data that *Dbh* expression was across both working and non-working cardiomyocyte cell types from E12.5 through to P3 (Figure 2ci). However, *Dbh* expression appeared particularly high in the His/Bundle Branch part of the proximal ventricular CCS (Figure 2ci). Concordantly, we saw *Dbh* spatial expression overlapping with a number of ventricular CCS markers, such as *Cntn2*, *Cacna2d2*, *Gja5*, *Hcn4*, and *Id2*, as shown in a representative slice from P3 in Figure 2cii. The spatial co-localisation of *Dbh* and CCS marker genes identified by Stereo-seq provides additional support to the association of *Dbh* with the developing CCS, particularly in the ventricular CCS.

Stereo-seq enabled spatiotemporal genetic fate mapping of *Dbh^+^-*derived cells across the murine heart. We also sought to understand the distribution of the *Dbh* lineage beyond real-time expression by using our *Dbh*^Cre^/Rosa26-tdTomato reporter line. Unfortunately, tdTomato-coding sequences in our generated libraries were not readily detectable due to the use of 3’-end 100 base pair sequencing, thus the expression levels of *Wpre* (expressed at 3’-end of the tdTomato reporter cassette construct) (Figure 2a) were used as a proxy for tdTomato expression in our sequencing data, which represent *Dbh*^+^-derived and potential *Dbh*^+^ cells. Overall, we identified the *Dbh*^+^-derived cardiomyocyte (*Dbh*^+^-derived CMs) lineage was associated with the CCS in a somewhat similar manner as real time *Dbh* expression. *Wpre* expression was highest in the CCS (Figure 2ci). However, we did also observe *Wpre* expression across the non-working and working myocardium from E12.5 through to P56 (Figure 2cii, Figure S13d). Of note, *Wpre* expression signal appeared stronger in the SAN than in the ventricular CCS, which was the opposite of *Dbh* expression observed in our Stereo-seq data and scRNA-Seq data (Figure 2ci, Figure S11). This likely reflects differences in the temporal expression dynamics between endogenous *Dbh* and the reporter construct, such that cells in the atrial CCS (e.g. SAN), might have ceased to express *Dbh* but continue to express *Wpre* as a descendant of a once *Dbh*^+^ progenitor, or in their own changing transcriptional program. The developmental multi-stage and spatially-resolved tracking of our tdTomato lineage reporter by Stereo-seq supported the expression of *Dbh* across cardiomyocyte lineages, but with particular association with the CCS.

Cardiomyocyte *Dbh* expression persisted into the adult murine heart. To confirm whether these developmental *Dbh*^+^-and/or *Wpre*^+^-cardiomyocyte populations were still present in adulthood, we examined adult (P56) heart slices prepared from the *Dbh*^Cre^/Rosa26-tdTomato mice through Stereo-seq. At P56, *Wpre*^+^ cardiomyocytes were numerous in the right atrium, SAN, AVN, and His-Purkinje network similar to in development (Figure S13d). A number of *Dbh*^+^-CMs were also identified by Stereo-seq in adult mouse heart slices, with similar spatial expression profiles as a number of ventricular CCS markers such as *Hcn4*, *Cacna2d2*, *Cntn2*, and *Slit2* (Figure S13b-d) supporting the persistence of *Dbh*^+^-CMs into the mature adult heart beyond the postnatal period.

### 3) Genetic fate mapping and multiplexed nucleic acid in situ hybridization support that *Dbh^+^*-CMs, and*Dbh^+^-*derived CMs populate the developing CCS

To validate the transcriptomic identification of *Dbh*^+^-CMs, genetic fate mapping was conducted in developing and postnatal hearts. We developed three distinct genetic mouse models for such a purpose: *Dbh*^Cre^/Rosa26-tdTomato, *Dbh*-Knockin-CFP (*Dbh*^CFP^) and *Dbh*^CreERT^/Rosa26-tdTomato inducible lines. Whole embryos (E8.5, E9.5, E10.5, E12.5, E14.5, n=5 embryos per stage) or isolated hearts (E12.5, E13.5, E14.5, E16.5, P3, P56, n=5 hearts per stage) were analyzed by using multiplex nucleic acid in situ hybridization (RNAscope), immunohistological staining, and confocal microscopic imaging.

Genetic fate mapping was initially conducted on the *Dbh*^Cre^/Rosa26-tdTomato mouse line to validate the findings from our Stereo-seq analysis on the same mouse line. The tdTomato fluorescence was first observed in the dorsal neural tube at E10.5, likely reflecting neural expression in the developing CNS. From E10.5 to E12.5, tdTomato-expressing cells branched into groups migrating to the heart region via the pharyngeal arches (Figure 3a). At E12.5, tdTomato expression was observed in the sympathetic ganglion chain adjacent to the spinal cord, and, of particular interest, within the sinus venosus (SV, the primordial SAN) and right atrium (RA) (Figure 3b and Figure S16a). Thus, supporting the population of the developing CCS with *Dbh^+^-*derived CMs, as denoted by tdTomato expression at the protein level within the primordial CCS.

**Figure 3.**
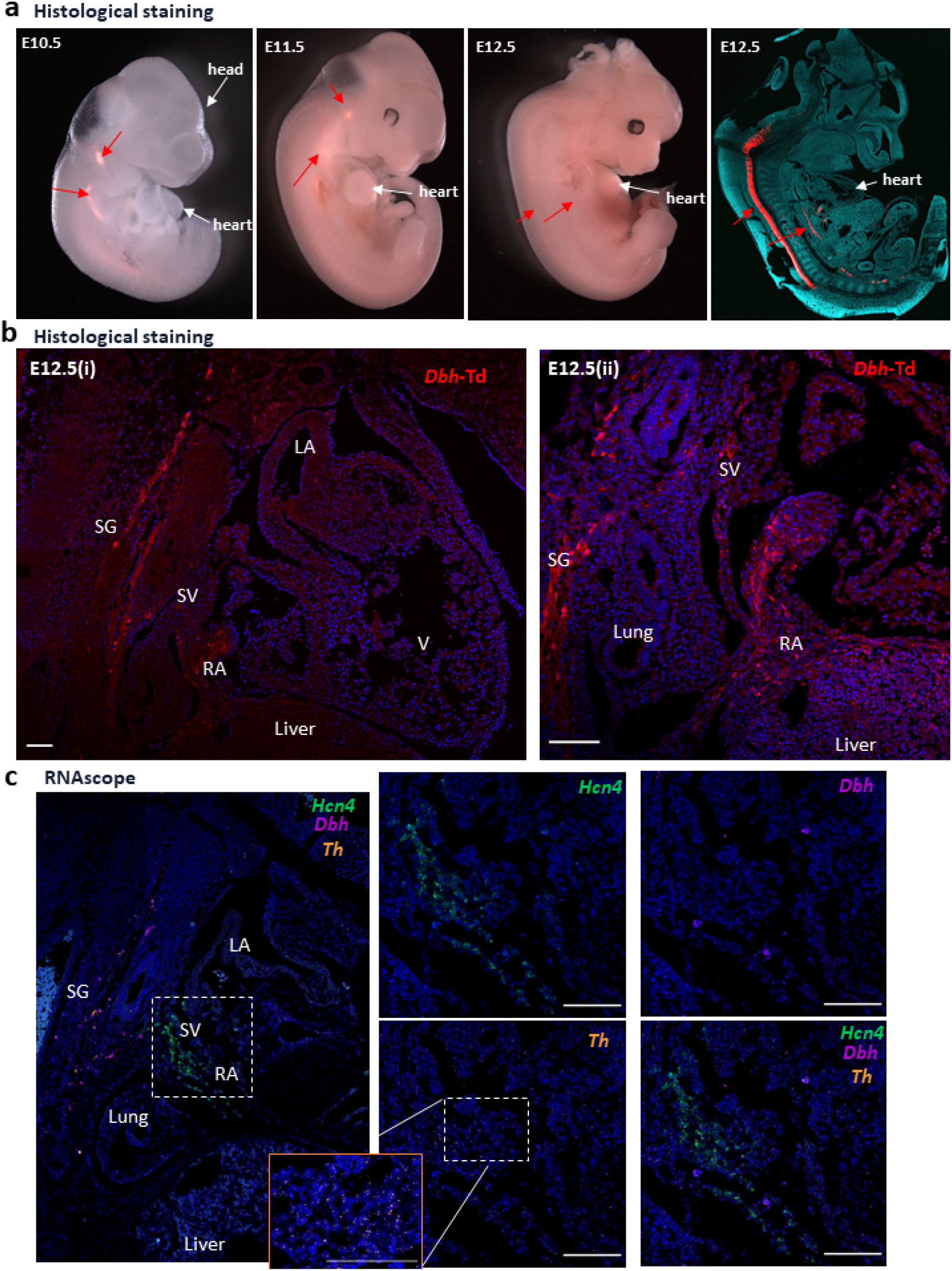
Prospective fate mapping analysis of *Dbh^Cre^*/Rosa26-tdTomato heart throughout the developmental stages. a. Representative gross images of tdTomato-expressing signals from E10.5 to E12.5 mouse embryos (left 3 panels) and an image of tdTomato-expressing signals of a sagittal section of a E12.5 embryo (far right panel). b. Representative images showing tdTomato-expressing cells in RA and SV region of a sagittal section of E12.5 embryos (i) and magnified SG and RA area (ii). c. Representative RNAscope images of *Dbh* (purple), *Hcn4* (green) and *Th* (orange) in comparable region of RV and SV on adjacent sagittal section of E12.5 embryo in b. Despite *Th*+ signals are in SV, there is no overlap with *Dbh^+^* and *Hcn4^+^* signals. Scale bar: 100 μm Red Arrowed: tdTomato-expressing cells SG: Sympathetic Ganglia SV: Sinus Venosus LA: Left Atrium RA: Right Atrium V: Ventricle

We then sought to understand whether these cells remained actively transcribing *Dbh*. Unfortunately, antibodies against the Dbh protein provided poor signals with little specificity. Thus, we utilized RNAscope to examine *Dbh* expression in cardiac tissue at the mRNA level. *Dbh* expression was observed in the SV and RA regions at E12.5 (Figure 3c). Furthermore, RNAscope demonstrated abundant *Hcn4*, the major gene associated with the central CCS^26^, expression in the SV and RA regions, partial co-localization with *Dbh* in the same region (Figure 3c), supporting that *Dbh* expression is associated with cardiac pacemaker tissue. Then, we sought to understand whether this *Dbh* expression arises from sympathetic neurons innervating the SV and RA. The expression of Tyrosine hydroxylase (*Th*), the major marker for sympathetic nerve system, was observed in the sympathetic ganglion chain correlating with the tdTomato signal from the sympathetic ganglia shown in Figure 3b and Figure S16b. However, notably, there is no overlap between *Dbh* and *Th* in SV and RA, indicating that the *Dbh* signal was not innervated cell in origin. These data in sum support that the *Dbh* is actively expressed by cardiomyocytes in the developing murine heart, and that these are associated with the CCS.

To further determine the relationship between *Dbh^+^* cells and the cardiac conduction system, RNAscope was applied on embryonic hearts at E12.5 (Figure 4). The expression and colocalization of *Dbh* and CCS markers *Cacna2d2*, *Hcn4*, *Id2*, *Shox2*, *Tbx18* were investigated by co-staining by multiplexed RNAscope in wild-type murine hearts from E12.5 to postnatal stages (Figure 4 and Figure S17). At E12.5, *Dbh* is abundant in the SV and AVN and is well overlapped with *Cacna2d2*. while it is less abundant in PKJ but is overlapped with *Hcn*4 in this region (Figure 4a). At this stage, the AVN region marked by *Id*2 expression also demonstrates a co-expression of *Id2* and *Dbh* (Figure 4b). At E14.5, *Cacna2d2*, *Id2*, *Shox2*, *Tbx18* and *Hcn4* are overlapped with *Dbh* in various degrees in different parts of CCS, illustrating the existence of *Dbh^+^-*cells in CCS regions at this stage (Figure S17a-b). Especially, Abundant *Dbh* is observed in the SAN and RA regions where *Cacna2d2, Id2* and *Hcn4* signals marked, shown in Figure S17b. At P3, notable *Dbh* signals were observed in SAN and both left and right PKJ, together with CCS marker *Cacna2d2* shown in Figure S17c. Taken together, these findings strongly suggest that *Dbh^+^* cells are potentially involved in CCS development and function.

**Figure 4.**
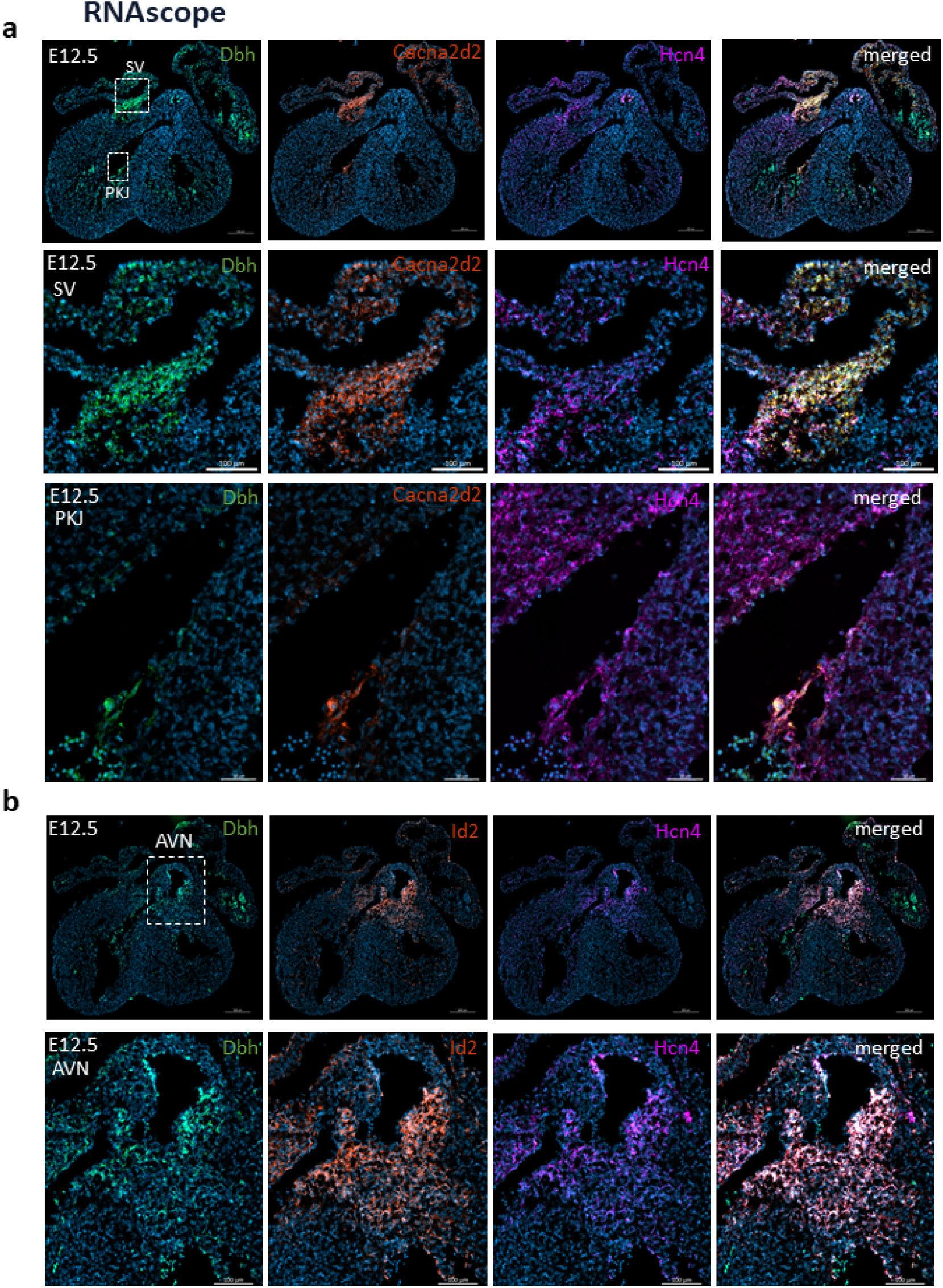
The use of RNAscope to confirm the expression pattern of *Dbh* and CCS markers in E12.5 hearts. a. Representative RNAscope images of the distribution of *Dbh* (green), CCS markers *Cacna2d2* (red) and *Hcn4* (magenta) in a whole field of E12.5 heart. The second row shows SV and the third row show PKJ. b. Representative RNAscope images of the distribution of *Dbh* (green), CCS markers *Id2* (red) and *Hcn4* (magenta) in a whole field of E12.5 heart. The second row shows AVN. SV: Sinus Venosus AVN: Atrioventricular node PKJ: Purkinje fiber network

The cell type identity and localization of *Dbh^+^-*derived cells and *Dbh^+^-*cells were further determined and delineated at E14.5, postnatal (P3) and adult (P56) stages. *Dbh*^Cre^/Rosa26-tdTomato mouse line labelled historically expressed *Dbh*^+^-cells as tdTomato fluorescence. We found that tdTomato-expressing cells were highly enriched in the cardiac conduction system and ventricular trabecular regions throughout these stages (n=5 hearts examined per stage). More specifically, tdTomato-expressing cells were detected in the AVN and His bundle regions (Figure 5a-c). Furthermore, some tdTomato expressing cells were also detected in the ventricular trabecular region (Figure 5a) across the heart development, which is consistent with the developmental origin of PKJ^5,6^. By staining with the myocyte marker α-Actinin, we further confirm that the majority of tdTomato-expressing cells are cardiomyocytes, thus further confirming the identity of the cells as *Dbh^+^*-derived CMs (Figure 5a-d).

**Figure 5.**
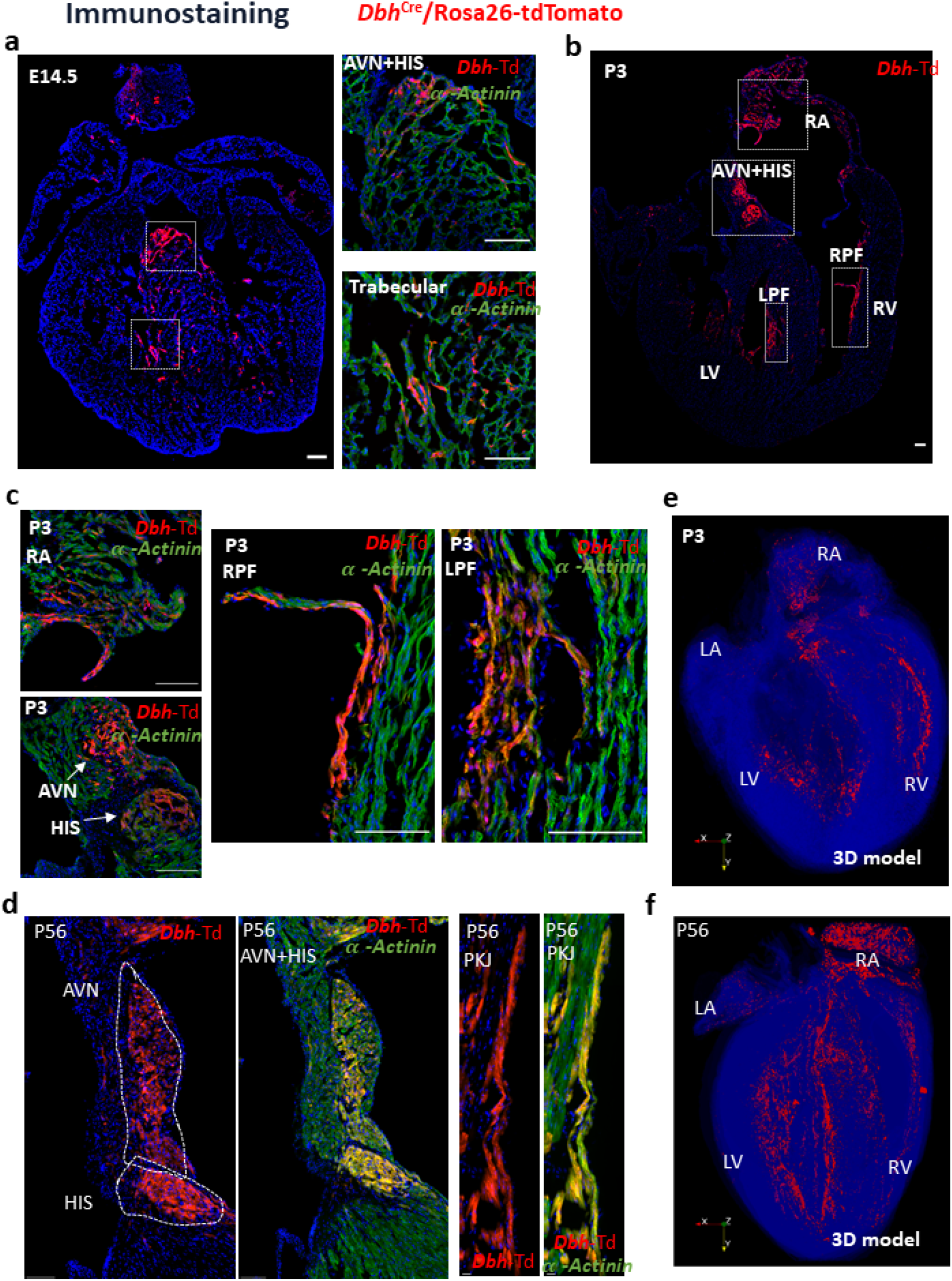
Fate mapping of *Dbh*^+^-derived cardiomyocytes in the developing and postnatal hearts by using *Dbh*^Cre^/Rosa26-tdTomato reporter. a. Representative images of immunostaining showing the co-expression of tdtomato (red) and α-actinin (green) illustrating the distribution of *Dbh*+-derived CMs in AVN and HIS and trabecular regions at E14.5. b. Representative image of tdTomato expressing cells in a four-chamber view section of P3 heart. c. Representative images of immunostaining showing the co-expression of tdTomato (red) and α-actinin (green), illustrating the distribution of *Dbh*^+^-derived cardiomyocytes in RA, HIS, LPF and RPF regions in a heart at P3. d. Representative images of immunostaining showing the co-expression of tdTomato (red) and α-actinin (green) illustrating the distribution of *Dbh*^+^-derived cardiomyocytes in AVN, HIS, PKJ regions in an adult heart. e. The distribution profile of *Dbh*^+^-derived CMs and multi-angle hyperspectral three-dimensional reconstruction results for *Dbh*^+^-derived CMs in a P3 heart sections. All results were written into VTK file format. f. The distribution profile of *Dbh*^+^-derived CMs and multi-angle hyperspectral three-dimensional reconstruction results for *Dbh*^+^-derived CMs in adult (P56) heart sections. All results were written into VTK file format. AVN: Atrioventricular node PKJ: Purkinje fiber network HIS: His-bundle branch LV: Left ventricular RV: Right ventricular RPF: Right Purkinje fiber network LPF: Left Purkinje fiber network RA: Right atria

To further elucidate spatially anatomical characteristics and expression pattern of *Dbh^+^*-derived CMs in postnatal and adult hearts, we interrogated the hearts with imaging and 3D computational image reconstruction. We characterized the spatial distribution of *Dbh^+^*-derived CMs across the whole heart and their structural relationship with specific components of the myocardium in neonatal (P3) and adult (P56) hearts (2-month-old) using *Dbh*^Cre^/Rosa26-tdTomato mouse line. Based on a series of microscopic histological images from P3 and adult hearts, we developed three-dimensional (3D) computational image reconstruction to determine the spatial distribution of *Dbh^+^*-derived CMs in P3 (Figure 5e) and adult (Figure 5f) hearts. The 3D models revealed spatial distribution characteristics of *Dbh^+^*-derived CMs in CCS (i.e., AVN, and His-Purkinje network) as shown in Figure 5e-f, consistent with our earlier Stereo-seq analyses shown in Figure 2d. Multi-angle view of the 3D reconstructions for such spatial distribution of *Dbh^+^*-derived CMs in P3 and adult hearts are also illustrated in videos S1 and S2. From E14.5 to adulthood (Figure S18a-b), *Dbh^+^*-derived CMs were abundant in the CCS regions where sympathetic innervation is enriched, as detected by Th immunostaining, particularly in the adult heart (Figure S18b).

### 4) The spatial distribution of *Dbh^+^*-CMs is closely associated with the ventricular CCS

We then developed a *Dbh^CFP^* mouse line to further reveal the expression and localisation of *Dbh* expressing cells (*Dbh*^+^-cells) in the developing and adult heart. Due to the weak genetic CFP signal, anti-Flag or anti-CFP antibodies were applied to enhance the CFP signal (defined by expressing CFP). As shown in Figure 6a, distinct *Dbh*^+^-CMs, confirmed with cardiomyocyte marker a-Actinin staining, were detected in E14.5 and adult (P56), mostly abundant in AVN and ventricular CCS. We further established *Dbh*^CreERT^/Rosa26-tdTomato inducible reporter mouse line that allows us to map the *Dbh*^+^-CMs under a temporally controlled manner. Adult mice were treated at week 8 with 5-day tamoxifen administration and dissected after one-week post-induction. The tdTomato fluorescence was observed especially in AVN and HIS regions of CCS (Figure 6b) as cardiomyocyte identity validated by cardiomyocyte marker α-Actinin staining, consistently with our results obtained in *Dbh*^Cre^/Rosa26-tdTomato and *Dbh^CFP^* mice lines (Figure 5d and Figure 6a), where sympathetic innervation is enriched, as detected by Th immunostaining as exemplified in Figure S19.

**Figure 6.**
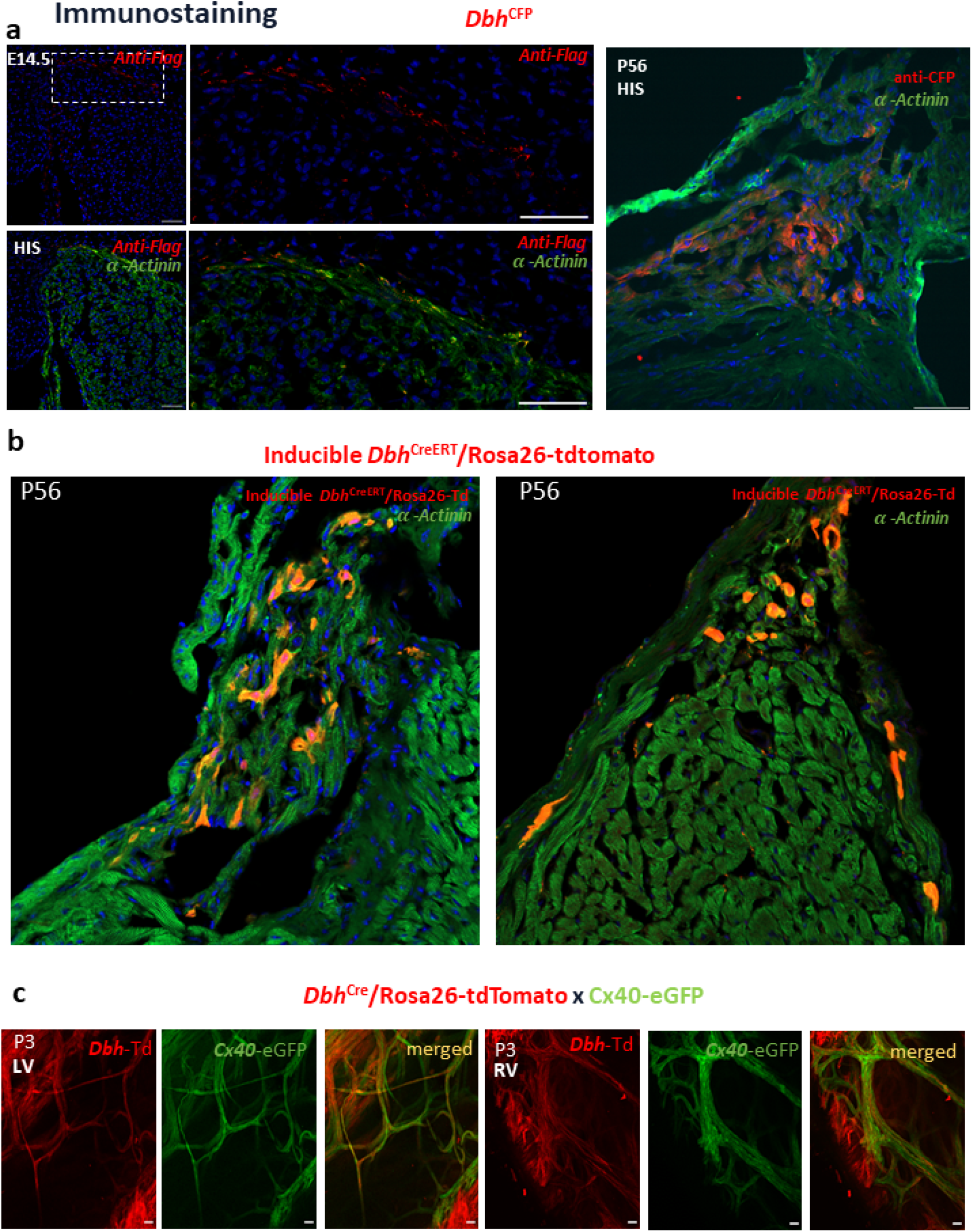
Validation of spatial distribution of *Dbh*^+^-CMs and their close association with CCS. a. Representative images of immunostaining with anti-Flag (red) and anti-CFP (red), plus α-actinin (green) antibodies showing the expression of *Dbh* -CMs in HIS region at E14.5 and adult (P56) by using *Dbh*^CFP^ reporter. b. Representative images in AVN and HIS with α-actinin (green) immunostaining showing the expression pattern of *Dbh*^+^-CMs by using *Dbh*^CreERT^/Rosa26-tdTomato inducible reporter at P56. c. Representative images showing co-localization of *Dbh*^+^-derived CMs and Cx40^+^ cells in PKJ network in both left and right ventricle in *Dbh*^Cre^/ChR2-tdTomato/Cx40-eGFP neonatal mouse hearts.

To further examine if *Dbh^+^*-derived CMs contribute to PKJ network, we then established mouse model *Dbh*^Cre^/Rosa26-tdTomato/Cx40-eGFP by crossing a *Dbh*^Cre^/Rosa26-tdTomato reporter line with Cx40 transgenic eGFP (Cx40-eGFP) line that demarcates the PKJ network. A clear co-localization of tdTomato and eGFP signals was observed in both left and right ventricular Purkinje fibre networks (Figure 6c) in *Dbh*^Cre^/Rosa26-tdTomato/Cx40-eGFP neonatal hearts (n=5 hearts examined). Notably, the *Dbh^+^*-derived CMs (tdTomato-expressing cells) are more extensive than eGFP^+^-cells in the ventricular endomyocardium. The result suggests that *Dbh^+^*-derived CMs form cardiomyocyte populations other than Purkinje conductive cells, which is in agreement with our earlier transcriptomic analyses (Figure 1c, Figure 2 c-d, Figure 5c).

In summary, by using three mouse lineage tracing models and various techniques, despite some minor differences in signal distribution due to the different technologies used, we can consistently validate the identity of the majority *Dbh^+^*-CMs and *Dbh^+^*-derived CMs in the developing, postnatal and mature hearts. Their unique localization in CCS and co-localized with CCS markers are uncovered and verified, strongly suggesting their physiological role in CCS, especially in HIS and PKJ where *Dbh^+^*-derived CMs are found to co-express with Cx40 proteins.

### 5) Electrophysiological characterization of *Dbh^+^-*derived CMs indicates they form a functional part of the ventricular CCS

To elucidate the physiological function of *Dbh^+^*-derived CMs in adult hearts, we interrogated the hearts with optogenetic electrophysiology. We determined the functional role of *Dbh^+^*-derived CMs by selectively expressing light-sensitive Channelrhodopsin-2 (ChR2) channels in these cells by crossing the *Dbh*^Cre^ line with the ChR2-tdTomato line^27^.The resulting line (*Dbh^Cre^*/ChR2-tdTomato) allows the selective stimulation of *Dbh^+^*-derived CMs by activating ChR2 expressed in these cells with photostimulation.

Firstly, we determined if *Dbh^+^*-derived CMs are associated with ventricular CCS, we compared photostimulation-induced electrophysiological characteristics of *Dbh^Cre^*/ChR2-tdTomato hearts with Cx40-CreERT/ChR2-tdTomato and MHC-Cre/ChR2-tdTomato hearts. Cx40-CreERT/ChR2-tdTomato mice have been used as an optogenetic tool for studying ventricular CCS by expressing ChR2 in Purkinje fibers^28^, while MHC-Cre/ChR2-tdTomato line has been used for studying cardiomyocytes in general as a non-selective cardiomyocyte ChR2 expressing mouse model. To enable timely and spatially controlled photostimulation of ChR2 hearts, we used a fiber optic delivering 470-nm light pulses (5 ms), generated by a time-controlled light emitting diode (LED) directed towards the epicardium of Langendorff-perfused hearts in LA, RA, LV and RV regions (Fig.7a). ECGs were then recorded for electrophysiological analysis. We compared the regional responsiveness and QRS waveform characteristics of Langendorff-perfused hearts from three lines measured at sinus rhythm or light pacing at cycle length 100-120 ms. MHC-Cre/ChR2-tdTomato (MHC-ChR2) hearts show uniformed responsiveness to photostimulation in four regions (LA,RA,LV,RV) of the heart with comparable morphology of QRS complex waveforms and durations to the sinus rhythm QRS (Fig.7b.c,d), while *Dbh* (*Dbh*-ChR2) and Cx40 (Cx40-ChR2) hearts show predominate RA and RV responses, but not in LV. As summarized in Figure 7c and exemplified in Figure 7d.e, RV epicardium photostimulation initiates ventricular ectopic-like QRS morphology that is distinct from sinus rhythm QRS morphology. This demonstrates that *Dbh*-ChR2 hearts have a similar photostimulation response to Cx40-ChR2 hearts, in contrast to MHC-ChR2 hearts, in terms of chamber-specificity and RV light pacing (LP)-induced QRS waveform characteristics.

**Figure 7.**
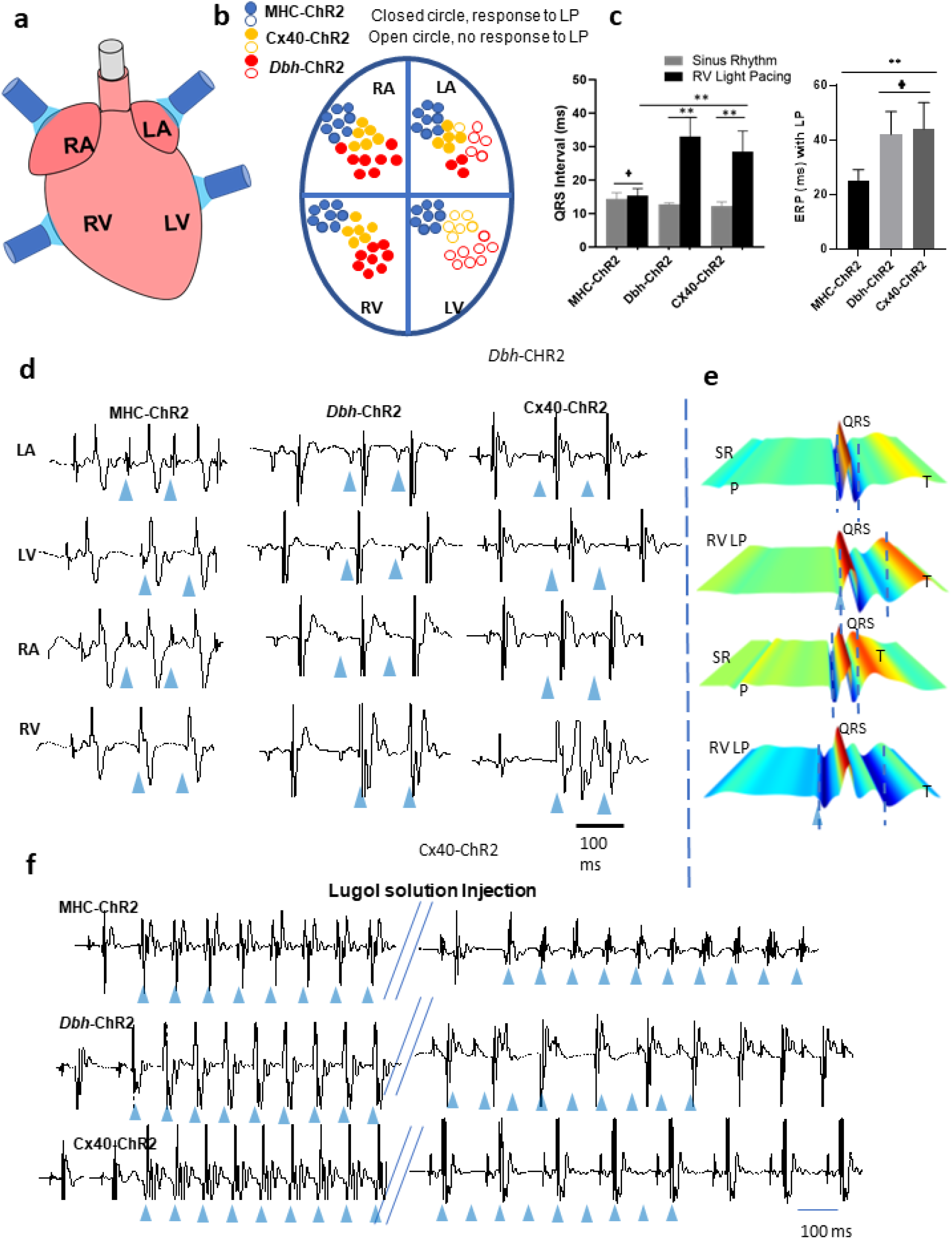
Direct optogenetic assessment of Purkinje fibers function. a. Localized light pacing in four different regions of the hearts with blue light isolated from optogenetic transgenic mouse lines expressing the fused protein tdTomato-ChR2 under the control of the promoters for MHC (MHC-ChR2), Dbh (Dbh-ChR2) and Cx40 (Cx40-ChR2). b. Summary of the electrical responsiveness to photostimulation by blue light in four different regions from MHC (MHC-ChR2), Dbh (Dbh-ChR2) and Cx40 (Cx40-ChR2) hearts assessed by ECG recordings. c. i) comparison of QRS intervals under sinus rhythm and RV epicardial photostimulation (light pacing, LP) in hearts from three optogenetic transgenic mouse lines; (QRS, MHC-ChR2: SR: 13.1 ± 2.7 ms. RVLP: 15.8 ± 1.9, n= 6; vs. *Dbh*-ChR2: SR: 12.0 ± 1.6 ms, RVLP: 32.9± 6.6 ms, n= 8 vs. Cx40-ChR2: SR: 12.3 ± 1.2. RVLP: 28.5 ± 6.2, n= 5). ii) comparison of QRS intervals under sinus rhythm and RV epicardial photostimulation (light pacing, LP) and effective refractory period (ERP) determined by RV epicardial optical programmed stimulation in hearts from three optogenetic transgenic mouse lines. (ERP: α-MyHC-ChR2, 25.2 ± 3.8 ms; n = 5; vs. *Dbh*-ChR2: 42.0 ± 8.4 ms, vs. RVLP Cx40-ChR2, 44.2 ± 8.7 ms, n=5-8 per group) * p<0.05, ** p< 0.01, + p>0.05. d. Representative ECG traces of ectopic beats originated by epicardial light stimulation of four regions from three optogenetic transgenic mouse lines, Blue arrows indicate the light pulses. e. Representative ECG waterfall plots by averaging 8-10 beats from Dbh (Dbh-ChR2) and Cx40 (Cx40-ChR2) hearts under sinus rhythm (SR) and RV epicardial photostimulation. f. Representative ECG traces of ectopic beats originated by epicardial light stimulation of the RV from three optogenetic transgenic mouse lines before and after (5 min) RV intracavital Lugol’s solution injection (n = 5-6 mice per mouse line). Blue arrows indicate the light pulses. Lugol’s solution treatment caused enlargement of the QRS complex and abolished light-induced ectopies in Dbh-ChR2 and Cx40-ChR2, but not MHC-ChR2 hearts.

We then characterised and compared the RV effective refractory periods (ERPs) determined by RV epicardial optical programmed pacing S1S2 protocol selectively photostimulation of *Dbh*-ChR2, Cx40-ChR2 and MHC-ChR2 expressing cells in these models. As summarised in Figure 7b, RV ERPs are significantly longer in both *Dbh*-ChR2 and Cx40-ChR2 hearts than those of MHC-ChR2 hearts (Figure 7c), the specific longer RV ERPs observed in *Dbh*-ChR2 and Cx40-ChR2 hearts indicate the association of *Dbh^+^-derived* CMs and Cx40^+^-derived cardiomyocytes with Purkinje fibers in RV as that reported previously in Cx40-CreERT/ChR2-tdTomato mice^28^.

Finally, to further interrogate the specific association of *Dbh^+^*-derived CMs with RV Purkinje fiber network, we applied iodine/potassium iodide solution (i.e. Lugol’s solution), a well-recognised approach used for chemical ablation of ventricular Purkinje fibers^28,29^. Regardless of the photoactivation site or intensity, the response to RV photoactivation of *Dbh*-ChR2 and Cx40-ChR2 hearts, but not MHC-ChR2 hearts, was abolished by intracavital injection of Lugol’s solution for 5 mins as exampled in Figure 7f. Lugol’s solution treatment caused prolongation of the QRS complex and abolished light-induced ectopies in *Dbh*-ChR2 and Cx40-ChR2, but not MHC-ChR2 hearts.

Taken together, the photostimulation-induced electrophysiological characteristics of *Dbh*-ChR2 and Cx40-ChR2 hearts are fundamentally similar, chemical ablation of ventricular Purkinje fibers by Lugol’s solution treatment further proves the association of the *Dbh^+^*-derived CM with Purkinje fibers, similar to the association of Cx40 derived myocytes with Purkinje fibers.

## Discussion

By utilising advanced and complementary transcriptomic analyses, genetic fate mapping, optogenetics, imaging and 3D computational image reconstruction, our study has interrogated developing and adult murine hearts, providing new insights into the development and physiology of the mammalian CCS through discovery and characterisation of novel *Dbh^+^*-CMs and *Dbh^+^*-derived CMs populations. There are several notable findings from our study:

Firstly, through scRNA-Seq with meaningful cardiomyocyte-focused quality control and utilizing a novel local and global structure-preserving dimensionality reduction technique (PHATE); we were able to discriminate rarer cardiac cell types, such as the CCS. We were left with some particular populations that were hard to classify, such as the AM-CCS. For example, AM-CCS was close in 3D PHATE space to other CCS populations (Data S2) and had expression of both atrial-like markers (*Myl7^hi^, Myl4^hi^*) and CCS-associated markers (Figure S9). We believe the AM-CCS group to likely represent either some transition zone element of the SAN, or perhaps interatrial conduction pathways. AM-CCS had low-level expression of both canonical and novel SAN markers, such as *Shox2*, *Smoc2*, *Hcn4*, and *Pcdh17*. However, AM-CCS had high expression of genes associated with the transition zone of the SAN such as *Dkk3* and *Scn5a^22^* and with other CCS components such as *Epha4*, *Nptn*, and *Gja1^22,30^*. However, it remains for future work to further characterize and validate the transcriptomic, proteomic, and physiomic signature of this population and others identified in this work.

Secondly, by undertaking integrated scRNA-Seq and spatially resolved transcriptomic analyses of the developing heart, we established a comprehensive spatiotemporally-defined transcriptional landscape of cell types populating the developing murine heart. Our spatially resolved transcriptomic Stereo-seq investigation identified major cardiac cell populations and structures through both unsupervised and supervised machine learning techniques. However, we note that there were some imperfects, but logical, predictions of cell types through supervised techniques (Figure S15). Integration of scRNA-Seq and spatially-resolved transcriptomics data remains a nascent field, and we expect future computational work to produce more accurate integration, perhaps through inclusion of prior knowledge and combinatorial scoring based on a cell’s given cassette of initially predicted labels. The utilization of the nanoscale Stereo-seq technology with high resolution (center-to-center spots distance of 500 or 715 nm) and larger capture size provides the leading resolution and tissue capture area compared to other reported spatial transcriptomic techniques, with sequencing depth and breadth comparable to many single-cell sequencing techniques, but with spatial information to further advance understanding^15^. The adoption of these novel technologies integrated into a “traditional” biological workflow at the hardware and software level helps advance the integration of bioinformatics-led discovery, reducing resource consumption in hypothesis generation and hypothesis ‘pruning’ prior to biological validation.

Thirdly, the most important, we identified novel cardiomyocyte populations, *Dbh*^+^-CMs and *Dbh^+^*-derived CMs and defined their association with the CCS in developing and mature murine hearts. Our investigation demonstrated that tdTomato reporter expression could first be detected in the neural tube at E10.5 and then in the pharyngeal arches between E10.5 to E12.5, which most likely reflects neural crest cell migration and their respective formation of sympathetic ganglia^31,32^. Our transcriptomic analyses indicate that although *Dbh^+^*-CMs represented only small percentages of the total working cardiomyocyte populations (Figure S11), the CCS appeared to be over-represented in *Dbh^+^* cardiomyocytes as percentages of their respective total cardiomyocyte lineage numbers. We are reassured that others have the *Dbh* signal present in their data from the developing and adult CCS, despite the fact we are the first to investigate further and characterize this signal. Others have utilized micro-dissection of the developing and adult CCS and observed, although not discussed, *Dbh* enrichment across the CCS, and in transcriptional programs associated with cardiac pacemaking tissue^*22,23,33*^. Our experiments demonstrate that *Dbh^+^-*CMs and *Dbh*^+^-derived CMs mainly populated the cardiac conduction system and ventricular trabecular regions throughout E14.5, and postnatal stages (Figure 5–6). Collectively, these results indicate that *Dbh*^+^-derived trabecular ventricular cardiomyocytes and *Dbh*^+^-derived ventricular PKJ myocytes likely share some common progenitor or transcriptional program, although we are unable to specify at what stage of development this progenitor or program arises or when any progeny commit to differential cell fates. We currently lack any evidence to prescribe any origin of *Dbh*^+^-CMs, or *Dbh*^+^-derived CMs, different to that of other cardiomyocytes. Further research is needed to understand the population of progenitors from which these cells arise.

Fourthly, we assayed the functional role of *Dbh*^+^-derived CMs by selectively expressing light-sensitive Channelrhodopsin-2 (*ChR2*) channels in these cells by crossing the *Dbh*^Cre^ line with the ChR2-tdTomato line. The genetic restriction of ChR2 expression allowed us to selectively interrogate the *Dbh-*derived cells as a functional part of the ventricular CCS by comparing them against a Cx40-ChR2 line, an established model for studying Purkinje fiber function^28^. In the alternative model used, the MHC promoter drives ChR2 expression to all cardiomyocyte types. Our results demonstrate that *Dbh*-ChR2 and Cx40-ChR2 hearts share remarkably similar photostimulation-induced electrophysiological characteristics and similar responses to intracavital Lugol’s solution injection. Due to the higher thickness of the LV wall, Purkinje fibers failed to be photoactivated with epicardial photostimulation, as observed in previous work^28^. Our data thus strongly support that *Dbh*^+^-CMs and/or *Dbh*^+^-derived CMs form a functional part of the ventricular CCS.

Finally, our findings suggest a potential relationship between sympathetic innervation and *Dbh^+^*-derived CMs, during CCS formation, or vice versa (Figure S18). Figure 8 summarise our hypothetical relationships of sympathetic innervation and *Dbh*^+^-derived CMs and *Dbh*^+^ CMs contributed CCS formation in developing and mature hearts. It is known that sympathetic innervation occurs concurrently with the development of the cardiovascular system^34^. The importance of the sympathetic nervous system, or some other catecholaminergic source, is supported by the evidence of embryonic death with cardiac developmental defects due to two most important catecholamine synthetic enzymes, *Th*-deficient^8^ and *Dbh*-deficient mice^9,10^. The relationship between sympathetic innervation and cardiac pathology in the adult^35^ and the perseverance of *Dbh^+^*-CMs into adult hearts, raises important pre-clinical questions about any putative interaction between the sympathetic innervation of the heart and *Dbh^+^*-CMs or *Dbh^+^*-derived CMs, which will be the future focus of our study.

**Figure 8.**
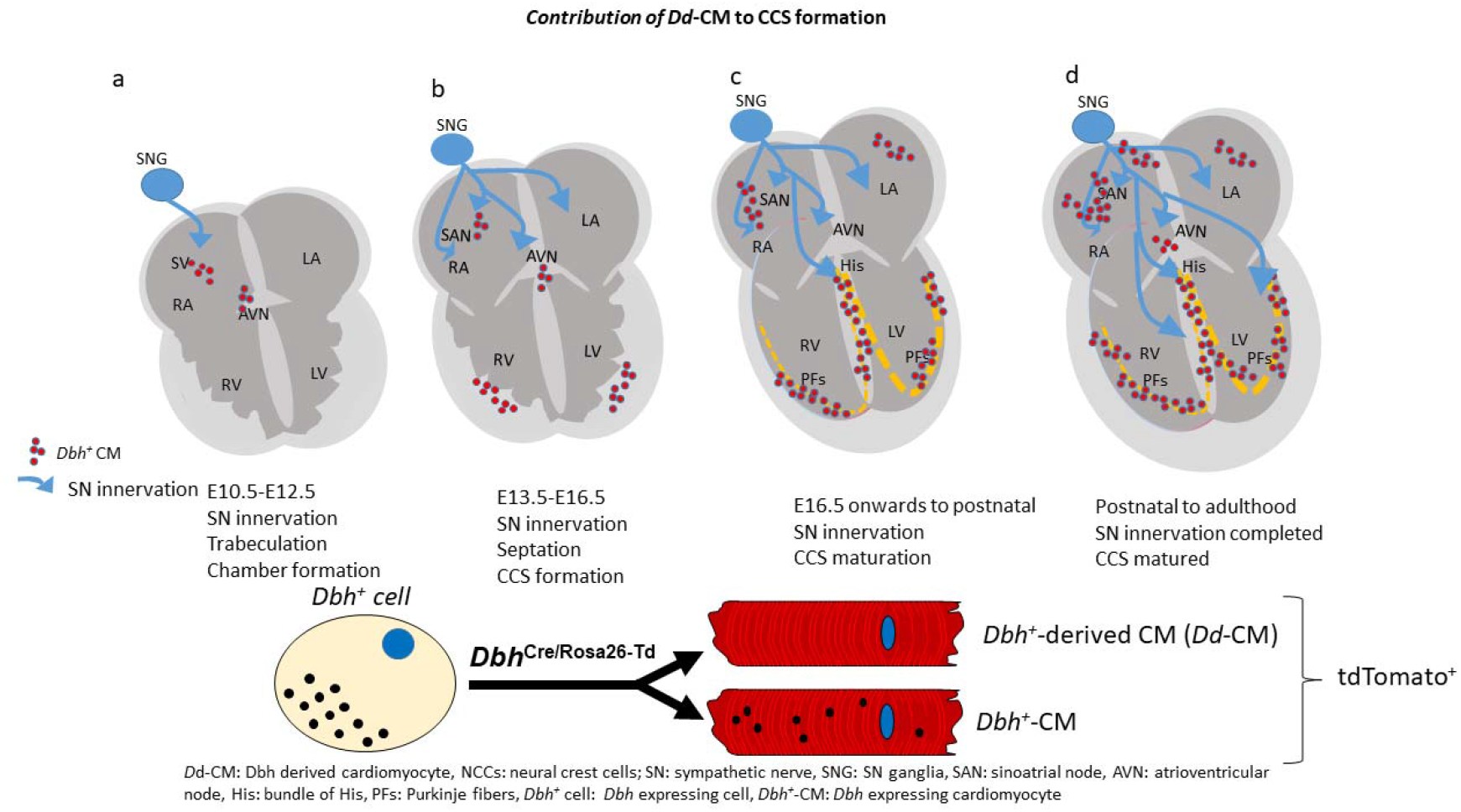
Proposed relationships of SN innervation and Dbh^+^-CM contributed CCS formation. a. Dbh^+^-CM-associated CCS formation occurred concurrently with sympathetic innervation from E12.5 after neural crest cells migration occurred in SN ganglia and heart region from E10.5. Such process is in parallel to the processes of ventricular trabeculation and chamber formation. b. From E14.5 onwards, the Th positive nerve fibres were clearly identified and they extended to the AV junction, His-Bundle and part of the ventricular myocardium. Such process occurred concurrently with Dbh^+^-CM-associated CCS formation. c. E16.5 onwards to postnatal, SN innervation continues and CCS is maturing. d. from postnatal to adulthood, SN innervation and maturation of CCS are completed.

In conclusion, we have studied the development and function of the murine cardiac conduction system by applying a sophisticated approach integrating single-cell RNA sequencing, high resolution spatially-resolved transcriptomics, unsupervised and supervised machine learning techniques, cell fate mapping, multi-modal imaging, and optogenetics. Our identification of *Dbh*^+^-CMs and *Dbh*^+^-derived CMs provide new insights into the mid-to-late development of the mammalian CCS through this previously unreported cardiac catecholaminergic cardiomyocyte population.

## Acknowledgements

We thank Peter Somogyi, FRS, FMedSci, PhD, FRS (Department of Pharmacology, University of Oxford) for help with imaging experiments and Wenyuan Zhang (Department of Zoology, University of Oxford) for help in preliminary bioinformatic analysis. We thank Prof. Gillian Douglas (Radcliffe Department of Medicine, University of Oxford) for providing MHC-Cre mouse line. We thank Niloufer G Irani and colleagues in Micron Advanced Bioimaging Unit at Department of Biochemistry (University of Oxford) for providing support and assistance for imaging work. We thank production team of China National GeneBank at Shenzhen for their assistance in providing Genebank resources. We thank Advanced Research Computing (ARC) service at University of Oxford for providing High Performance Computing (HPC) resource for the data process and analysis. This work was supported by the National Natural Science Foundation of China [No. 81670310, X.T, No. 31300948, X.T, No. 81700308 X.O & No, 31871181, X.O, M.L.], Outstanding Youth Foundation of Sichuan Province, China [No. 20JDJQ0047, X.T.], the British Heart Foundation, UK [PG/11/59/29004 (M.L.) PG/14/80/31106 (M.L.), PG/16/67/32340 (M.L.), PG/21/10512 (M.L.), FS/PhD/20/29053 (M.L., H.Z.), Medical Research Council, UK G100647 (M.L.), Wellcome Strategic Awards 091911/B/10/Z and 107457/Z/15/Z) (Micron Advanced Bioimaging Unit). We thank Dr. Tania Zaglia for scientific advice (Department of Biomedical Sciences, University of Padova).

## Author Contributions

M.L., W.S., X.T. designed the study, planned the experiments and supervised the study. M.L., W.S., T.Y.S., A.G.R. and X.T. interpreted the results and wrote and finalized the manuscript. T.Y.S. conducted most immunohistochemistry, in situ hybridization experiments, imaging, and lineage analysis. A.G.R. analysed scRNA-Seq and Stereo-seq data, performed cell typing and prepared scRNA-Seq and Stereo-seq Figures. H.R., Z.P., K.K., X.O., T.C. and X.F. processed embryos and hearts for single-cell library preparation for scRNA-Seq. H.R., Z.P., T.C. processed embryos and hearts for Stereo-seq experiment. Y.A, X.G. jointly with A.G.R. prepared Stereo-seq data. T.S. conducted preliminary scRNA-Seq data analysis. Y.A, A.C, Z. S. provided support for Stereo-seq and data analysis. T.Y.S., A.G.R. M.L, P.L, Z.L. and Z.P. conducted opogenetic electrophysiological experiments, data analysis and prepared the figure. Y.L., YY.L. Performed embryonic tissue sectioning, immunohistochemical staining and the imaging. W.H., H.Z. performed three-dimensional image computational reconstruction. H.R., Z.P. T.Y.S. and Y.H. managed mouse lines. Z.X. performed imaging analysis. N.S. provided support for imaging and helped with manuscript preparation, LM provided Cx40cre and Cx40GFP mouse lines and helped with data interpretation. All authors read and approved the final manuscript.

Data and materials availability: The *Dbh*^Cre^ mouse lines is available from M.L. and X.T. under a material transfer agreement with the University of Oxford and Southwest Medical University. Interactive versions of certain data are included with paper.

## Methods

### EXPERIMENTAL MODEL AND SUBJECT DETAILS

No statistical methods were used to predetermine sample size. The experiments were not randomized and investigators were not blinded to allocation during experiments and outcome assessment.

#### Animals and ethical approval

All animal experiments were performed on mice neonatal or adult mice in accordance with the United Kingdom Animals (Scientific Procedures) Act 1986 and were approved by the University of Oxford Pharmacology ethical committee (approval ref. PPL: PP8557407) or Animal Care and Use Committee of the Southwest Medical University, Sichuan (China) (No: 20160930) in conformity with the national guidelines under which the institution operates. All mice including mutant mice and wild-type (WT) littermates used in this study were maintained in pathogen-free facilities at the University of Oxford or Southwestern Medical University. Mice were given *ad libitum* access to food and water.

#### Generation of mouse models

The *Dbh*^Cre^ mouse model is generated by using CRISPR/Cas9 technique that the 2A-iCre-WPRE-polyA expression box is knocked in at the stop codon site of the *Dbh* gene through homologous recombination technology. The brief processes include: 1) Cas9 mRNA and gRNA are obtained by in vitro transcription; 2) homologous recombination vector is constructed by In-Fusion cloning method, which contains a 3.0 kb 5’homology arm, 2A-iCre-WPRE-polyA segment and 3.0 kb 3’homology arm; 3) Cas9 mRNA, gRNA and donor vectorwere microinjected into the fertilized eggs of C57BL/6J mice to obtain F0 generation mice. The positive F0 generation mice identified by PCR amplification and sequencing were mated with C57BL/6J mice to obtain F1 generation mice.

*Dbh*^Cre^/Rosa26-tdTomato mice were then obtained by crossing *Dbh*^Cre^ mice with B6.Cg Gt(ROSA)26Sortm9(CAG-tdTomato)Hze/J strain (Stock No. 007909, Jackson Labs) for lineage tracing study. *Dbh*^Cre^/ChR2-tdTomato mice were obtained by crossing *Dbh*^Cre^ mice with B6.Cg-Gt (ROSA) 26Sortm27.1(CAG-COP4*H134R/tdTomato) Hze/J strain (Stock No. 012567, Jackson Labs). The resulting offspring have the STOP cassette deleted in cardiomyocytes, driving the expression of ChR2 [hChR2 (H134R)-tdTomato fusion protein].

*Dbh*^CreERT^ mouse model is generated by using CRISPR/Cas9 technique that the 2A-iCre-WPRE-polyA expression box is knocked in at the stop codon site of the *Dbh* gene through homologous recombination technology. The *Dbh*^CreERT^/Rosa26-tdTomato mice was generated by crossing *Dbh*^CreERT^ mice with C57BL/6JSmoc-Gt(ROSA)26Sorem(CAG-LSL-tdTomato)1Smocmice from Shanghai Model Organisms Center, Inc. (Shanghai, China).

*Dbh*^CFP^ mouse model is generated by using CRISPR/Cas9 technique that the IRES-CFP-5×Flag-WPRE-polyA expression box is knocked in at the stop codon site of the *Dbh* gene through homologous recombination technology.

*Dbh*^Cre^/ChR2-tdTomato/Cx40-eGFP mice were obtained by crossed *Dbh*^Cre^/ChR2-tdTomato heterozygous positive mice with a Cx40-eGFP mouse line ^36^

For in vivo optogenetic study, transgenic adult male mice (8 weeks), with a genetic background C57B6J, expressing cre-recombinase under the control of either the MHC (B6.FVB-Tg(Myh6-cre)2182Mds/J, Stock No. 011038, Jackson Labs) or Cx40 promoter were bred with B6.Cg-Gt (ROSA) 26Sortm27.1(CAG-COP4*H134R/tdTomato) Hze/J strain (Stock No. 012567, Jackson Labs). The resulting offspring had the STOP cassette deleted in the heart, resulting in cardiomyocyte (MHC-ChR2) or Purkinje fiber (Cx40-ChR2) expression of the hChR2(H134R)-tdTomato fusion protein. MHC-Cre mouse line was provided by Prof. Gillian Douglas (Radcliffe Department of Medicine, University of Oxford). Cx40-CreERT was imported from Prof. Lucile Miquerol’s Lab (Institut de Biologie du Développement de Marseille-Luminy).

##### Tamoxifen Treatment

Adult (week 8) mice (Cx40-ChR2 and *Dbh*^CreERT^/Rosa26-tdTomato transgenic mouse model) were injected with Tamoxifen (Sigma) (150 mg/kg, gavage) for 5 consecutive days. After one-week post-induction recovery, mice underwent subsequent experiment.

### METHOD DETAILS

#### Experimental design

##### Replication

Each of the developmental embryos or heart tissues were collected at unique time points, i.e. 8.5, 10.5, 12.5, 14.5 and P3. Samples are biological replicates to compared gene expression detected by SrT and scRNA-Seq. Consecutive tissue sections from the same heart tissue were considered technical replicates in the SrT (Figure 1a, Figure 2a) and RNAscope and immunohistology (Figure 3c; Figure 4–6, Figures S1, S12) experiments. However, it is important to notice that consecutive sections are highly similar but not identical.

##### Embryo and heart dissection

The whole embryos or hearts were dissected from developmental stages at E10.5, E11.5, E12.5, E13.5, E14.5, E16.5 and neonatal stage. Embryos or hearts were dissected in cold PBS (Life Technologies, CAT# 14190250), de-yolked and placed in PBS on ice until dissociation. The tissues were fixed with cold 4% paraformaldehyde (PFA) in 1× PBS for 15-20 minutes. After short fixation, the embryos and heart regions were either stored in 70% ethanol at 4°C for future use or embedded in optimal cutting temperature (OCT) and frozen in cold isopentane cooled by dry ice and stored at −80°C. Coronal cryosections (10 μm) were collected on a cryostat (Leica), mounted onto superfrozen glass slides (VWR) and stored at −80°C. For adult heart sample preparation, the mice were killed by cervical dislocation, the heart was dissected and perfused PBS solution by through aorta. The heart was then fixed with cold 4% paraformaldehyde (PFA) in 1× PBS for no more than 30 min and moved into the 10% sucrose for 1 hour, subsequently transferred to 20% sucrose for 1 hour, and then cryoprotected into 30% sucrose for overnight at 4°C. The heart was then embedded in OCT and frozen on isopentane-dry ice slurry and stored at −80°C. Coronal cryosections (10 μm) were collected on a cryostat (Leica), mounted onto super-frost glass slides (VWR) and stored at −80°C.

##### 10× Genomics Chromium Single-cell RNA sequencing

The single cells were either isolated from embryos at E8.5 and E10.5 stages, or embryonic hearts at E12.5, E14.5 and E16.5 stages, and neonatal mouse cardiomyocytes (from P3) using the standard enzymatic method described previously^37^. Single-cell suspension samples were quality tested, quality control standards: cell activity > 80%, cell concentration 700-1200 cells / sL, cell diameter 5-40 m, the total number of cells up to 500,000. Single cells were prepared following the protocol from 10× Genomics, Inc (Pleasanton, CA). The protoplast suspension was loaded into Chromium microfluidic chips with 30 (v3) chemistry and barcoded with a 10× Chromium Controller (10× Genomics). RNA from the barcoded cells was subsequently reverse-transcribed and sequencing libraries constructed with reagents from a Chromium Single Cell 30 v3 reagent kit (10× Genomics) according to the manufacturer’s instructions. Sequencing was performed with Illumina HiSeq according to the manufacturer’s instructions (Illumina). FastQC was used to perform basic statistics on the quality of the raw reads.

Raw reads were demultiplexed and mapped to the reference genome by 10× Genomics Cell Ranger pipeline(https://support.10xgenomics.com/single-cell-geneexpression/) using default parameters. All downstream single-cell analyses were performed using Cell Ranger and Seurat^38^ unless mentioned specifically. In brief, for each gene and each cell barcode (filtered by CellRanger^38^), unique molecule identifiers were counted to construct digital expression matrices in detail: cellranger count takes FASTQ files performs alignment, filtering, barcode counting, and UMI counting. It uses the Chromium cellular barcodes to generate feature barcode matrices.

##### Initial cell typing in single-cell RNA sequencing data

To ensure robust and reliable transcriptomic signal-to-noise ratios, without impairing sensitivity to small signals, we filtered out all cells with unique RNA counts (nUMI) <300, or distinct genes (nFeatures) <270 to remove under-sampled cells and simple cells such as erythrocytes. The ratio of mitochondrial transcripts to nuclear genome-derived transcripts is often used as a metric of cell stress or quality in single-cell RNA sequencing. 5% is typically used as a ceiling, but this is not supported across cell types and can fail to identify damaged cells, particularly cardiomyocytes, which can reach ~30%^24^ and exclude particular cardiomyocyte populations^39^. Given the changing nature of mitochondrial biogenesis across the embryonic to postnatal mouse heart^40^, we utilized a dynamically changing filter for mitochondrial transcript ratios: E8.5: 5%, E10.5: 5%, E12.5: 7.5%, E14.5: 10%, E16.5: 15%, P3: 20%. We normalized cell libraries through SCTransform, which accounts for preservation of differential variation between highly variant and lowly variant genes^41^. We utilized Uniform Manifold Approximation and Projection (UMAP), following principal component analysis (PCA) and PCA dimension selection, to enable human-interpretable visualization of the transcriptomic space through dimensionality reduction. We visually examined the data for differences between batches of cells collected at each stage. Only P3 had significant batch differences. There are extensive suggested solutions for “correcting” batch differences^42^. However, from a statistical fundamentals perspective both a priori and empirically, such methods have been shown to produce aberrant downstream results^43^ and so were not used here. We visualized UMAP in 3D and used Louvain clustering, and labelled clusters based on expression profiles. Further sub-clustering was performed as necessary, and a small number of cells were manually assigned where appropriate. Above was done using Seurat ^44^ functions unless specified.

##### Identification of Dbh^+^ cardiomyocytes

To improve the discrimination of cardiomyocyte-lineage specific cell types in the Transcriptomic space, we isolated the cardiomyocyte-lineage cell types identified transcriptomically: Cardiac Progenitors, Early Cardiomyocytes, Immature Atrial Cardiomyocytes, Immature Ventricular Cardiomyocytes, Cardiac Conduction System, ECM Atrial Cardiomyocytes, ECM Ventricular Cardiomyocytes, Atrial Cardiomyocytes, Ventricular Cardiomyocytes, and Myh6+ Ventricular Cardiomyocytes. We then re-normalised their raw nUMI counts, using SCTransform, and used PHATE^21^ for dimensionality reduction and clustering. PHATE preserves global and local structure in the transcriptomic space of genes observed across cells: intuitive visualisation of high-dimensional gene expression data must balance within cell type (local) and between cell-type (global) transcriptional differences: local discrimination of transcriptional differences within cell types may be obscured by preservation of global between-cell type differences. PHATE enables intuitive interpretation of datasets, particularly so for “lineage” type datasets in our experience. Cell types were assigned by comparison with literature expression profiles. We used the parallel implementation of Seurat’s FindMarkers[https://github.com/vertesy/Seurat.multicore], implemented with min.pct =0.2, logfc.threshold=0.5, and test.use=“bimod” to compare differential gene expression between cell types. In selecting *Dbh*^+^ cells from this lineage, we isolated all cells from the cardiomyocyte lineage with a raw nUMI for *Dbh* > 0. We then re-normalised these cells, using SCTransform, and performed PHATE dimensionality reduction. We used cell labels identified from the cardiomyocyte-lineage. To identify whether *Dbh* expression (defined as UMI>0 or UMI=0), was independent of cell type, we performed a two-tailed Chi Squared test for independence, using only cardiomyocyte lineage cell types composed primarily of cells beyond E10.5, as before this *Dbh* expression was extremely limited in cell types. Therefore, we excluded Heart fields, Endocardial-gene rich cardiomyocytes, Developing cardiomyocytes, Misc., and Primary heart tube from testing.

##### Stereo-seq spatial transcriptomics

###### Tissue Collection

The hearts at different stages (E12.5, E14.5, P3, and adult (P56)) were dissected from the wild type or *Dbh*^Cre^/Rosa26-tdTomato mice and washed with PBS quickly. For adult heart dissection, the mice were killed by schedule 1, the heart was dissected and perfused PBS solution by through aorta and rings collected after embedded in Tissue-Teck OCT (Sakura, 4583). The heart samples were embedded in OCT. The OCT and frozen on isopentane-dry ice slurry and stored at −80°C. Cryosectioning was performed with Leica CM1950 cryostat and the sections were cut with 10 μm thickness. Then the tissue sections were adhered to Stereo-seq chips immediately for the following experiments. Adjacent sections would be adhered to a glass slide to check tdTomato fluorescence.

###### Stereo-seq experiment

Stereo-seq library preparation and sequencing were previously described^15^. After tissue sections were adhered, Stereo-seq chips were incubated at 37 L slide dryer for 3 min. Then the chips were immersed in prechilled methanol for 30 min. After fixation, tissue sections were stained with ssDNA reagent (Invitrogen, Q10212) for 5 min, and then washed using 0.1× SSC buffer (Ambion, AM9770) supplemented by 0.05 U/μl RNase inhibitor (0.1× SSC+RI). The ssDNA images were captured with Ti-7 Nikon Eclipse microscope using FITC channel. After imaging, tissue permeabilization was performed with 0.1% pepsin (Sigma, P7000) in 0.01 M HCl buffer (pH = 2) at 37 L incubator for optimal time, which is 6 min for E12.5, E14.5 and P3 heart sections, and 18 min for adult heart sections. After washing with 0.1× SSC+RI, Reverse transcription was performed with SuperScript II (Invitrogen, 18064014, 10 U/μL reverse transcriptase, 1 mM dNTPs, 1 M betaine solution PCR reagent, 7.5 mM MgCl2, 5 mM DTT, 2 U/μL RNase inhibitor, 2.5 μM Stereo-TSO and 1× First-Strand buffer) at 42 L incubator for overnight. After washing twice with 0.1× SSC buffer, tissues on the chip surface were digested with Tissue Removal Buffer (10 mM Tris-HCl, 25 mM EDTA, 100 mM NaCl, 0.5% SDS) at 37 L incubator for 30 min. After washing with 0.1× SSC buffer for two times, the chips were treated with Exonuclease I (NEB, M0293L) at 37 L incubator for 1 h. After washing once with 0.1x SSC buffer, cDNA was amplified using KAPA HiFi HotStart Ready Mix (Roche, KK2602) with 0.8 μM cDNA-PCR primer. For library preparation, cDNA products were further fragmented using in-house Tn5 transposase, amplified with KAPA HiFi HotStart Ready Mix, and purified using AMPure XP beads (Beckman Coulter, A63882). The purified PCR products were used to make DNBs (DNA nanoballs) which were sequenced with a MGI DNBSEQ-Tx sequencer.

###### Stereo-seq data analysis

Raw data processing method was carried as previously described^23^. Briefly, CID and MID were extracted from the read 1, and the cDNA sequences were extracted from the read 2 in the FASTQ files from MGI DNBSEQ Tx sequencer. CID mapping allows 1 base mismatch. MID reads with N bases or more than two low quality score bases (score < 10) were removed. The remaining DNA reads were aligned using STAR against the reference mouse genome (mm10) supplemented with WPRE sequences obtained from transgenic plasmid. To each gene of each CID, reads with same MIDs were only kept one for counting. After annotation and count, gene expression matrices with CID were generated.

###### Unsupervised clustering

Gene expression matrices were divided into bins with 20 × 20 DNBs for E12.5, E14,5, and P3 hearts, and 50 × 50 DNBs for adult heart. Density plot was used to visualize the gene count distribution in each bin, and the minor peak containing bins with low gene counts were excluded. Genes expressed in fewer than 5 bins were also discarded. After obtaining the final gene expression matrices of bin20 or bin50, data for all slices including three E12.5 slices, six E14.5 slices, and six P3 slices were merged to do the unsupervised clustering using Seurat. Briefly, SC Transform function was used to normalize the data and variable genes were identified. Then Run PCA was used for dimension reduction and run UMAP was used to indicate the two-dimensional projection with dims= 30. FindNeighbors and FindClusters were further used to identify clusters. Find All Markers was used to identify the differentially expressed genes for each group with the parameters: min.pct = 0.1, logfc.threshold = 0.25. Each cluster was annotated using several known cell type-specific marker genes. We assigned cluster labels using gene expression profiles and bin location.

###### Integration of single cell and spatial transcriptomic data

To elucidate probabilistic spatial distributions of our cell types identified in our single-cell RNA sequencing data; we combined cells identified from the cardiomyocyte lineage with those identified in the wider dataset, to give a representative transcriptional reference library of potential cell types in the developing murine heart from E12.5 to P3. We combined: Atrial Cardiomyocytes, Ventricular Cardiomyocytes, Immature Atrial Cardiomyocytes, Immature Ventricular Cardiomyocytes, Immature Trabecular Ventricular Cardiomyocytes, Trabecular Ventricular Cardiomyocytes, Sinoatrial Node, Atrioventricular Node, Purkinje Fibres, Fibroblast-like, Smooth Muscle-like, Endothelium, Epicardium, Endocardium, Neural Crest, Haematopoetic Precursors, Immune Cells, and Platelets. We re-normalised the raw nUMI of this library using SCTransform. To identify specific and robust marker genes for each cell type, we used SMASH, and in brief, kept the top 50 genes (Table S2) that contributed most to the classification of each cell type through identifying gene-wise Gini importance from the trained supervised ensemble machine learning classifier algorithm.

###### Immunofluorescence

The frozen sections were post fixed in 4% PFA fixation for 15 min at room temperature. The sections were washed in PBS for three times and 5 minutes for each wash, followed by permeabilization in 0.1% TritonX-100/PBS for 30 min. After three washes in PBS, the tissue sections were blocked by 0.1% Tween20, 10% Normal serum for 1 hour at room temperature. The sections were then incubated in the primary antibody for overnight at 4 °C. The slides were put in RT for 1 hour for recovery. After three washes, the sections were incubated in the secondary antibody for 2 hours. Finally, the mounting medium with DAPI were applied and nailed polish was used to seal the slides with coverslip.

###### Multiplex nucleic acid in situ hybridization (RNAscope)

Each *RNAscope* experiment was replicated at least twice for identifying spatial expression of genes. The fixed frozen tissue slides were thawed at room temperature for 15 min and baked at 60°C for 30 min. Afterwards, slides were post-fixed for 15 min in chilled 4% PFA, and serially dehydrated through 50%, 70%, 100%, and 100% ethanol for 5 minutes each. After dry for 5 minutes, the sections were applied the RNAscope hydrogen peroxide for 10 minutes. 1× Target retrieval reagent covered the sections for 5 minutes at 95°C followed by two washes in distilled water. Tissue sections were permeabilized the protease digestion (ACD Protease III) for 20 minutes. The probes in C1 and C3 channels were labelled by using Opal 520, and 690 fluorophores (Akoya Biosciences, diluted 1:200) respectively. The C4 probe was developed using TSA biotin (TSA Plus Biotin Kit, Perkin Elmer, 1:500). Nuclei staining was performed with DAPI. Imaging was performed on a Zeiss Black software in the Zeiss confocal LSM780 microscope using 10× or 20× air-immersion objectives or 63 water-immersion objective. The excitation (Ex) laser and emission (Em) filter wavelengths were: DAPI (375nm; 435-480 nm), Opal 520(488 nm; 500-550 nm), Opal 690 (676 nm; 690-700 nm).

###### 3D reconstruction of the heart

Three-dimensional image reconstruction was conducted to present the spatial architecture and morphology of cell distributions at the microscopic level (around μm) for the whole heart. The method was based on a modified procedure of our previous approach ^45^ to proceed fluorescence images of multiple coronal slices of adult and neonatal murine cardiac tissue. To establish a 3D atlas of whole heart with details of spatial distribution of positive and negative labelled cardiac cells, each two-dimensional (2D) fluorescence image was first pre-processed for registration, segmentation, and enhancement before final fusion for 3D reconstruction of the adult and neonatal hearts. Input: Raw 2D fluorescence images were used after spatial resolution between the two groups were unified (the resolution of the adult mouse cardiac images was downsized as they are about 20-fold of those of the neonate ones). Registration: Registration involves two steps: the contour extraction and the affine transformation. To account for the distinctively dispersed cell spatial distribution in the raw images, the Flood Fill algorithm was used to calculate numerous limited bounded regions, which corresponded to the position of holes. Once the positioning process was completed, a bitwise NOT inverted operation was performed to generate a new matrix, which was then logically computed with the raw image matrix to achieve the hole filling target. Finally, the Sobel operator method was utilized to construct a base-image emphasizing edge, followed which the new edge was then added to the former layer of each image to execute the affine transformation (rotation and translation) process. Contrast Enhancement: By modifying relative parameters gain (γ) and bias (β) in the following equation, we were able to optimize image features and boost color contrast using a common linear transformation method: gx,y=yfx,y+/3. In this equation, fx,y is the input pixel value, gx,y is the processed pixel value, the gain parameter y is used to increase contrast ratio, and the bias parameter /3 is used to control brightness.

###### Channel Split

Once the multiple RGB images were enhanced, channel split measures were taken to separate them into red, green, and blue channels to implement the pixel aggregation operation. The specific procedures are bellowing: On the one hand, repeated the Hole Filling method (see the registration method) based on the blue channel and reassign all pixel values to 40 to create the basic contour. On the other hand, extracted red and blue channels and rewrite these pixel values separately to 180 and 10, and then clear the green channel values.

###### Image Merge

To construct the processed image and carry out a series of canonical images, we further merge these images with the same weight mechanism. Each image has a light grey base color for the heart profile and a bright color for a positive fluorescence cell. Frequency Analysis: With the purpose of quantify the pretreatment process, in this study we implemented the frequency analysis before 3D model building.

###### 3D Reconstruction

Finally, all images were written to the same data file of VTK format (http://www.vtk.org/) to expedite hand visualization based on Paraview software (http://www.paraview.org/).

###### Optogenetic Electrophysiology

Optical stimulation of ChR2 light-sensitive channel. Whole hearts were paced through the activation of ChR2 light-sensitive channels. This was achieved by the delivery of 470 nm blue light pulses (5–10 ms pulse width) generated by OptoFlash (Cairn Research). Pulses were triggered by a 1401 digitiser and Spike 2 software (Cambridge Electronic Design). Approximate blue light intensity was measured with a 818-ST2 Wand Detector connected to a 843R Power meter (both Newport Corporation, CA, USA) and expressed normalised for the area being illuminated through simulating the average light intensity reached to the surface of the tissue by mimicking the distance of fibre-heart.

To monitor cardiac rhythms, we carried out ex vivo ECG analysis on langendorff perfused hearts. ECG parameters including RR interval, PR interval, QRS and QT durations were measured. Langendorff-perfused ex vivo hearts from *Dbh*^Cre^/ChR2 mice were subjected to programmed light stimulation (PLS) while ECGs were measured. Two protocols were carried out as follows: (a) continuous pacing protocol: Stimuli were delivered continuously with a constant frequency between 8-10 Hz. (b) S1S2 pacing protocol: a pacing train of eight stimuli (S1) was delivered at a basic cycle length of 100ms, with a single (S2) premature extra stimulus introduced at progressively shorter intervals until the refractory period was reached.

Measurement of the Ventricular ERP. The ventricular ERP (VERP) of both the working cardiomyocytes and Purkinje fibers was evaluated in *Dbh*^Cre^-ChR2, MHC-Cre/ChR2-tdTomato and Cx40-CreERT/ChR2-tdTomato, respectively, by using S1S2 pacing protocol. The epicardial surface of the RV was photostimulated by a train of 10 light pulses (5-10 ms), at a frequency of 10 Hz (cycle length, 100 ms) (S1), followed by a premature optical stimulus (S2) at a progressively shorter coupling interval until the S2 failed to trigger ectopic beats at the VERP.

Pharmacological Purkinje Fiber Ablation. We carefully injected 500-1000 μl of Lugol’s solution [5% (wt/v) iodine and 10% (wt/v) potassium iodide, in distilled water] into the RV cavity by using a Hamilton syringe supporting a 34G needle followed by the continuous pacing protocol with RV epicardial light pacing.

---

Supplementary Figure 1. **Summary of single-cell RNA sequencing computational workflow**

a i) Isolated cells were sequenced by 10 x Genomics Illumina HiSeq, and then underwent initial read-wise quality control and UMI barcode matrix construction. Cells then underwent cell-wise quality control, with cells kept only if they met all of the following characteristics described to the right. We normalised from raw RNA counts through SCTtransform, to account for variance in library size. We performed principal component analysis to identify and select only those components contributing to the majority of computational and biological variation for downstream analyses. We performed dimensionality reduction using UMAP, to reduce the dimensionality of the dataset around 10,000x, to enable more intuitive engagement. We performed Louvain clustering to identify sufficiently similar cells within our data to produce clusters. We labelled clusters with biological identities based on their gene expression, using both normalised count data and their Pearson’s residuals from respective average gene expression. Any ambiguous clusters were analysed further, including looking at wider gene expression, subclustering, and in a small number of cases assigning biological identities directly to cells. viii) We visualised the data in its completed format as presented in the paper.

b We selected only those cell types involved in the cardiomyocyte lineage, from mesoderm through to P3 cardiomyocytes (See methods), and subset their raw count data from the original raw matrix. We normalised from raw RNA counts through SCTtransform, to account for variance in library size. We performed principal component analysis to identify and select only those components contributing to the majority of computational and biological variation for downstream analyses. We performed dimensionality reduction using PHATE, to reduce the dimensionality of the dataset around 10,000x, to enable more intuitive engagement. We performed k-means clustering on the PHATE operator, similar to spectral clustering, to identify sufficiently similar cells within our data to produce clusters. We labelled clusters with biological identities based on their gene expression, using both normalised count data and their Pearson’s residuals from respective average gene expression. Any ambiguous clusters were analysed further, including looking at wider gene expression. We visualised the data in its completed format as presented in the paper.

c We selected all cells from the cardiomyocyte lineage that had raw UMI counts of *Dbh>0*. We visualised the data in its completed format as presented in the paper.

Supplementary Figure 2. **Quality control metrics for single-cell data**.

a) A UMAP plot of all initial cells post-quality control, coloured by Log_10_(nUMI) and a violin plot demonstrating the same across different stages.

b) A UMAP plot of all initial cells post-quality control, coloured by nGene and a violin plot demonstrating the same across different stages.

c) A UMAP plot of all initial cells post-quality control, coloured by the ratio of mitochondrial nUMI to total nUMI and a violin plot demonstrating the same across different stages.

Supplementary Figure 3. **scRNAseq identified 4 major groups of cell types across the developing mouse heart**.

a) i) UMAP plot of all initial cells post-quality control, coloured by cell type identified after unsupervised clustering. This dataset includes E8.5 and E10.5 whole embryos, and whole isolated hearts from E12.5, E14.5, E16.5, and P3 mice. Corresponding cell types are labelled below.

ii) UMAP plot of all initial cells post-quality control, coloured by stage of tissue isolation.

iii) UMAP plot of all initial cells post-quality control, coloured by predicted Cell Cycle stage identified using Seurat.

Supplementary Figure 4. **Key marker genes for each population initially identified within our scRNA-Seq dataset**.

Each cell type corresponding to Supplementary Figure 3 cell types, is identified in bold, with some respective key exemplary key markers underneath. Gene^+^ indicates a gene is expressed above the population level in this group. Gene^hi^ indicates that a gene is highly expressed above the population level in this group. Gene^lo^ indicates a gene is expressed below the level expected for it to alter the classification of a particular group and is typically in reference to sub-population expression levels rather than the whole dataset.

Supplementary Figure 5a. **Expression of example marker genes across different cell populations identified in whole dataset**.

Each panel displays a UMAP plot of the whole dataset with cells coloured by the expression of the specific gene identified at the top of the plot. The expression is given in post-SCT normalized counts. The cell type that the individual genes are representative of are indicated above each plot, with corresponding colours and numbers as in Supplementary Figure 3.

Supplementary Figure 5b. **Expression of example marker genes across different cell populations identified in whole dataset**.

Each panel displays a UMAP plot of the whole dataset with cells coloured by the expression of the specific gene identified at the top of the plot. The expression is given in post-SCT normalized counts. The cell type that the individual genes are representative of are indicated above each plot, with corresponding colours and numbers as in Supplementary Figure 3.

Supplementary Figure 5c. **Expression of example marker genes across different cell populations identified in whole dataset**.

Each panel displays a UMAP plot of the whole dataset with cells coloured by the expression of the specific gene identified at the top of the plot. The expression is given in post-SCT normalized counts. The cell type that the individual genes are representative of are indicated above each plot, with corresponding colours and numbers as in Supplementary Figure 3.

Supplementary Figure 5d. **Expression of example marker genes across different cell populations identified in whole dataset**.

Each panel displays a UMAP plot of the whole dataset with cells coloured by the expression of the specific gene identified at the top of the plot. The expression is given in post-SCT normalized counts. The cell type that the individual genes are representative of are indicated above each plot, with corresponding colours and numbers as in Supplementary Figure 3.

Supplementary Figure 6. **Cell types display a range of associated biologically defining marker genes in our single-cell RNA sequencing analyses**.

a) Cell types identified in Supplementary Figure 3 from initial analysis of all cells post-quality control. The annotation on the right describes the relative proportions of each cluster in terms of developmental stage. Values for expression are normalised by row and then by column for visualization purposes.

b) Cell types identified in Figure 1b from cardiomyocyte lineage. The annotation on the right describes the relative proportions of each cluster in terms of developmental stage. Values for expression are normalised by row and then by column for visualisation purposes.

c) i) A 2D plot of *Tnnt2* expression across *Dbh^+^* cardiomyocytes. ii) 2D plots of various CCS markers across *Dbh^+^* cardiomyocytes.

Supplementary Figure 7. **Key marker genes for each population initially identified within our scRNA-Seq dataset cardiomyocyte lineage**.

Each cell type corresponding to Supplementary Figure 3 cell types, is identified in bold, with some respective key exemplary key markers underneath. Gene^+^ indicates a gene is expressed above the population level in this group. Gene^hi^ indicates that a gene is highly expressed above the population level in this group. Gene^lo^ indicates a gene is expressed below the level expected for it to alter the classification of a particular group and is typically in reference to sub-population expression levels rather than the whole dataset.

Supplementary Figure 8a. **Expression of example marker genes across different cell populations identified within the cardiomyocyte lineage**.

Each panel displays a PHATE plot of the cardiomyocyte lineage with cells coloured by the expression of the specific gene identified at the top of the plot. The expression is given in log counts.

Supplementary Figure 8b. **Expression of example marker genes across different cell populations identified within the cardiomyocyte lineage**.

Each panel displays a PHATE plot of the cardiomyocyte lineage with cells coloured by the expression of the specific gene identified at the top of the plot. The expression is given in log counts.

Supplementary Figure 8c. **Expression of example marker genes across different cell populations identified within the cardiomyocyte lineage**.

Each panel displays a PHATE plot of the cardiomyocyte lineage with cells coloured by the expression of the specific gene identified at the top of the plot. The expression is given in log counts.

Supplementary Figure 9. **We can resolve several distinct CCS-associated populations within our cardiomyocyte lineage**.

A heatmap displaying cell types identified in Figure 1b from cardiomyocyte lineage with the expression of a range of CCS-associated marker genes. The annotation on the right describes the relative proportions of each cluster in terms of developmental stage. Values for expression are normalised by row and then by column for visualisation purposes.

Supplementary Figure 10. **Expression of *Dbh* across the whole dataset and across the cardiomyocyte lineage**

The top panel displays the expression of Dbh across the whole dataset UMAP plot in post-SCT normalized counts. Dbh expression can be observed primarily across cardiomyocytes and the developing brainstem/brain. The bottom panel displays Dbh expression across the cardiomyocyte lineage PHATE plot in log counts.

Supplementary Figure 11. ***Dbh* expression as a percentage of each cell type expressing <u>*Dbh*</u>**.

The top panel displays the percent of each cell type expressing *Dbh* across the cardiomyocyte lineage. The bottom panel displays *Dbh* expression in terms of absolute numbers and percentages in a tabular fashion for the same data as the top panel. *Dbh* positive expression was defined as >0 counts in the raw data.

Supplementary Figure 12. **Unsupervised clustering identified cell types across the developing murine heart using Stereo-Seq**.

a) Total bin number, nCount, nFeature, and mitochondrial percentage across different stages. Neo represents P3. Adult represents P56.

Supplementary Figure 13a. **Stereo-Seq faithfully recapitulates known gene expression changes across the developing, perinatal, and mature murine heart**.

Each image displays a representative plot of the respective gene (listed at start of row), for the respective stage (listed at top of column). High gene expression is indicated by yellow and low gene expression is indicated by purple. Expression has been normalised on a per stage basis.

Supplementary Figure 13b. **Stereo-Seq faithfully recapitulates known gene expression changes across the developing, perinatal, and mature murine heart**.

Each image displays a representative plot of the respective gene (listed at start of row), for the respective stage (listed at top of column). High gene expression is indicated by yellow and low gene expression is indicated by purple. Expression has been normalised on a per stage basis.

Supplementary Figure 13c. **Stereo-Seq faithfully recapitulates known gene expression changes across the developing, perinatal, and mature murine heart**.

Each image displays a representative plot of the respective gene (listed at start of row), for the respective stage (listed at top of column). High gene expression is indicated by yellow and low gene expression is indicated by purple. Expression has been normalised on a per stage basis.

Supplementary Figure 13d. **Stereo-Seq faithfully recapitulates known gene expression changes across the developing, perinatal, and mature murine heart**.

Each image displays a representative plot of the respective gene (listed at start of row), for the respective stage (listed at top of column). High gene expression is indicated by yellow and low gene expression is indicated by purple. Expression has been normalised on a per stage basis. N.B. *Dbh* and *Wpre vs* developmental stages have been transposed with respect to rows vs columns for genes and stages as in other parts of this figure.

Supplementary Figure 14. **Unsupervised clustering identified cell types across the developing murine heart**.

a) A dot plot demonstrating scaled expression of selected genes across cell types identified through unsupervised clustering of E12.5, E14.5, and P3 samples. Expression has been scaled for visualization purposes.

Supplementary Figure 15. **Integrated single-cell and spatial transcriptomics resolves even rare cell types across the heart, but retains mismatch between transcriptomic and biological feature importance**.

Supervised machine learning techniques can produce accurate classifiers for most cell types based on gene expression and marker gene selection.

We used SMASH to identify biologically relevant marker genes with robust computational signatures sufficient to facilitate cross-dataset and cross-platform supervised learning based classification of pixels within our spatial transcriptomics data with maximally informative, but interpretable, unique gene sets per cell type as classification label. This matrix demonstrates the highly accurate classification of cell types by a trained classifier, on our separate test data, for subsequent identification of cell type marker genes in single-cell RNA sequencing data by SMASH. Y-axis true labels refers to the set of cell-wise labels provided for our dataset, based on our initial analysis. The x-axis predicted labels are the same labels, with a high number (dark red) at the intersect of the two in the main body of the matrix indicating that the trained classifier was able to correctly identify the cell label. The scale shows the percentage of classification, with dark red indicating a higher percentage of the true label on y, was predicted to be the cell type on the x axis. Each row sums to 100.00, representing all the cells within the given cell type label on the y-axis, have been classified with predicted label(s).

b) Representatives slices from E12.5 (i), E14.5 (ii), and P3 (iii), coloured by cell type identified through RCTD predicted cell type and corresponding to cell labels in panel c.

c) Labels for cell types of respective colour in panel B.

d) i) Slices from P3 hearts, coloured by the RCTD predicted score for Purkinje fibres, with red indicating higher likelihood. ii) Slices from P3 hearts, coloured by the RCTD predicted score for the atrioventricular node, with red indicating higher likelihood. iii) Slices from P3 hearts, coloured by the RCTD predicted score for the sinoatrial node, with red indicating higher likelihood.

All slices are presented in the same anatomical orientation with respect to left to right.

Abbreviations: VM - ventricular cardiomyocytes; eAM –immature atrial cardiomyocytes; AM – atrial cardiomyocytes; eVM-trab – early trabecular ventricular cardiomyocytes; PKJ – Purkinje fibres; SAN – sinoatrial node; AM-CCS – atrial cardiac conduction system; eVM – immature ventricular cardiomyocytes; VM-trab – trabecular ventricular cardiomyocytes; AVN – atrioventricular node.

Supplementary Figure 16. **RNAscope ISH technology showing the relationships bewtween tdTomato-expressing cells with CCS and sympathetic innervation**.

a. Genetic fluorescence showing the tdTomato-expressing cells in E12.5.

b. E12.5 embryo section showing the TH(Cyan) expression in the tdTomato-expressing cell positive region, suggesting the association with sympathetic innervation.

Scale bar: 100um

SG: Sympathetic Ganglia

SV: Sinus Venosus

LA: Left Atrium

RA: Right Atrium

V: Ventricle

Supplementary Figure 17. **Spatio-temporal lineage tracing identified the association between *Dbh*^+^ cells and cardiac conduction system (CCS) at E14.5 and P3**

a. RNAscope showing the distribution of Dbh probe (green) with CCS markers including Cacna2d2 (magenta), Id2(magenta), Shox2(magenta), Tbx18(magenta) in the whole field images of E14.5

b. RNAscope showing the coexpression of Dbh probe (green) with CCS markers including Cacna2d2 (magenta), Id2(magenta), Hcn4(magenta) at RA and SAN regions at E14.5

c. RNAscope showing the distribution of Dbh (green) with CCS marker Cacna2d2 (magenta) at RA and left and right PKJ regions at P3

Suppl Figure 18a.Immunofluorescence of Dbh+ CMs in AVN+HIS bundle and Trabecular regions stained with Th (cyan) revealing the association between *Dbh*^+^ derived CMs with sympathetic innervation at E14.5.

Suppl Figure 18b. Immunofluorescence of Dbh+ derived CMs in SAN, RA,RV and RPF regions stained α-actinin (green) and Th (magenta) revealling the Dbh+ CMs and the association with sympathetic innervation at adult stage.

scale bar: 100um

LA: Left Atrium

RA: Right Atrium

LV: Left Ventricle

RV: Right Ventricle

AVN: Atrioventricular node,

HIS: Bundle of His,

Suppl Figure 19. Immunofluorescence of Dbh+ CMs in HIS and RBB regions stained α-actinin (green) and Th (magenta) revealling the Dbh+ CMs and the association with sympathetic innervation at adult stage by using *Dbh*^CreERT^/R26-tdTomato inducible reporter mouse line Scale Bar:100 μm,

**Online Video 1 (**Related to Figure 6): 3D reconstruction of *Dd*+-CMs in P3 murine heart

**Online Video 2 (**Related to Figure 6): 3D reconstruction of *Dd*+-CMs in adult murine heart

Data S1, Data S2, Data S3, Lei, Ming (2021), “3D interactive data”, Mendeley Data, V1, doi: 10.17632/hrzszgpnfp.

**Data S1 (**Related to Figure 1): A 3D interactive plot of UMAP dimensions from cell types identified in our original whole embryo (E8.5, E10.5) and whole heart (E12.5, E14.5, E16.5, P3) scRNAseq post-quality control. Cells are coloured by cell identity. Hovering over a cell produces a readout of the stage of a cell’s identity, stage, nUMI, nGene, Ratio of mtUMI/nUMI, and its position in 3D space.

**Data S2 (**Related to Figure 1): A 3D interactive plot of PHATE dimensions from cells identified within our cardiomyocyte-lineage. Cells are coloured by cell identity. Hovering over a cell produces a readout of the stage of a cell’s identity, stage, nUMI, nGene, Ratio of mtUMI/nUMI, and its position in 3D space.

**Data S3 (**Related to Figure 1): A 3D interactive plot of PHATE dimensions from *Dbh+* cells identified within our cardiomyocyte-lineage. Cells are coloured by cell identity. Hovering over a cell produces a readout of the stage of a cell’s identity, stage, nUMI, nGene, Ratio of mtUMI/nUMI, and its position in 3D space.

**Table S1 (**Related to Figure 1): Differential expression of genes between cell types within our cardiomyocyte lineage. Each sheet corresponds to its respective cell type. p_val is the pvalue for a significantly differentially expressed gene. Avg_log2FC is the average log2 fold change between for this gene compared from this cell type to all other cell types. Pct.1 is the percentage of cells expressing this gene in this given cell type, and Pct.2 is for the percentage of cells expressing this gene in all other cell types. P_val_adj is the p-value adjusted for multiple hypothesis testing by Bonferroni’s correction.

**Table S2**the top 50 genes that contributed most to the classification of each cell type by using SMASH method.

