## Supplementary material for "Integrated transcriptome and lineage analyses reveal novel catecholaminergic cardiomyocytes contributing to the cardiac conduction system in murine heart": Suppl Figures

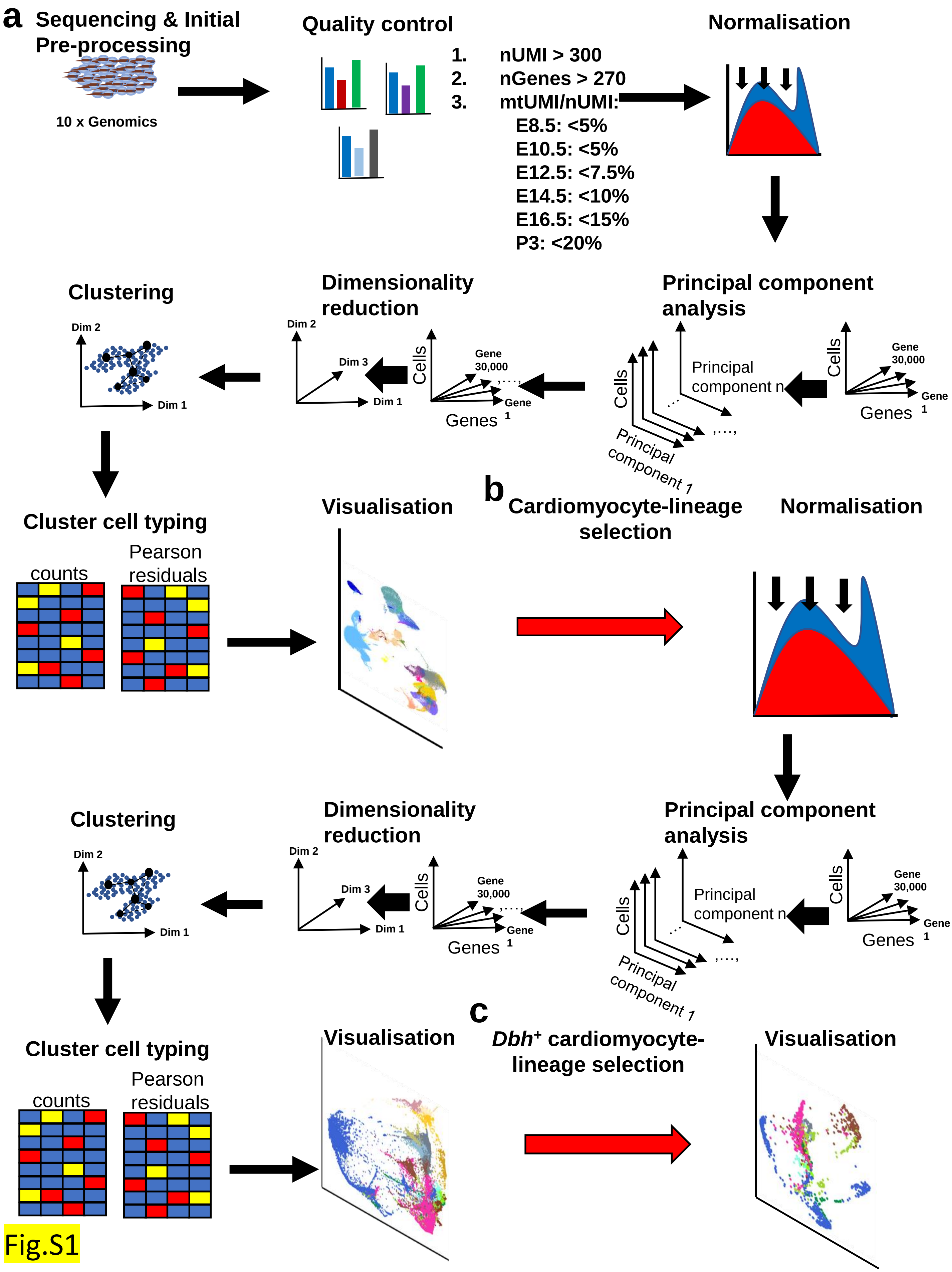
a
Sequencing & Initial Pre-processing
10 x Genomics
Quality control
1. nUMI > 300
2. nGenes > 270
3. mtUMI/nUMI:
E8.5: <5%
E10.5: <5%
E12.5: <7.5%
E14.5: <10%
E16.5: <15%
P3: <20%
Normalisation
Principal component analysis
Cells
Genes
Gene 1
Gene 30,000
Principal component n
Principal component 1
Dimensionality reduction
Dim 1
Dim 2
Dim 3
Clustering
Cluster cell typing
counts
Pearson residuals
Visualisation
b
Cardiomyocyte-lineage selection
Normalisation
Principal component analysis
Dimensionality reduction
Clustering
Cluster cell typing
counts
Pearson residuals
Visualisation
c
Dbh+ cardiomyocyte-lineage selection
Visualisation

Fig.S1

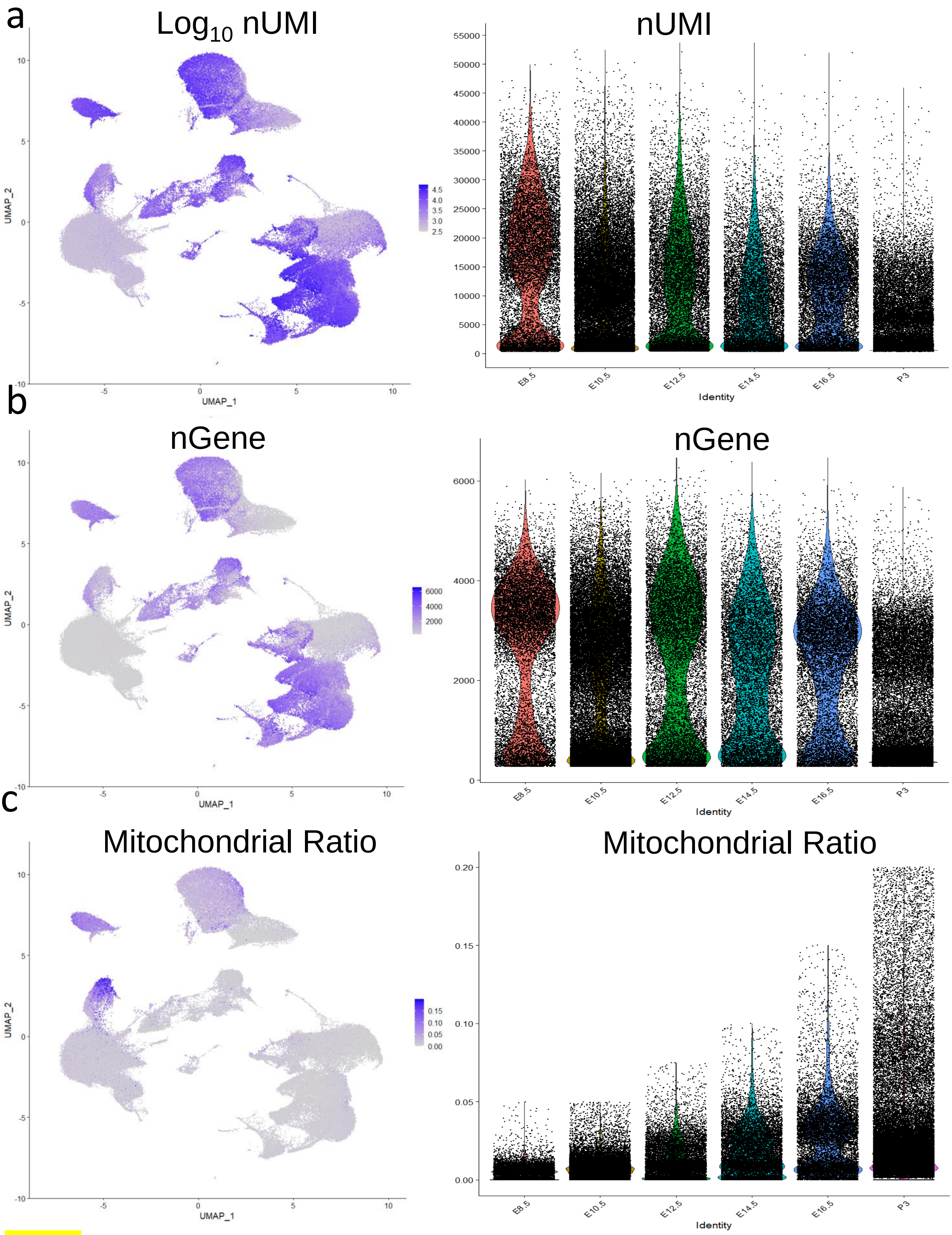

a
Log10 nUMI
nUMI
b
nGene
nGene
c
Mitochondrial Ratio
Mitochondrial Ratio
UMAP_1
UMAP_2
Identity
E8.5
E10.5
E12.5
E14.5
E16.5
P3


Fig.S2

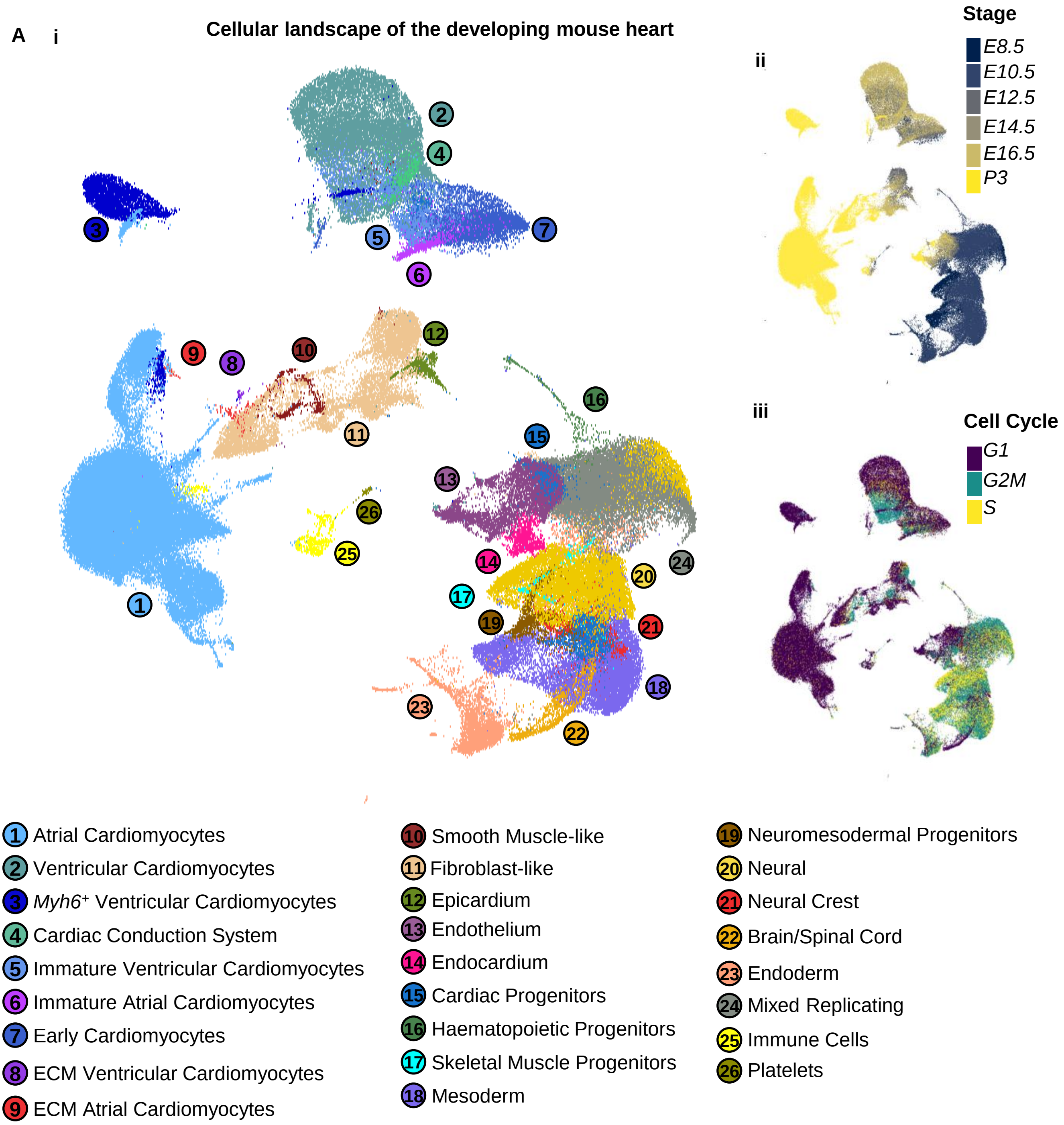
A
i
Cellular landscape of the developing mouse heart
ii
Stage
E8.5
E10.5
E12.5
E14.5
E16.5
P3
iii
Cell Cycle
G1
G2M
S
1 Atrial Cardiomyocytes
2 Ventricular Cardiomyocytes
3 Myh6+ Ventricular Cardiomyocytes
4 Cardiac Conduction System
5 Immature Ventricular Cardiomyocytes
6 Immature Atrial Cardiomyocytes
7 Early Cardiomyocytes
8 ECM Ventricular Cardiomyocytes
9 ECM Atrial Cardiomyocytes
10 Smooth Muscle-like
11 Fibroblast-like
12 Epicardium
13 Endothelium
14 Endocardium
15 Cardiac Progenitors
16 Haematopoietic Progenitors
17 Skeletal Muscle Progenitors
18 Mesoderm
19 Neuromesodermal Progenitors
20 Neural
21 Neural Crest
22 Brain/Spinal Cord
23 Endoderm
24 Mixed Replicating
25 Immune Cells
26 Platelets

Fig.S3

**Mixed Replicating**
*Snrpg*$^{hi}$
*Rp-genehi*
*Ptma*$^{hi}$
*Other*

**Mesoderm**
*Foxd1*$^{+}$
*Prrx1*$^{+}$
*Meox1*$^{+}$

**Neural**
*Sox2*$^{+}$
*Otx2*$^{+}$
*Pax6*$^{+}$

**Endoderm**
Pyy$^{+}$
Epcam$^{+}$
Afp$^{+}$
Ttr$^{+}$

**Cardiac Progenitors**
Mesoderm Genes$^{+}$
*Hand1*$^{hi}$
*Wnt2*$^{hi}$
*Bmp4*$^{+}$

**Haematopoietic Progenitors**
*Redrum*$^{+}$
*Hemgn*$^{+}$
*Hbb-genes*$^{+}$

**Skeletal Muscle Progenitors**
Mesoderm Genes$^{+}$
*Myf6*$^{+}$
*Myog*$^{+}$

**Neuromesodermal Progenitors**
Neural Genes$^{+}$
Mesoderm Genes$^{+}$
*Sox2*$^{+}$
*T*$^{hi}$

**Neural Crest**
Neural Genes$^{+}$
*Sox10*$^{+}$
*Foxd3*$^{+}$

**Early Cardiomyocytes**
*Tnni1*$^{hi}$
*Smpx*$^{hi}$
*Tnni3*$^{lo}$
*Myl7*$^{+}$
*Tagln*$^{+}$
*Pln*$^{+}$
*Atp2a2*$^{lo}$

**Endocardium**
Endothelium Genes$^{+}$
*Etv2*$^{hi}$
*Tal1*$^{+}$

**Endothelium**
*Ecscr*$^{hi}$
*Emcn*$^{hi}$
*Cdh5*$^{hi}$
*Kdr*$^{hi}$

**Epicardium**
*Upk3b*$^{hi}$
*Upk1b*$^{hi}$
*Aldh1a2*$^{hi}$

**Smooth Muscle-like**
*Rgs5*$^{hi}$
*Myh11*$^{hi}$
*Lmod1*$^{hi}$

**Fibroblast-like**
*Fbln2*$^{hi}$
*Fbn1*$^{hi}$
*Postn*$^{hi}$
*Col1a1*$^{hi}$

**Brain/Spinal Cord**
Neural Genes$^{+}$
*Isl1*$^{+}$
*Neurog1*$^{+}$
*Onecut2*$^{+}$

**Early Ventricular Cardiomyocytes**
Early Cardiomyocyte Genes$^{+}$
*Myl7*$^{lo}$
*Myl2*$^{hi}$
*Myh7*$^{+}$
*Pln*$^{+}$

**Early Atrial Cardiomyocytes**
Early Cardiomyocyte Genes$^{+}$
*Myl7*$^{hi}$
*Myl2*$^{lo}$
*Sln*$^{+}$

**Platelets**
*Ppbp*$^{hi}$
*Pf4*$^{hi}$
*Gp1bb*$^{hi}$
*Gp6*$^{hi}$

**Ventricular Cardiomyocytes**
Atrial Genes$^{lo/-}$
*Myl2*$^{hi}$
*Myh7*$^{hi}$
*Kcne1*$^{hi}$
*Gja1*$^{hi}$

***Cardiac Conduction System***
*Hcn4*$^{+}$
*Cacna2d2*$^{hi}$
*Tbx3*$^{+}$
*Shox2*$^{+}$
*Cacna1g*$^{+}$

**Atrial Cardiomyocytes**
Ventricular Genes$^{lo/-}$
*Myl7*$^{hi}$
*Nppa*$^{hi}$
*Sln*$^{hi}$
*Myh6*$^{+}$

**Immune Cells**
*C1qb*$^{hi}$
*Tyrobp*$^{hi}$
*C1qc*$^{hi}$
*Fcer1g*$^{hi}$
*Lyz2*$^{hi}$
*Fcgr3*$^{hi}$

**ECM Ventricular Cardiomyocytes**
Ventricular Cardiomyocyte Genes$^{+}$
Fbn1$^{+}$
Postn$^{+}$
Col1a1$^{+}$

***Myh6*$^{+}$ Ventricular Cardiomyocytes**
Ventricular Genes$^{hi}$
Atrial Genes$^{lo}$
*Myh6*$^{hi}$

**ECM Atrial Cardiomyocytes**
Atrial Cardiomyocyte Genes$^{+}$
Fbn1$^{+}$
Postn$^{+}$
Col1a1$^{+}$

Fig.S4

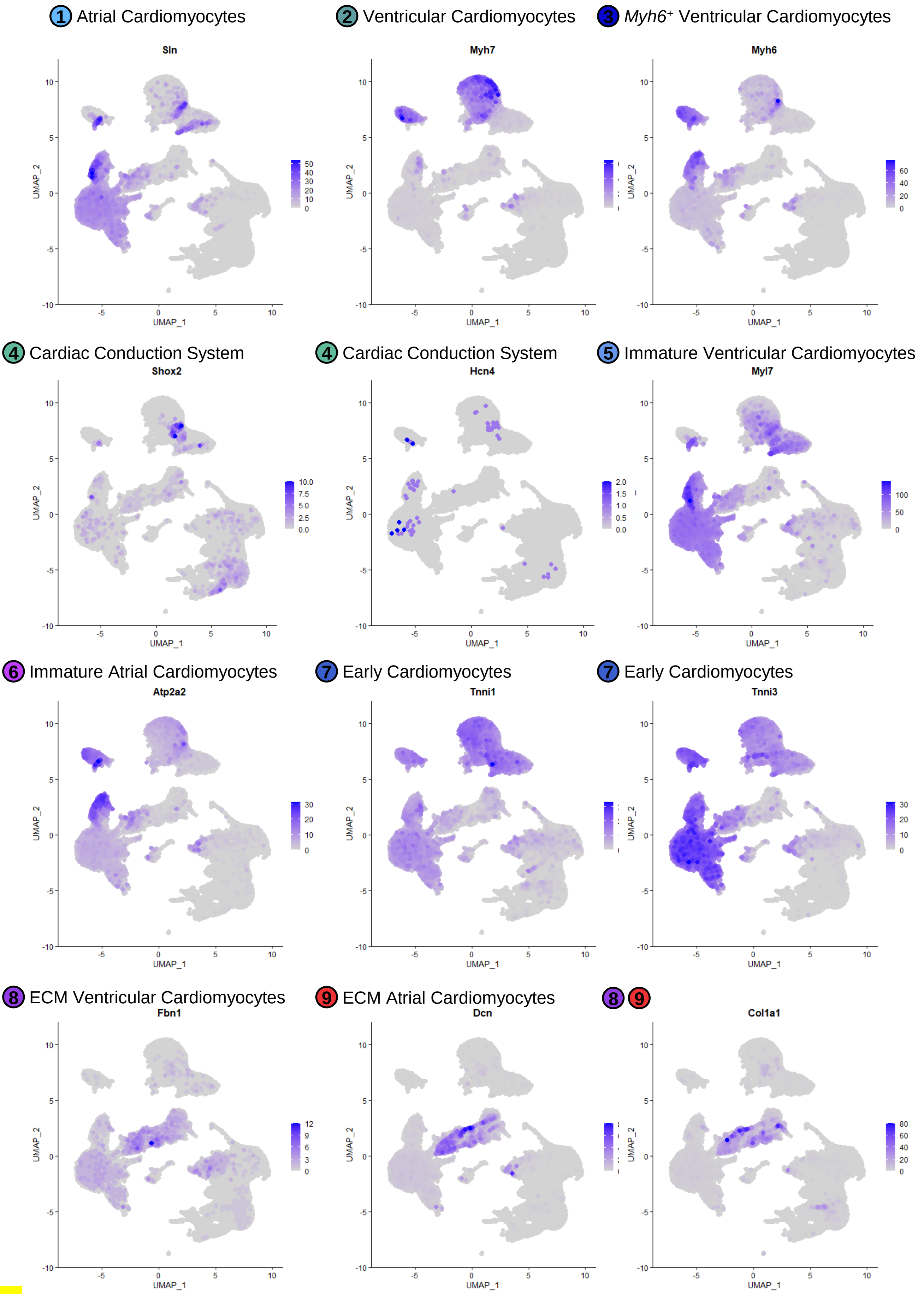
1 Atrial Cardiomyocytes
Sln
2 Ventricular Cardiomyocytes
Myh7
3 *Myh6*+ Ventricular Cardiomyocytes
Myh6
4 Cardiac Conduction System
Shox2
4 Cardiac Conduction System
Hcn4
5 Immature Ventricular Cardiomyocytes
Myl7
6 Immature Atrial Cardiomyocytes
Atp2a2
7 Early Cardiomyocytes
Tnni1
7 Early Cardiomyocytes
Tnni3
8 ECM Ventricular Cardiomyocytes
Fbn1
9 ECM Atrial Cardiomyocytes
Dcn
8 9
Col1a1
UMAP_1
UMAP_2

Fig.S5a

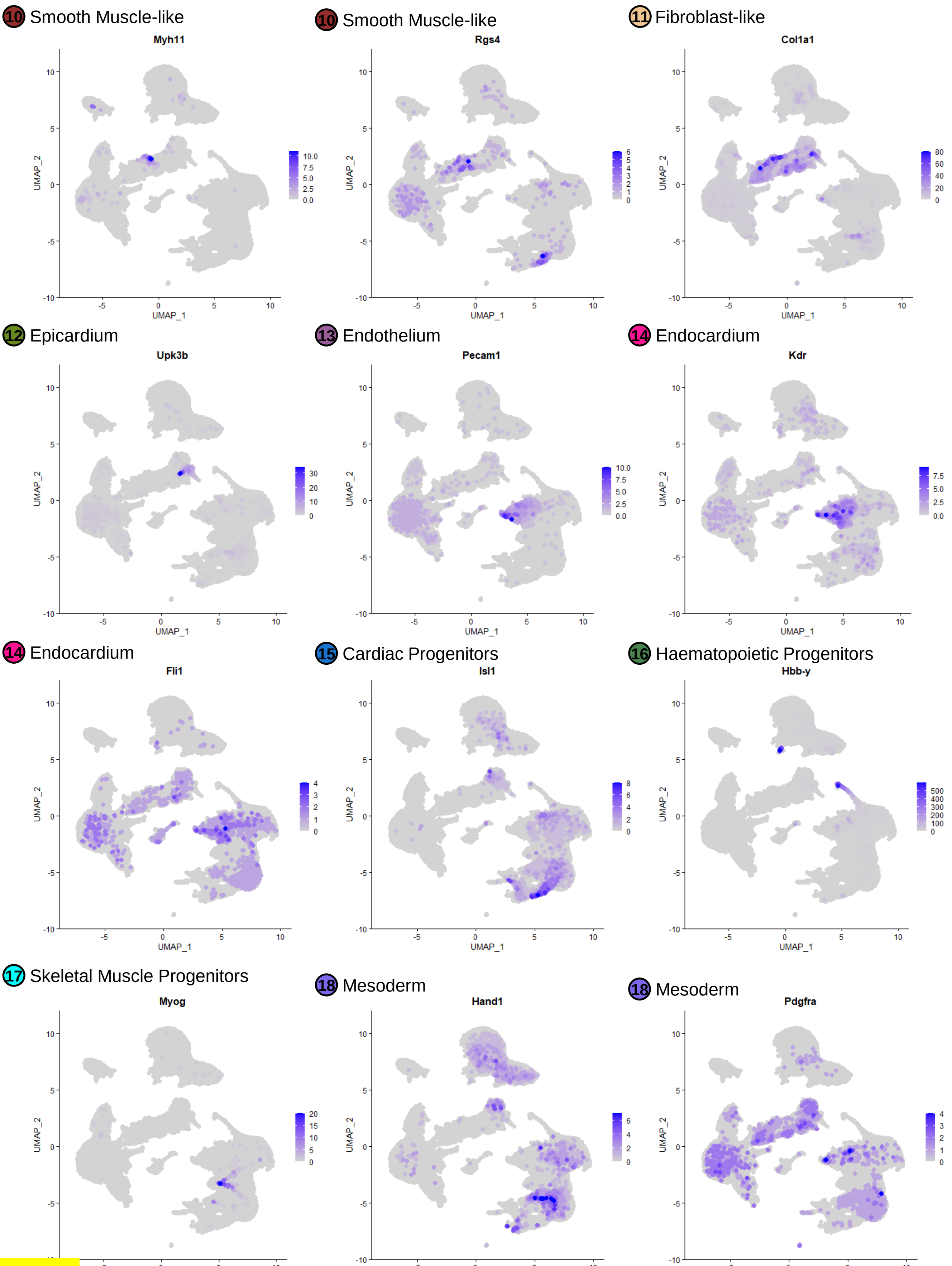
10 Smooth Muscle-like
Myh11
10 Smooth Muscle-like
Rgs4
11 Fibroblast-like
Col1a1
12 Epicardium
Upk3b
13 Endothelium
Pecam1
14 Endocardium
Kdr
14 Endocardium
Fli1
15 Cardiac Progenitors
Isl1
16 Haematopoietic Progenitors
Hbb-y
17 Skeletal Muscle Progenitors
Myog
18 Mesoderm
Hand1
18 Mesoderm
Pdgfra
UMAP_1
UMAP_2

Fig.S5b

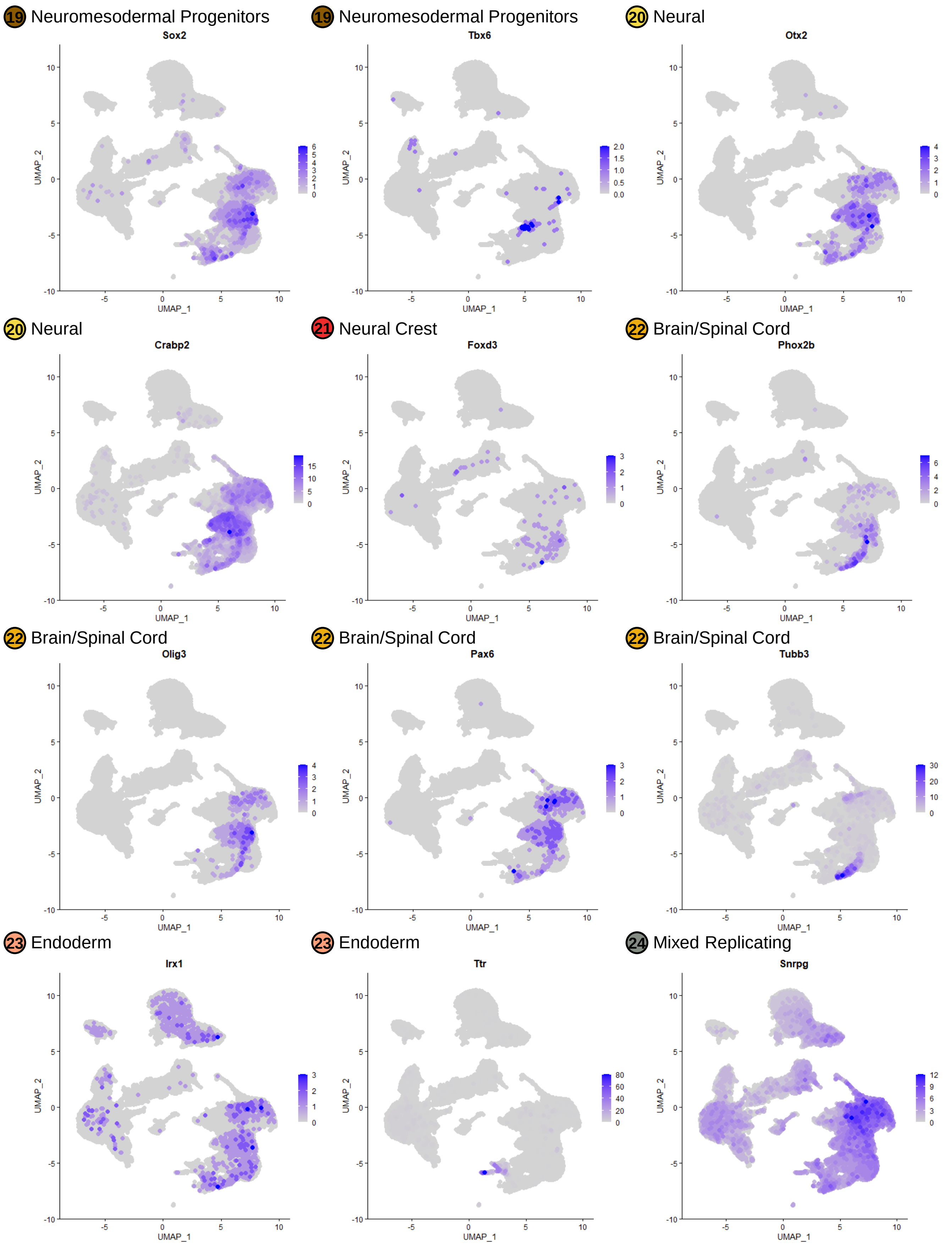

19 Neuromesodermal Progenitors
Sox2
19 Neuromesodermal Progenitors
Tbx6
20 Neural
Otx2
20 Neural
Crabp2
21 Neural Crest
Foxd3
22 Brain/Spinal Cord
Phox2b
22 Brain/Spinal Cord
Olig3
22 Brain/Spinal Cord
Pax6
22 Brain/Spinal Cord
Tubb3
23 Endoderm
Irx1
23 Endoderm
Ttr
24 Mixed Replicating
Snrpg
UMAP_1
UMAP_2


Fig.S5c

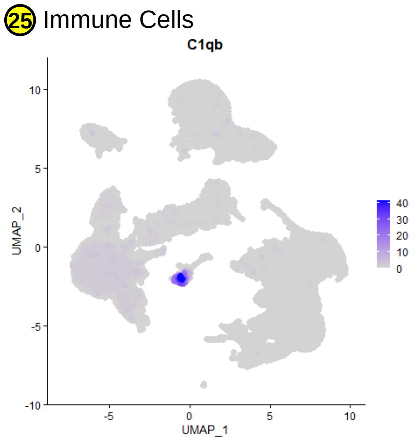
25 Immune Cells
C1qb
UMAP_2
UMAP_1
40
30
20
10
0

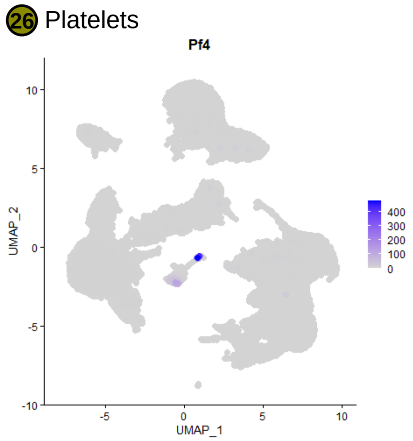
26 Platelets
Pf4
UMAP_2
UMAP_1
400
300
200
100
0

Fig.S5d

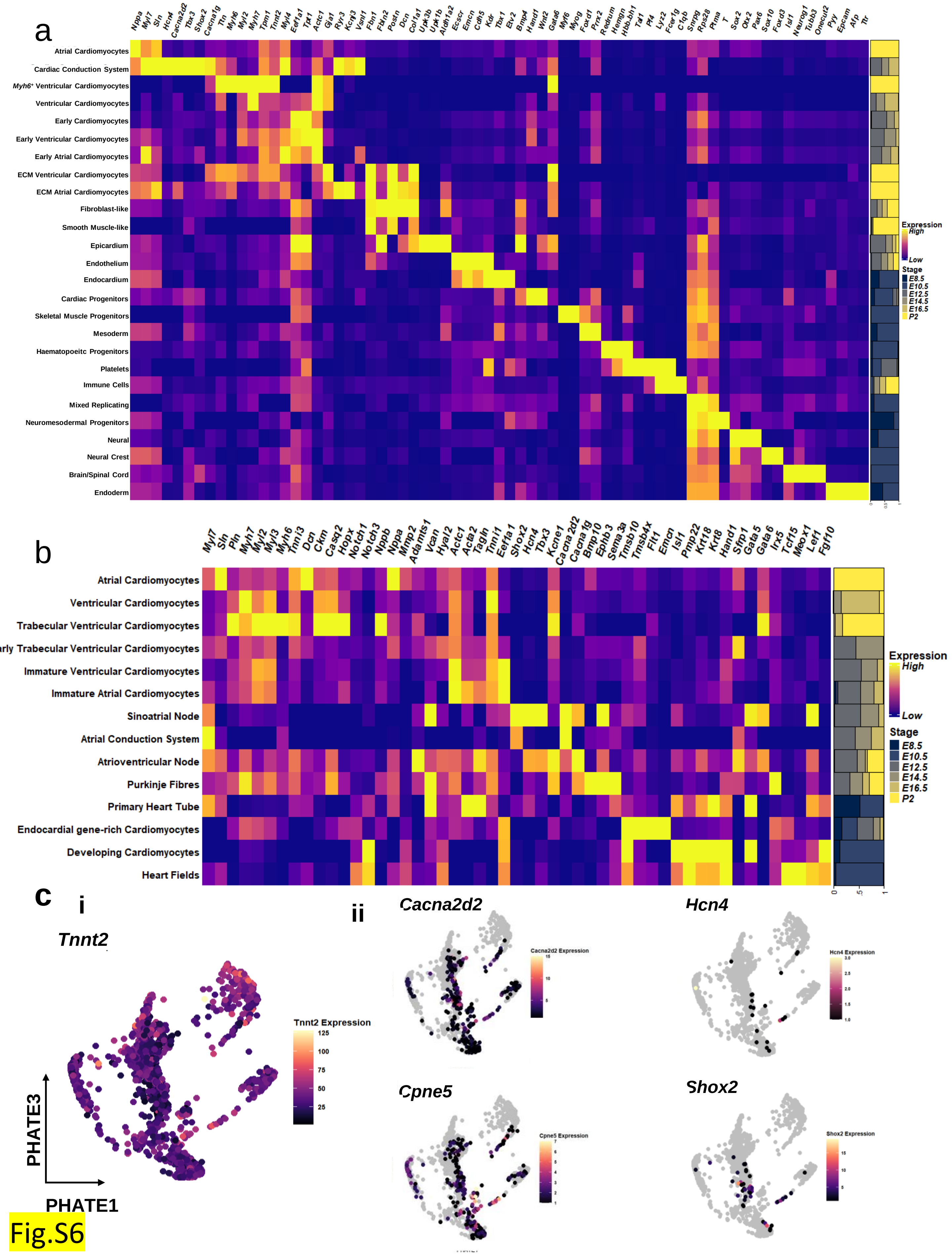

a
Nppa Myl7 Sln Hcn4 Cacna2d2 Tbx3 Shox2 Cacna1g Ttn Myh6 Myl2 Myh7 Tpm1 Tnnt2 Myl4 Eef1a1 Tpt1 Actc1 Gja1 Ryr3 Kcnj3 Vsnl1 Fbn1 Fbln2 Postn Dcn Col1a1 Upk3b Upk1b Aldh1a2 Ecscr Emcn Cdh5 Kdr Tbx1 Etv2 Bmp4 Hand1 Wnt2 Gata6 Myf6 Myog Foxd1 Prrx2 Redrum Hemgn Hbb.bh1 Tal1 Pf4 Lyz2 Fcer1g C1qb Snrpg Rps28 Ptma T Sox2 Otx2 Pax6 Sox10 Foxd3 Isl1 Neurog1 Tubb3 Onecut2 Pyy Epcam Afp Ttr
Atrial Cardiomyocytes
Cardiac Conduction System
Myh6+ Ventricular Cardiomyocytes
Ventricular Cardiomyocytes
Early Cardiomyocytes
Early Ventricular Cardiomyocytes
Early Atrial Cardiomyocytes
ECM Ventricular Cardiomyocytes
ECM Atrial Cardiomyocytes
Fibroblast-like
Smooth Muscle-like
Epicardium
Endothelium
Endocardium
Cardiac Progenitors
Skeletal Muscle Progenitors
Mesoderm
Haematopoeitc Progenitors
Platelets
Immune Cells
Mixed Replicating
Neuromesodermal Progenitors
Neural
Neural Crest
Brain/Spinal Cord
Endoderm
Expression
High
Low
Stage
E8.5
E10.5
E12.5
E14.5
E16.5
P2
b
Myl7 Sln Pln Myh7 Myl2 Myl3 Myh6 Tnni3 Dcn Ckm Casq2 Hopx Notch1 Notch3 Nppb Nppa Mmp2 Adamts1 Vcan Hyal2 Actc1 Acta2 Tagln Tnni1 Eef1a1 Shox2 Hcn4 Tbx3 Kcne1 Cacna2d2 Cacna1g Bmp10 Ephb3 Sema3a Tmsb10 Tmsb4x Flt1 Emcn Isl1 Pmp22 Krt18 Krt8 Hand1 Sfrp1 Gata5 Gata6 Irx5 Tcf15 Meox1 Lef1 Fgf10
Atrial Cardiomyocytes
Ventricular Cardiomyocytes
Trabecular Ventricular Cardiomyocytes
rly Trabecular Ventricular Cardiomyocytes
Immature Ventricular Cardiomyocytes
Immature Atrial Cardiomyocytes
Sinoatrial Node
Atrial Conduction System
Atrioventricular Node
Purkinje Fibres
Primary Heart Tube
Endocardial gene-rich Cardiomyocytes
Developing Cardiomyocytes
Heart Fields
Expression
High
Low
Stage
E8.5
E10.5
E12.5
E14.5
E16.5
P2
c
i
Tnnt2
Tnnt2 Expression
125
100
75
50
25
PHATE3
PHATE1
ii
Cacna2d2
Hcn4
Cpne5
Shox2


Fig.S6

**Heart Fields**
*Osr1*$^{hi}$
*Hand1*$^{hi}$
*Wnt2*$^{hi}$
*Krt8*$^{+}$

**Miscellaneous**
Markers for other cell types$^{-/lo}$
Cell damage markers$^{hi}$
E.g. *Dnajb3*$^{hi}$

**Developing Cardiomyocytes**
*Acta2*$^{hi}$
*Hmga2*$^{hi}$
*Pmp22*$^{hi}$
*Krt8/18*$^{hi}$
*Isl1*$^{hi}$
*Notch3*$^{hi}$

**Endocardial Gene-rich Cardiomyocytes**
*Myl7*$^{+}$
*Myl2*$^{+}$
*Tnni1*$^{+}$
*Tmsb4x*$^{hi}$
*Flt1*$^{hi}$
*Emcn*$^{hi}$

**Primary Heart Tube**
Developing Cardiomyocyte Genes$^{+}$
*Isl1*$^{lo}$
*Notch3*$^{lo}$
*Acta2*$^{+}$
*Tagln*$^{+}$

**Immature Atrial Cardiomyocytes**
Atrial Cardiomyocyte Genes$^{lo}$
*Tnni3*$^{lo}$
*Tnni1*$^{hi}$
*Acta2*$^{hi}$

**Immature Ventricular Cardiomyocytes**
Ventricular Cardiomyocyte Genes$^{lo}$
*Tnni3*$^{lo}$
*Tnni1*$^{hi}$
*Acta2*$^{hi}$

**Early Trabecular Ventricular Cardiomyocytes**
Ventricular Cardiomyocyte Genes$^{+}$
*Nppb*$^{+}$
*Nppa*$^{+}$
*Vcan*$^{hi}$
*Hyal2*$^{hi}$
*Tnni1*$^{hi}$

**Atrial Cardiomyocytes**
*Myl7*$^{hi}$
*Myh6*$^{hi}$
*Myl4*$^{hi}$
*Nppa*$^{hi}$
*Sln*$^{hi}$
*Tnni3*$^{hi}$

**Ventricular Cardiomyocytes**
*Myh7*$^{hi}$
*Myl2*$^{hi}$
*Pln*$^{hi}$
*Tnni3*$^{hi}$

**Trabecular Ventricular Cardiomyocytes**
Ventricular Cardiomyocyte Genes$^{+}$
*Myh6*$^{hi}$
*Nppb*$^{hi}$
*Hopx*$^{hi}$
*Nppa*$^{+}$
*Casq2*$^{hi}$

**Atrial Cardiac Conduction System**
*Myl7*$^{+}$
*Tnnt2*$^{+}$
*Cacna2d2*$^{hi}$
*Igfbp5*$^{hi}$
*Nppa*$^{lo}$

**Sinoatrial Node**
*Myl7*$^{+}$
*Tnnt2*$^{+}$
*Cacna2d2*$^{hi}$
*Shox2*$^{hi}$
*Igfbp5*$^{hi}$
*Vsnl1*$^{+}$
*Hcn4*$^{+}$

**Atrioventricular Node**
*Myl7*$^{+}$
*Tnnt2*$^{+}$
*Rspo3*$^{hi}$
*Cacna2d2*$^{hi}$
*Cacna1g*$^{hi}$
*Bmp2*$^{hi}$
*Igfbp5*$^{hi}$
*Shox2*$^{-}$
*Vsnl1*$^{-}$

**Purkinje Fibres**
Trabecular Ventricular Cardiomyocyte Genes$^{+}$
*Nppa*$^{lo}$
*Nppb*$^{lo}$
*Igfbp5*$^{hi}$
*Slit2*$^{hi}$
*Sema3a*$^{hi}$
*Cacna1g*$^{hi}$
*Ephb3*$^{hi}$

Fig.S7

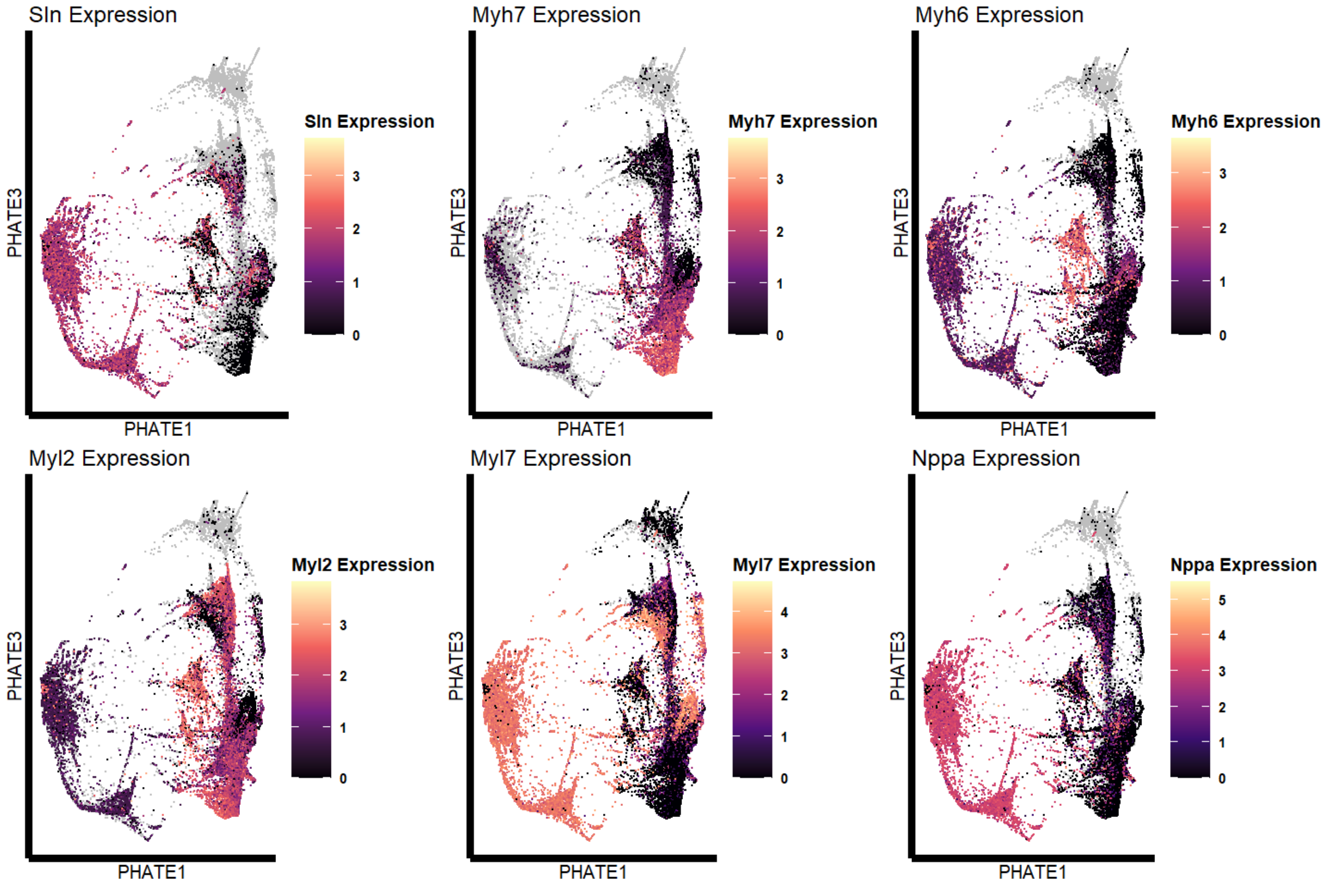
Sln Expression
PHATE3
PHATE1
Sln Expression
3
2
1
0
Myh7 Expression
PHATE3
PHATE1
Myh7 Expression
3
2
1
0
Myh6 Expression
PHATE3
PHATE1
Myh6 Expression
3
2
1
0
Myl2 Expression
PHATE3
PHATE1
Myl2 Expression
3
2
1
0
Myl7 Expression
PHATE3
PHATE1
Myl7 Expression
4
3
2
1
0
Nppa Expression
PHATE3
PHATE1
Nppa Expression
5
4
3
2
1
0

Fig.S8a

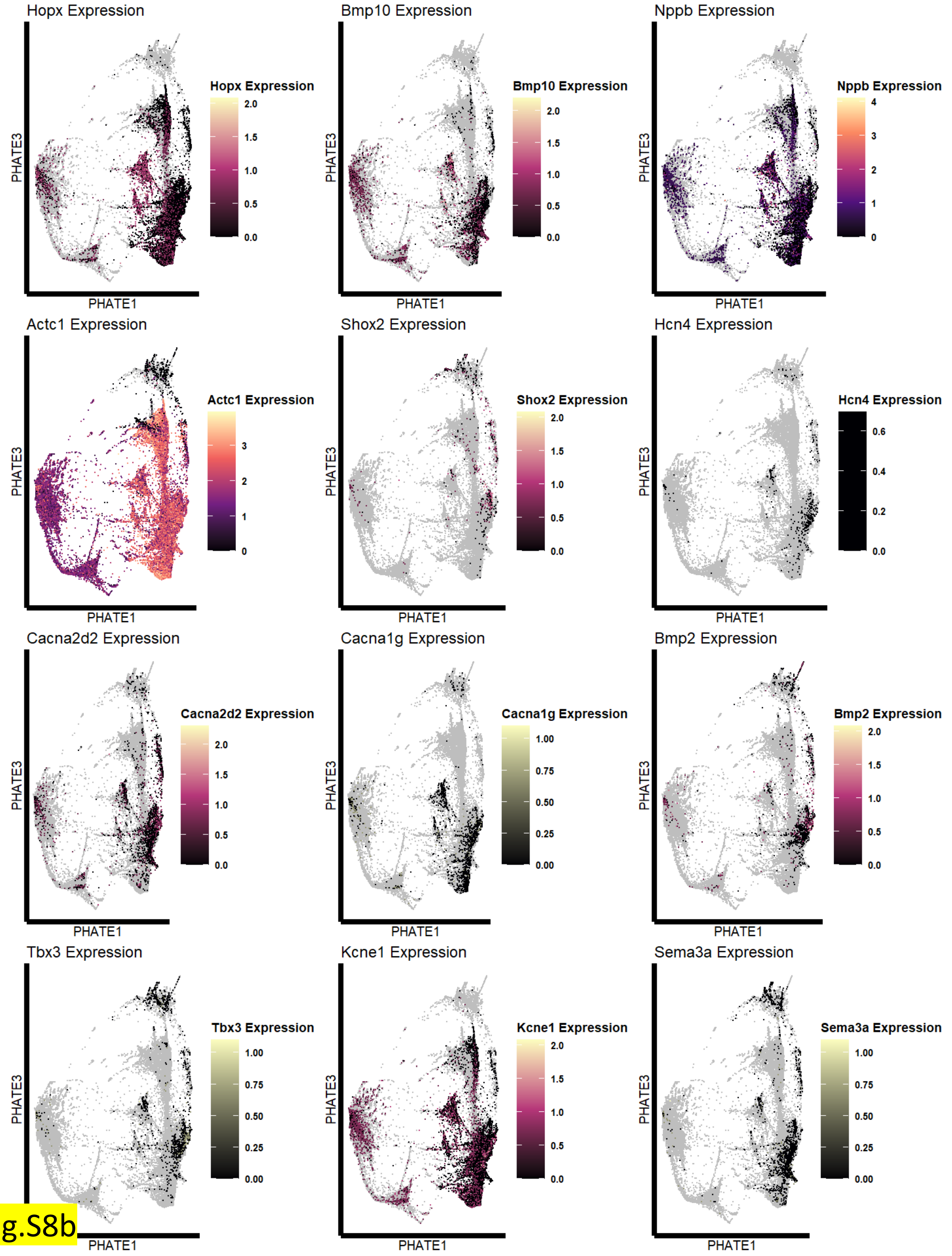
Hopx Expression
PHATE3
PHATE1
Hopx Expression
2.0
1.5
1.0
0.5
0.0
Bmp10 Expression
PHATE3
PHATE1
Bmp10 Expression
2.0
1.5
1.0
0.5
0.0
Nppb Expression
PHATE3
PHATE1
Nppb Expression
4
3
2
1
0
Actc1 Expression
PHATE3
PHATE1
Actc1 Expression
3
2
1
0
Shox2 Expression
PHATE3
PHATE1
Shox2 Expression
2.0
1.5
1.0
0.5
0.0
Hcn4 Expression
PHATE3
PHATE1
Hcn4 Expression
0.6
0.4
0.2
0.0
Cacna2d2 Expression
PHATE3
PHATE1
Cacna2d2 Expression
2.0
1.5
1.0
0.5
0.0
Cacna1g Expression
PHATE3
PHATE1
Cacna1g Expression
1.00
0.75
0.50
0.25
0.00
Bmp2 Expression
PHATE3
PHATE1
Bmp2 Expression
2.0
1.5
1.0
0.5
0.0
Tbx3 Expression
PHATE3
PHATE1
Tbx3 Expression
1.00
0.75
0.50
0.25
0.00
Kcne1 Expression
PHATE3
PHATE1
Kcne1 Expression
2.0
1.5
1.0
0.5
0.0
Sema3a Expression
PHATE3
PHATE1
Sema3a Expression
1.00
0.75
0.50
0.25
0.00

Fig.S8b

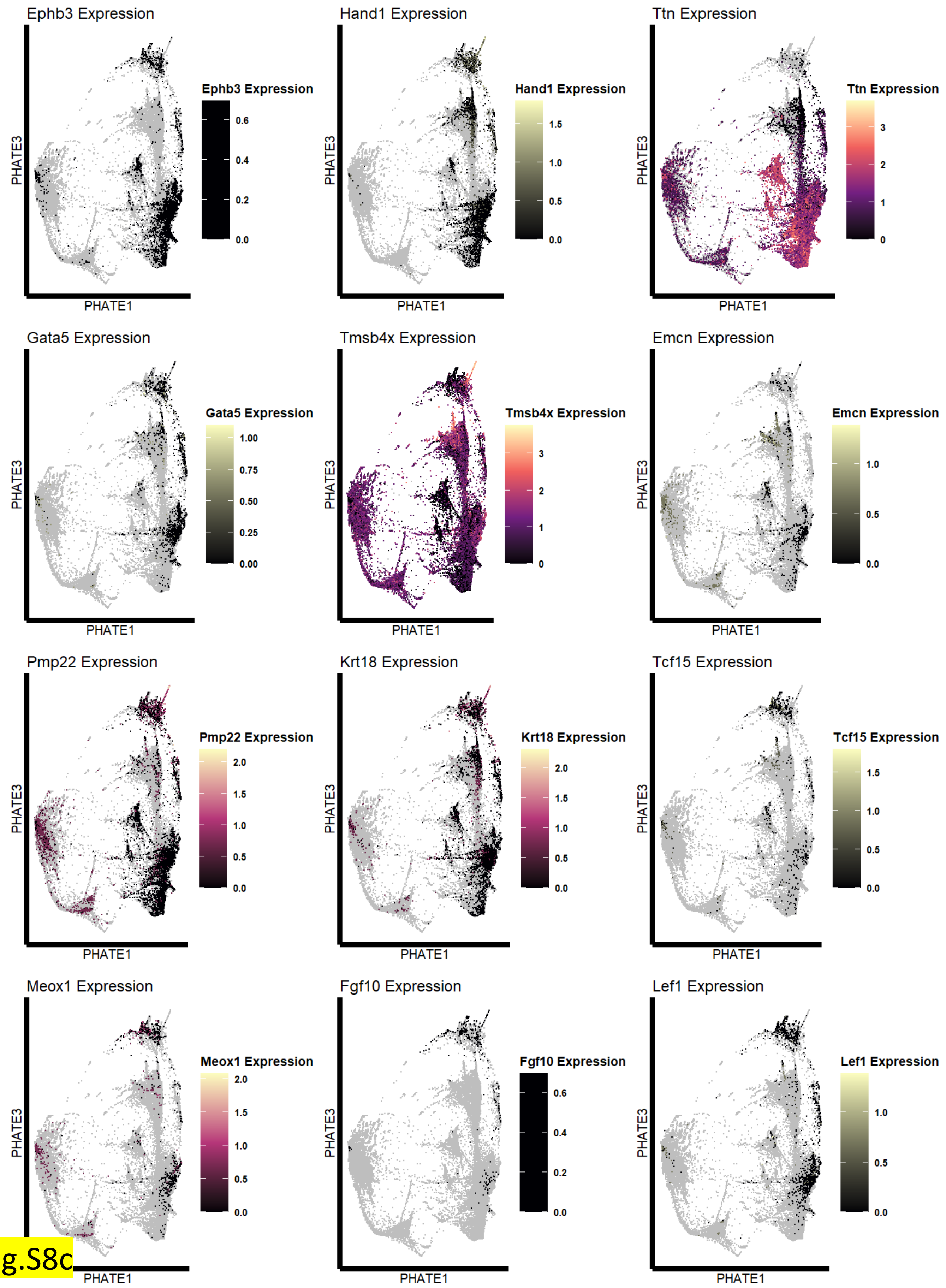

Ephb3 Expression
Ephb3 Expression
0.6
0.4
0.2
0.0
PHATE3
PHATE1
Hand1 Expression
Hand1 Expression
1.5
1.0
0.5
0.0
PHATE3
PHATE1
Ttn Expression
Ttn Expression
3
2
1
0
PHATE3
PHATE1
Gata5 Expression
Gata5 Expression
1.00
0.75
0.50
0.25
0.00
PHATE3
PHATE1
Tmsb4x Expression
Tmsb4x Expression
3
2
1
0
PHATE3
PHATE1
Emcn Expression
Emcn Expression
1.0
0.5
0.0
PHATE3
PHATE1
Pmp22 Expression
Pmp22 Expression
2.0
1.5
1.0
0.5
0.0
PHATE3
PHATE1
Krt18 Expression
Krt18 Expression
2.0
1.5
1.0
0.5
0.0
PHATE3
PHATE1
Tcf15 Expression
Tcf15 Expression
1.5
1.0
0.5
0.0
PHATE3
PHATE1
Meox1 Expression
Meox1 Expression
2.0
1.5
1.0
0.5
0.0
PHATE3
PHATE1
Fgf10 Expression
Fgf10 Expression
0.6
0.4
0.2
0.0
PHATE3
PHATE1
Lef1 Expression
Lef1 Expression
1.0
0.5
0.0
PHATE3
PHATE1


Fig.S8c

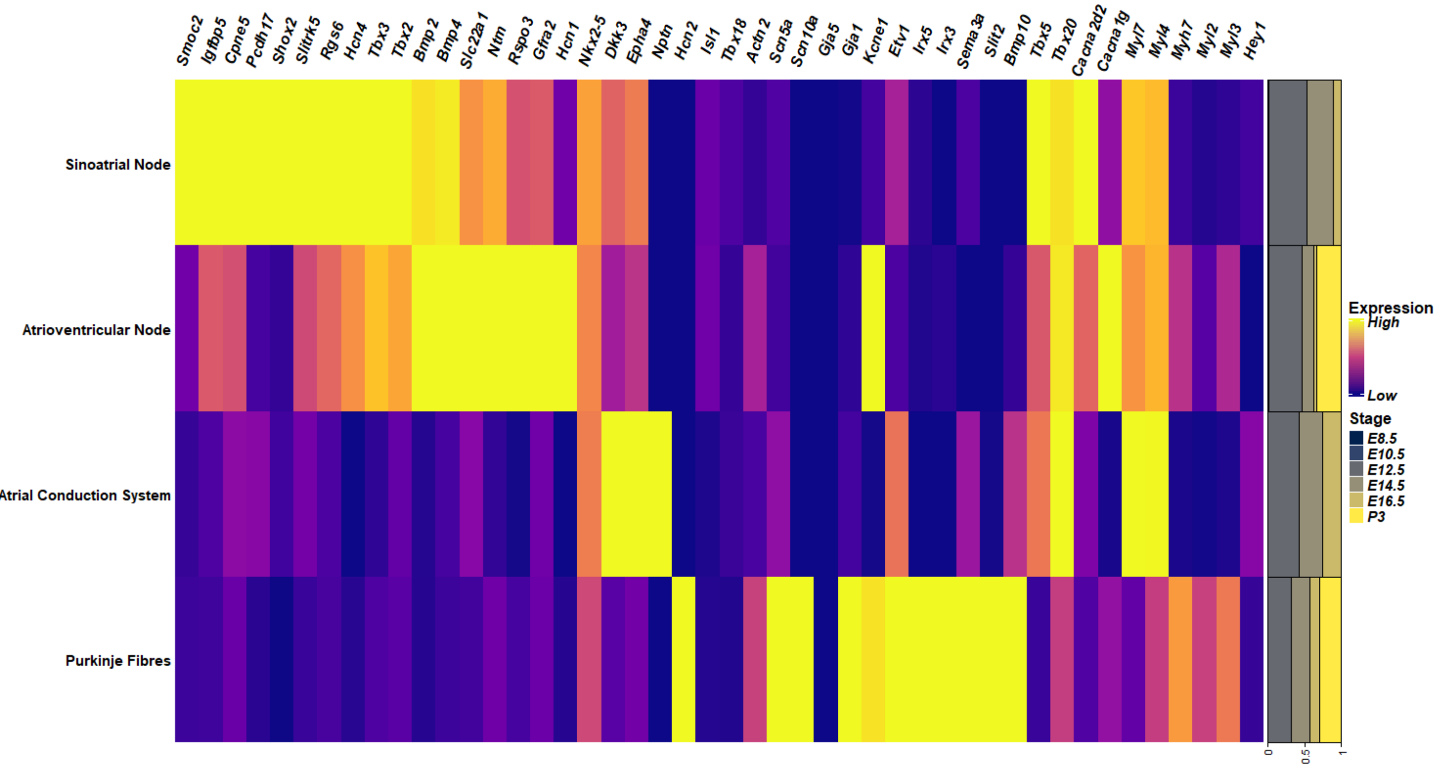

Smoc2
Igfbp5
Cpne5
Pcdh17
Shox2
Slitrk5
Rgs6
Hcn4
Tbx3
Tbx2
Bmp2
Bmp4
Slc22a1
Ntm
Rspo3
Gfra2
Hcn1
Nkx2-5
Dkk3
Epha4
Nptn
Hcn2
Isl1
Tbx18
Actn2
Scn5a
Scn10a
Gja5
Gja1
Kcne1
Etv1
Irx5
Irx3
Sema3a
Slit2
Bmp10
Tbx5
Tbx20
Cacna2d2
Cacna1g
Myl7
Myl4
Myh7
Myl2
Myl3
Hey1
Sinoatrial Node
Atrioventricular Node
Atrial Conduction System
Purkinje Fibres
Expression
High
Low
Stage
E8.5
E10.5
E12.5
E14.5
E16.5
P3
0
0.5
1


Fig.S9

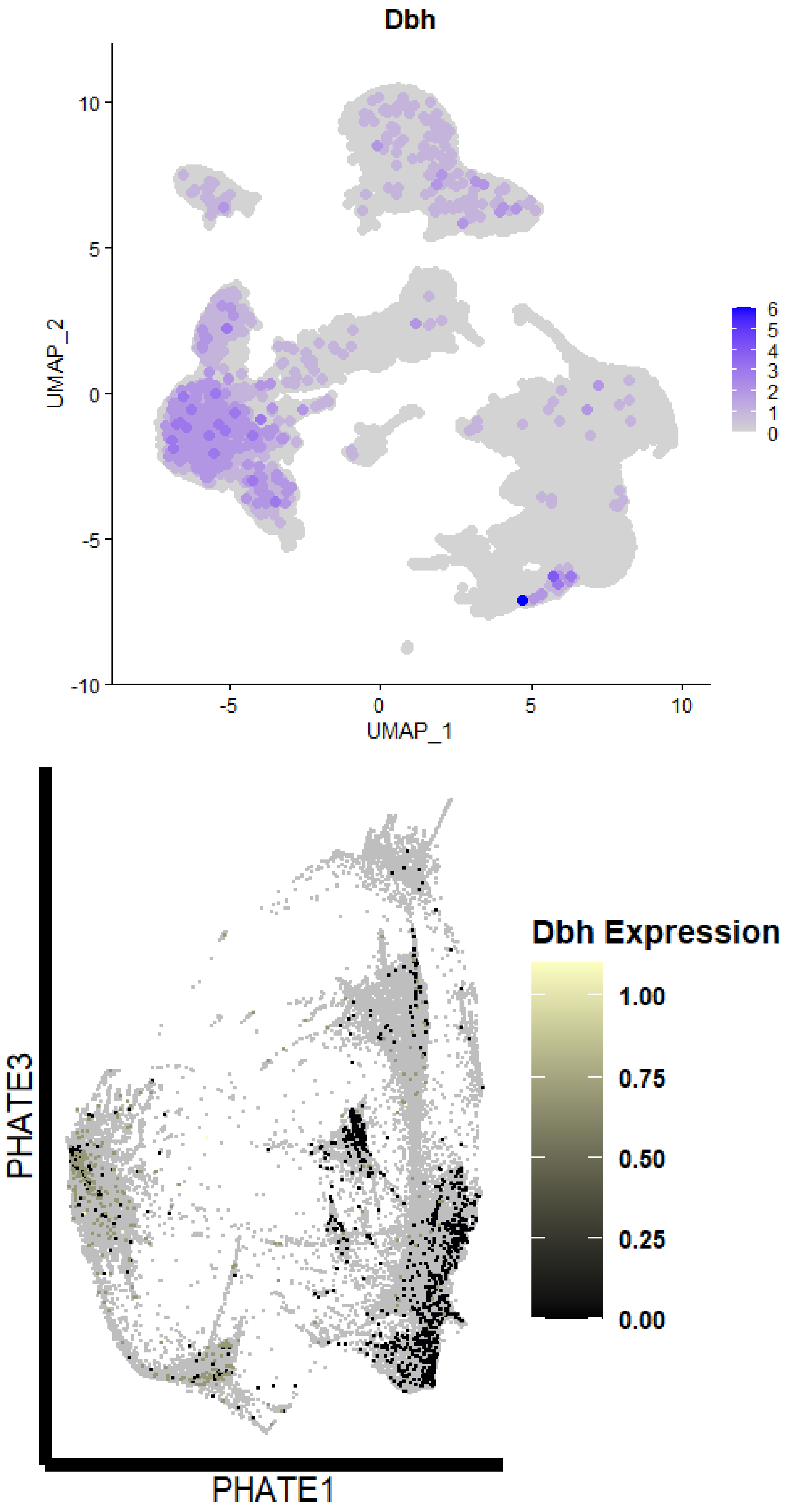
Dbh
10
5
0
-5
-10
UMAP_2
-5
0
5
10
UMAP_1
6
5
4
3
2
1
0
PHATE3
PHATE1
Dbh Expression
1.00
0.75
0.50
0.25
0.00

Fig.S10

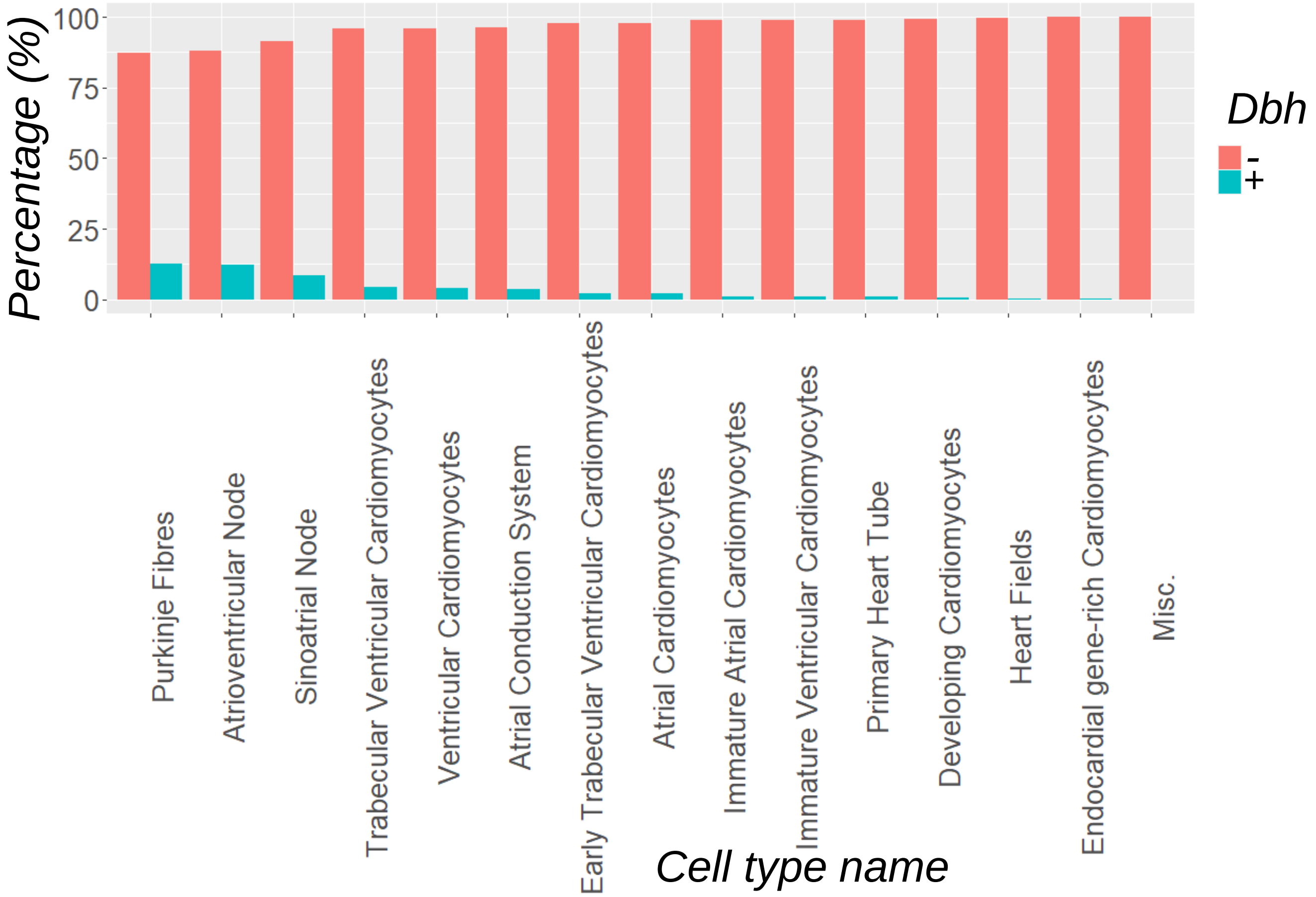


| Cell type | Total number of cells in type (N) | Number of cells *Dbh+* (N) | Number of cells *Dbh-* (N) | Percentage *Dbh+* (%) |
|---|---|---|---|---|
| Atrial Cardiomyocytes | 74021 | 1575 | 72446 | 2.1 |
| Early Trabecular Ventricular Cardiomyocytes | 5439 | 123 | 5316 | 2.3 |
| Heart Fields | 1783 | 5 | 1778 | 0.3 |
| Ventricular Cardiomyocytes | 7692 | 300 | 7392 | 3.9 |
| Immature Ventricular Cardiomyocytes | 6897 | 72 | 6825 | 1.0 |
| Atrioventricular Node | 486 | 59 | 427 | 12.1 |
| Purkinje Fibres | 1606 | 205 | 1401 | 12.8 |
| Primary Heart Tube | 561 | 5 | 556 | 0.9 |
| Trabecular Ventricular Cardiomyocytes | 4069 | 170 | 3899 | 4.2 |
| Immature Atrial Cardiomyocytes | 1752 | 17 | 1735 | 1.0 |
| Endocardial gene-rich Cardiomyocytes | 1407 | 2 | 1405 | 0.1 |
| Developing Cardiomyocytes | 793 | 5 | 788 | 0.6 |
| Sinoatrial Node | 286 | 24 | 262 | 8.4 |
| Atrial Conduction System | 760 | 29 | 731 | 3.8 |
| | 153 | 0 | 153 | 0.0 |

Fig.S11

**a**

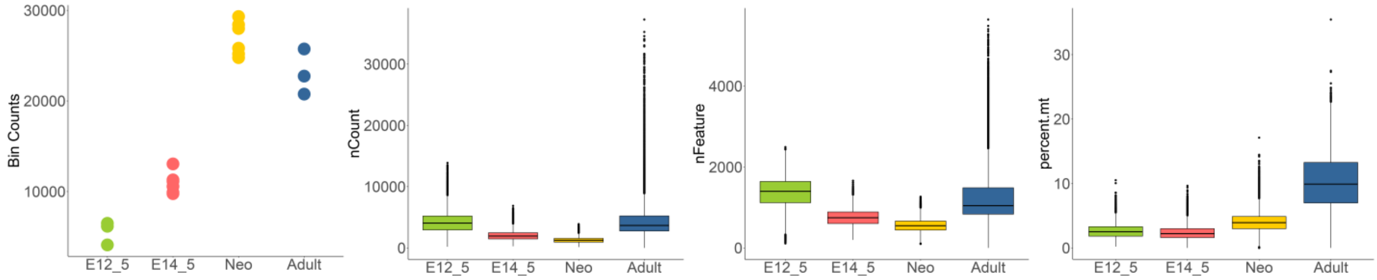

Bin Counts
30000
20000
10000
E12_5
E14_5
Neo
Adult
nCount
30000
20000
10000
0
nFeature
4000
2000
0
percent.mt
30
20
10
0


Fig.S12

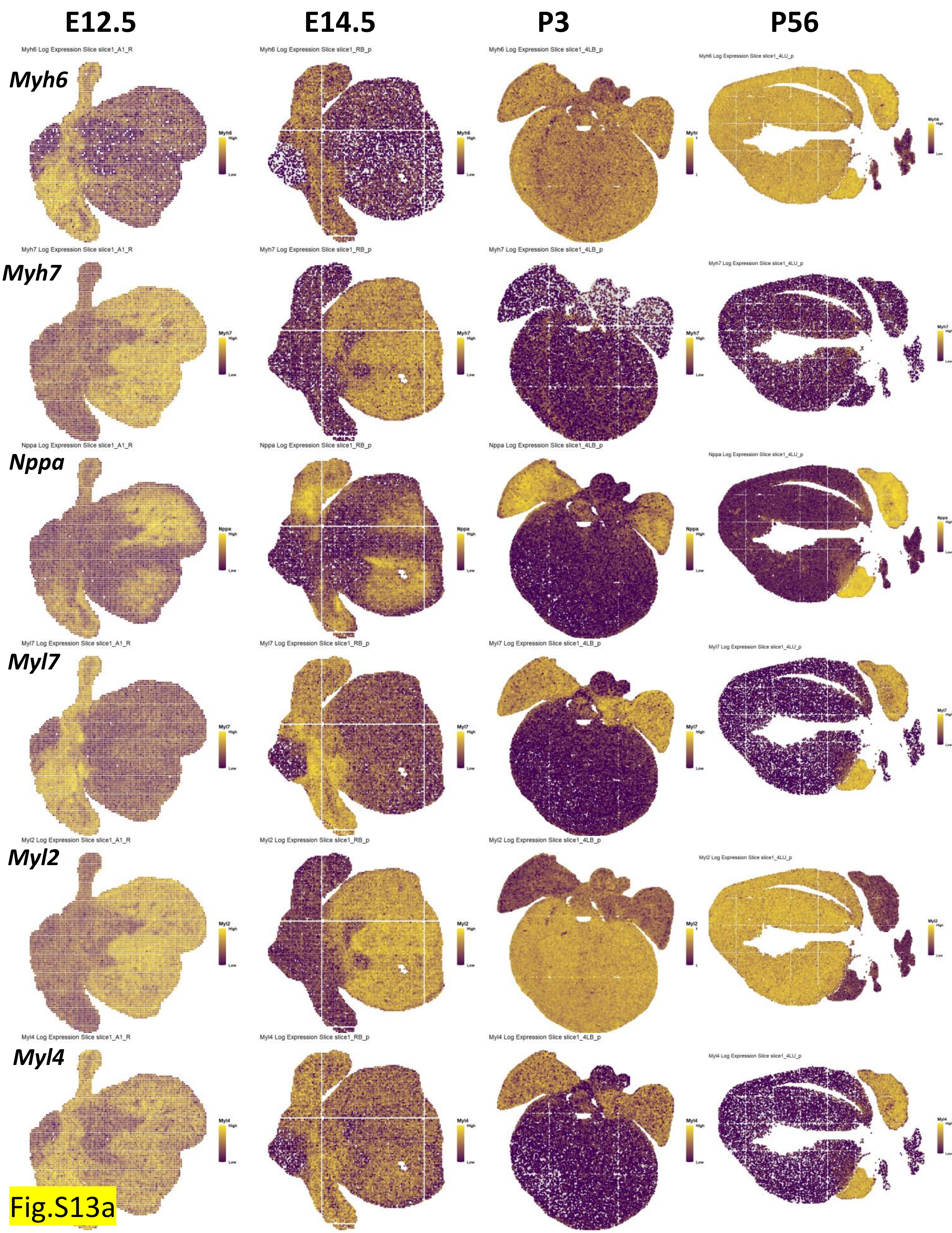
E12.5
E14.5
P3
P56
Myh6
Myh7
Nppa
Myl7
Myl2
Myl4
Myh6 Log Expression Slice slice1_A1_R
Myh6 Log Expression Slice slice1_RB_p
Myh6 Log Expression Slice slice1_4LB_p
Myh6 Log Expression Slice slice1_4LU_p
Myh7 Log Expression Slice slice1_A1_R
Myh7 Log Expression Slice slice1_RB_p
Myh7 Log Expression Slice slice1_4LB_p
Myh7 Log Expression Slice slice1_4LU_p
Nppa Log Expression Slice slice1_A1_R
Nppa Log Expression Slice slice1_RB_p
Nppa Log Expression Slice slice1_4LB_p
Nppa Log Expression Slice slice1_4LU_p
Myl7 Log Expression Slice slice1_A1_R
Myl7 Log Expression Slice slice1_RB_p
Myl7 Log Expression Slice slice1_4LB_p
Myl7 Log Expression Slice slice1_4LU_p
Myl2 Log Expression Slice slice1_A1_R
Myl2 Log Expression Slice slice1_RB_p
Myl2 Log Expression Slice slice1_4LB_p
Myl2 Log Expression Slice slice1_4LU_p
Myl4 Log Expression Slice slice1_A1_R
Myl4 Log Expression Slice slice1_RB_p
Myl4 Log Expression Slice slice1_4LB_p
Myl4 Log Expression Slice slice1_4LU_p
High
Low

Fig.S13a

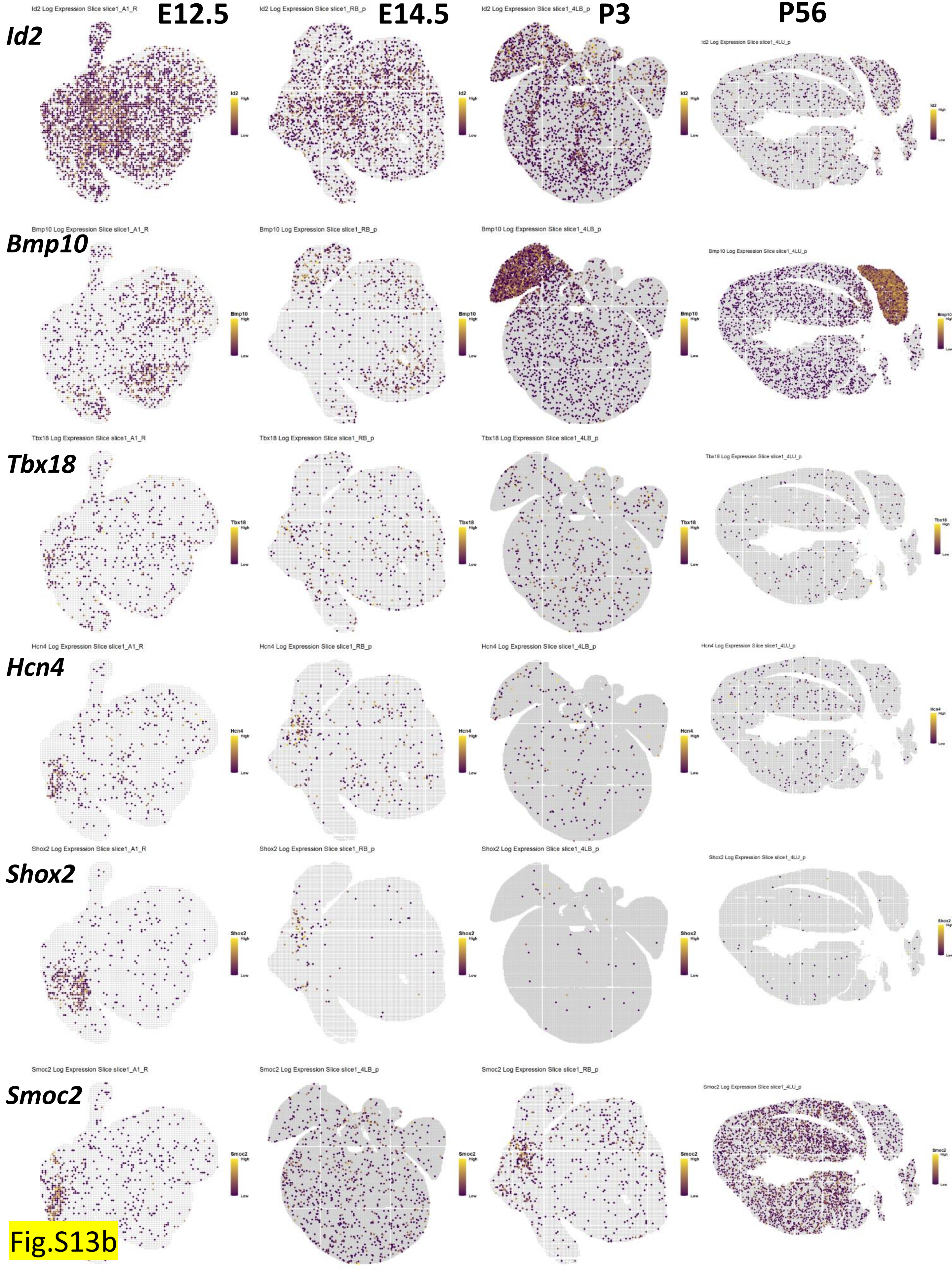

E12.5
E14.5
P3
P56
Id2
Bmp10
Tbx18
Hcn4
Shox2
Smoc2
Id2 Log Expression Slice slice1_A1_R
Id2 Log Expression Slice slice1_RB_p
Id2 Log Expression Slice slice1_4LB_p
Id2 Log Expression Slice slice1_4LU_p
Bmp10 Log Expression Slice slice1_A1_R
Bmp10 Log Expression Slice slice1_RB_p
Bmp10 Log Expression Slice slice1_4LB_p
Bmp10 Log Expression Slice slice1_4LU_p
Tbx18 Log Expression Slice slice1_A1_R
Tbx18 Log Expression Slice slice1_RB_p
Tbx18 Log Expression Slice slice1_4LB_p
Tbx18 Log Expression Slice slice1_4LU_p
Hcn4 Log Expression Slice slice1_A1_R
Hcn4 Log Expression Slice slice1_RB_p
Hcn4 Log Expression Slice slice1_4LB_p
Hcn4 Log Expression Slice slice1_4LU_p
Shox2 Log Expression Slice slice1_A1_R
Shox2 Log Expression Slice slice1_RB_p
Shox2 Log Expression Slice slice1_4LB_p
Shox2 Log Expression Slice slice1_4LU_p
Smoc2 Log Expression Slice slice1_A1_R
Smoc2 Log Expression Slice slice1_4LB_p
Smoc2 Log Expression Slice slice1_RB_p
Smoc2 Log Expression Slice slice1_4LU_p
High
Low


Fig.S13b

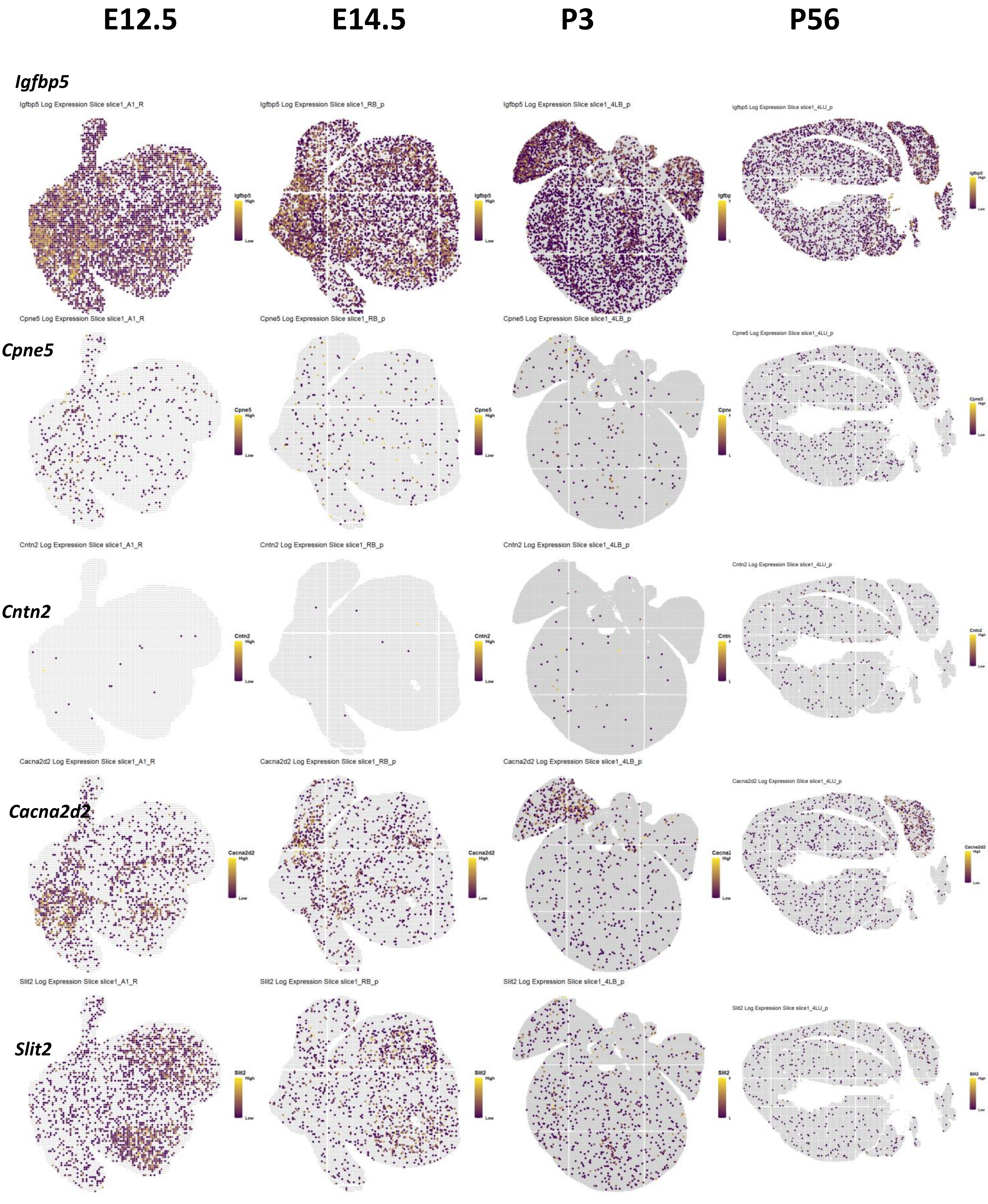
E12.5
E14.5
P3
P56
Igfbp5
Igfbp5 Log Expression Slice slice1_A1_R
Igfbp5 Log Expression Slice slice1_RB_p
Igfbp5 Log Expression Slice slice1_4LB_p
Igfbp5 Log Expression Slice slice1_4LU_p
Cpne5
Cpne5 Log Expression Slice slice1_A1_R
Cpne5 Log Expression Slice slice1_RB_p
Cpne5 Log Expression Slice slice1_4LB_p
Cpne5 Log Expression Slice slice1_4LU_p
Cntn2
Cntn2 Log Expression Slice slice1_A1_R
Cntn2 Log Expression Slice slice1_RB_p
Cntn2 Log Expression Slice slice1_4LB_p
Cntn2 Log Expression Slice slice1_4LU_p
Cacna2d2
Cacna2d2 Log Expression Slice slice1_A1_R
Cacna2d2 Log Expression Slice slice1_RB_p
Cacna2d2 Log Expression Slice slice1_4LB_p
Cacna2d2 Log Expression Slice slice1_4LU_p
Slit2
Slit2 Log Expression Slice slice1_A1_R
Slit2 Log Expression Slice slice1_RB_p
Slit2 Log Expression Slice slice1_4LB_p
Slit2 Log Expression Slice slice1_4LU_p
High
Low

Fig.S13c

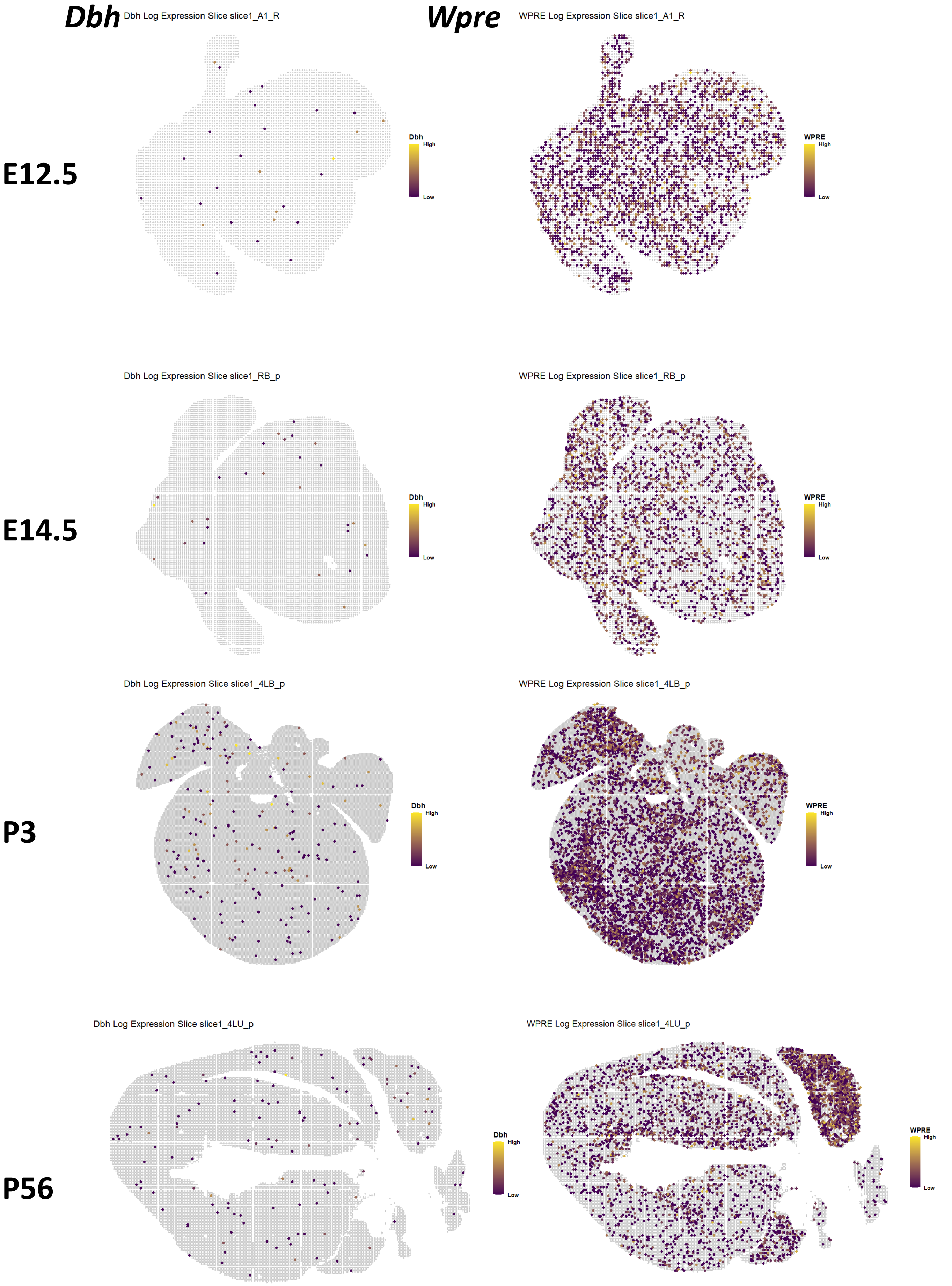

Dbh
Wpre
E12.5
E14.5
P3
P56
Dbh Log Expression Slice slice1_A1_R
WPRE Log Expression Slice slice1_A1_R
Dbh Log Expression Slice slice1_RB_p
WPRE Log Expression Slice slice1_RB_p
Dbh Log Expression Slice slice1_4LB_p
WPRE Log Expression Slice slice1_4LB_p
Dbh Log Expression Slice slice1_4LU_p
WPRE Log Expression Slice slice1_4LU_p
Dbh
WPRE
High
Low


Fig.S13d

**a**

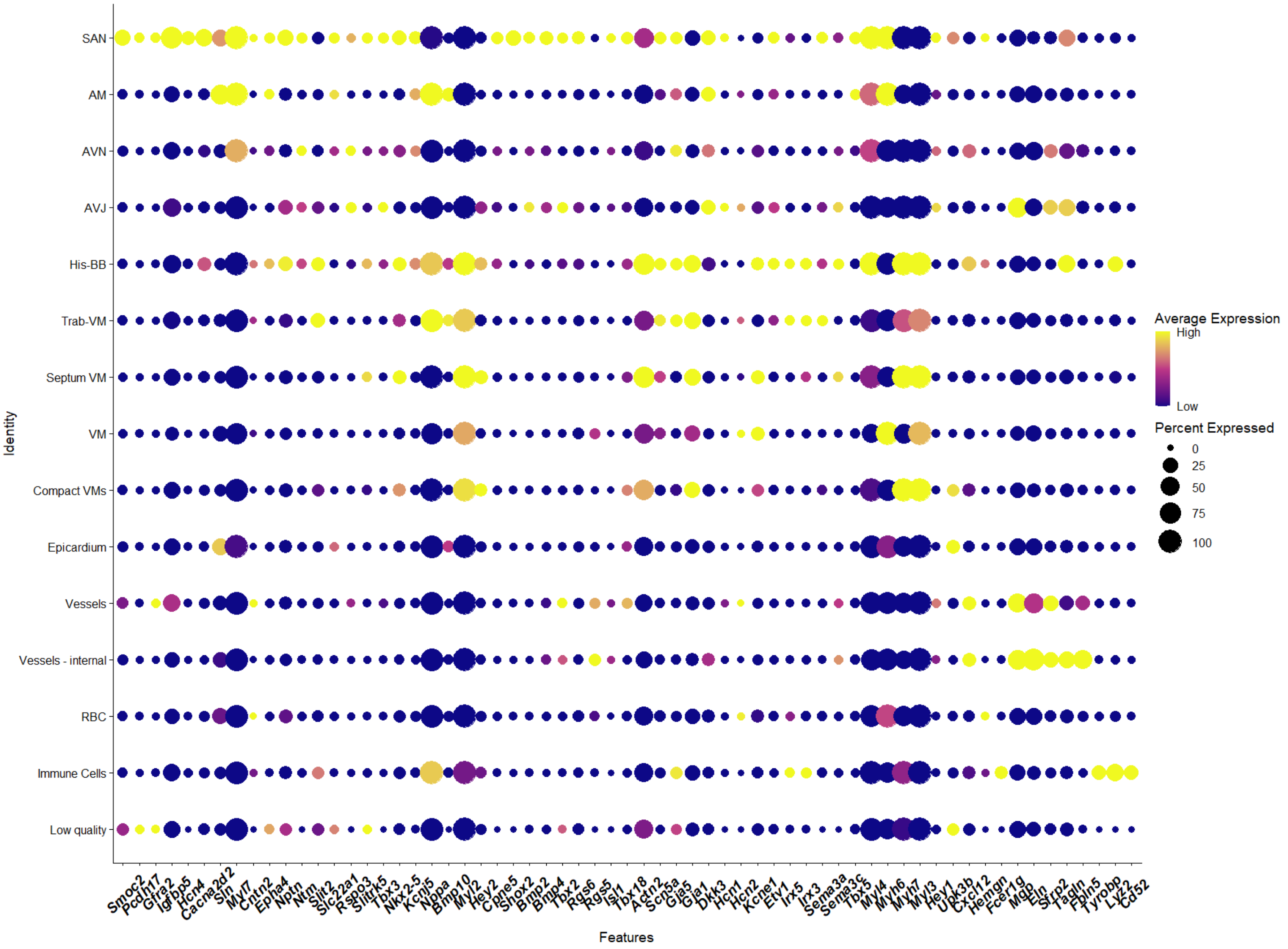

SAN
AM
AVN
AVJ
His-BB
Trab-VM
Septum VM
VM
Compact VMs
Epicardium
Vessels
Vessels - internal
RBC
Immune Cells
Low quality
Identity
Smoc2
Pcdh17
Gfra2
Igfbp5
Hcn4
Cacna2d2
Sln
Myl7
Cntn2
Epha4
Nptn
Ntm
Slit2
Slc22a1
Rspo3
Slitrk5
Tbx3
Nkx2-5
Kcnj5
Nppa
Bmp10
Myl2
Hey2
Cpne5
Shox2
Bmp2
Bmp4
Tbx2
Rgs6
Rgs5
Isl1
Tbx18
Actn2
Scn5a
Gja5
Gja1
Dkk3
Hcn1
Hcn2
Kcne1
Etv1
Irx5
Irx3
Sema3a
Sema3c
Tbx5
Myl4
Myh6
Myh7
Myl3
Hey1
Upk3b
Cxcl12
Hemgn
Fcer1g
Mgp
Eln
Sfrp2
Tagln
Fbln5
Tyrobp
Lyz2
Cd52
Features
Average Expression
High
Low
Percent Expressed
0
25
50
75
100


Fig.S14

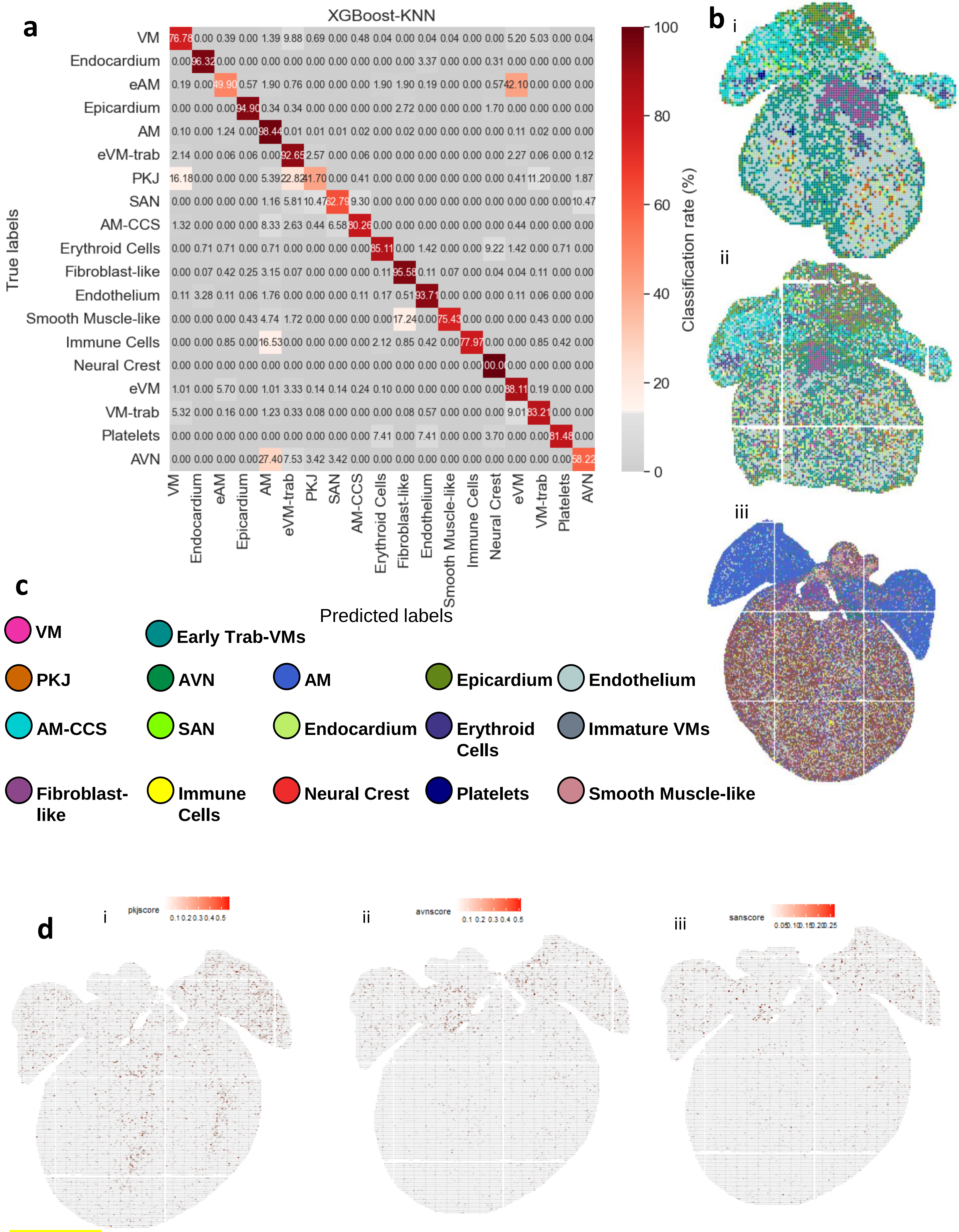
a
XGBoost-KNN
True labels
VM 76.78 0.00 0.39 0.00 1.39 9.88 0.69 0.00 0.48 0.04 0.00 0.04 0.04 0.00 0.00 5.20 5.03 0.00 0.04
Endocardium 0.00 96.32 0.00 0.00 0.00 0.00 0.00 0.00 0.00 0.00 0.00 3.37 0.00 0.00 0.31 0.00 0.00 0.00 0.00
eAM 0.19 0.00 49.90 0.57 1.90 0.76 0.00 0.00 0.00 1.90 1.90 0.19 0.00 0.00 0.57 42.10 0.00 0.00 0.00
Epicardium 0.00 0.00 0.00 94.90 0.34 0.34 0.00 0.00 0.00 0.00 2.72 0.00 0.00 0.00 1.70 0.00 0.00 0.00 0.00
AM 0.10 0.00 1.24 0.00 98.44 0.01 0.01 0.01 0.02 0.00 0.02 0.00 0.00 0.00 0.00 0.11 0.02 0.00 0.00
eVM-trab 2.14 0.00 0.06 0.06 0.00 92.65 2.57 0.00 0.06 0.00 0.00 0.00 0.00 0.00 0.00 2.27 0.06 0.00 0.12
PKJ 16.18 0.00 0.00 0.00 5.39 22.82 41.70 0.00 0.41 0.00 0.00 0.00 0.00 0.00 0.00 0.41 11.20 0.00 1.87
SAN 0.00 0.00 0.00 0.00 1.16 5.81 10.47 62.79 9.30 0.00 0.00 0.00 0.00 0.00 0.00 0.00 0.00 0.00 10.47
AM-CCS 1.32 0.00 0.00 0.00 8.33 2.63 0.44 6.58 80.26 0.00 0.00 0.00 0.00 0.00 0.00 0.44 0.00 0.00 0.00
Erythroid Cells 0.00 0.71 0.71 0.00 0.71 0.00 0.00 0.00 0.00 85.11 0.00 1.42 0.00 0.00 9.22 1.42 0.00 0.71 0.00
Fibroblast-like 0.00 0.07 0.42 0.25 3.15 0.07 0.00 0.00 0.00 0.11 95.58 0.11 0.07 0.00 0.04 0.04 0.11 0.00 0.00
Endothelium 0.11 3.28 0.11 0.06 1.76 0.00 0.00 0.00 0.11 0.17 0.51 93.71 0.00 0.00 0.00 0.11 0.06 0.00 0.00
Smooth Muscle-like 0.00 0.00 0.00 0.43 4.74 1.72 0.00 0.00 0.00 0.00 17.24 0.00 75.43 0.00 0.00 0.00 0.43 0.00 0.00
Immune Cells 0.00 0.00 0.85 0.00 16.53 0.00 0.00 0.00 0.00 2.12 0.85 0.42 0.00 77.97 0.00 0.00 0.85 0.42 0.00
Neural Crest 0.00 0.00 0.00 0.00 0.00 0.00 0.00 0.00 0.00 0.00 0.00 0.00 0.00 0.00 00.0 0.00 0.00 0.00 0.00
eVM 1.01 0.00 5.70 0.00 1.01 3.33 0.14 0.14 0.24 0.10 0.00 0.00 0.00 0.00 0.00 88.11 0.19 0.00 0.00
VM-trab 5.32 0.00 0.16 0.00 1.23 0.33 0.08 0.00 0.00 0.00 0.08 0.57 0.00 0.00 0.00 9.01 83.21 0.00 0.00
Platelets 0.00 0.00 0.00 0.00 0.00 0.00 0.00 0.00 0.00 7.41 0.00 7.41 0.00 0.00 3.70 0.00 0.00 81.48 0.00
AVN 0.00 0.00 0.00 0.00 27.40 7.53 3.42 3.42 0.00 0.00 0.00 0.00 0.00 0.00 0.00 0.00 0.00 0.00 58.22
VM Endocardium eAM Epicardium AM eVM-trab PKJ SAN AM-CCS Erythroid Cells Fibroblast-like Endothelium Smooth Muscle-like Immune Cells Neural Crest eVM VM-trab Platelets AVN
Predicted labels
100 80 60 40 20 0
Classification rate (%)
b
i
ii
iii
c
VM
Early Trab-VMs
PKJ
AVN
AM
Epicardium
Endothelium
AM-CCS
SAN
Endocardium
Erythroid Cells
Immature VMs
Fibroblast-like
Immune Cells
Neural Crest
Platelets
Smooth Muscle-like
d
i
pkjscore
0.1 0.2 0.3 0.4 0.5
ii
avnscore
0.1 0.2 0.3 0.4 0.5
iii
sanscore
0.05 0.10 0.15 0.20 0.25

Fig.S15

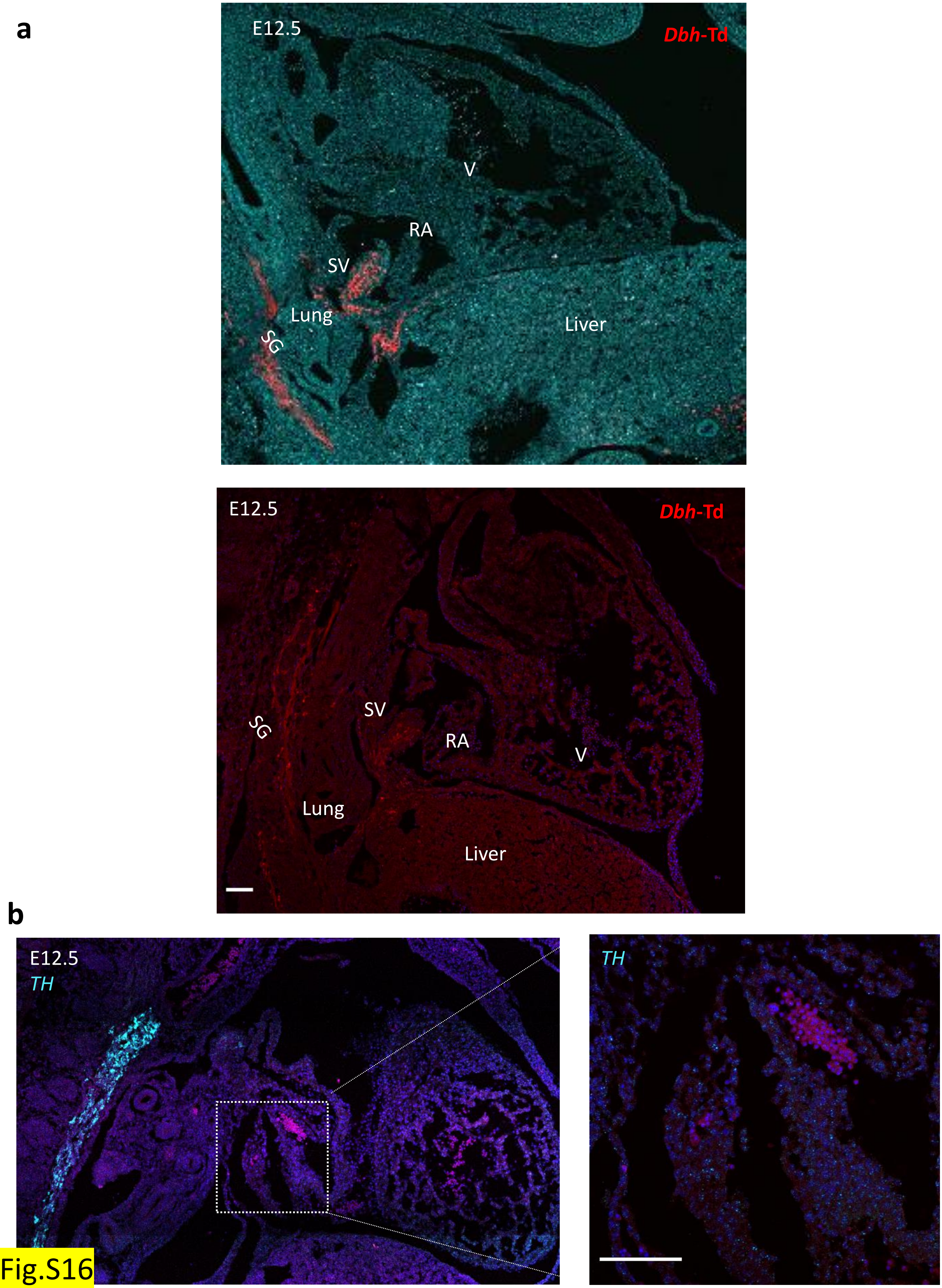
a
E12.5
Dbh-Td
V
RA
SV
Lung
SG
Liver
E12.5
Dbh-Td
SV
SG
RA
V
Lung
Liver
b
E12.5
TH
TH

Fig.S16

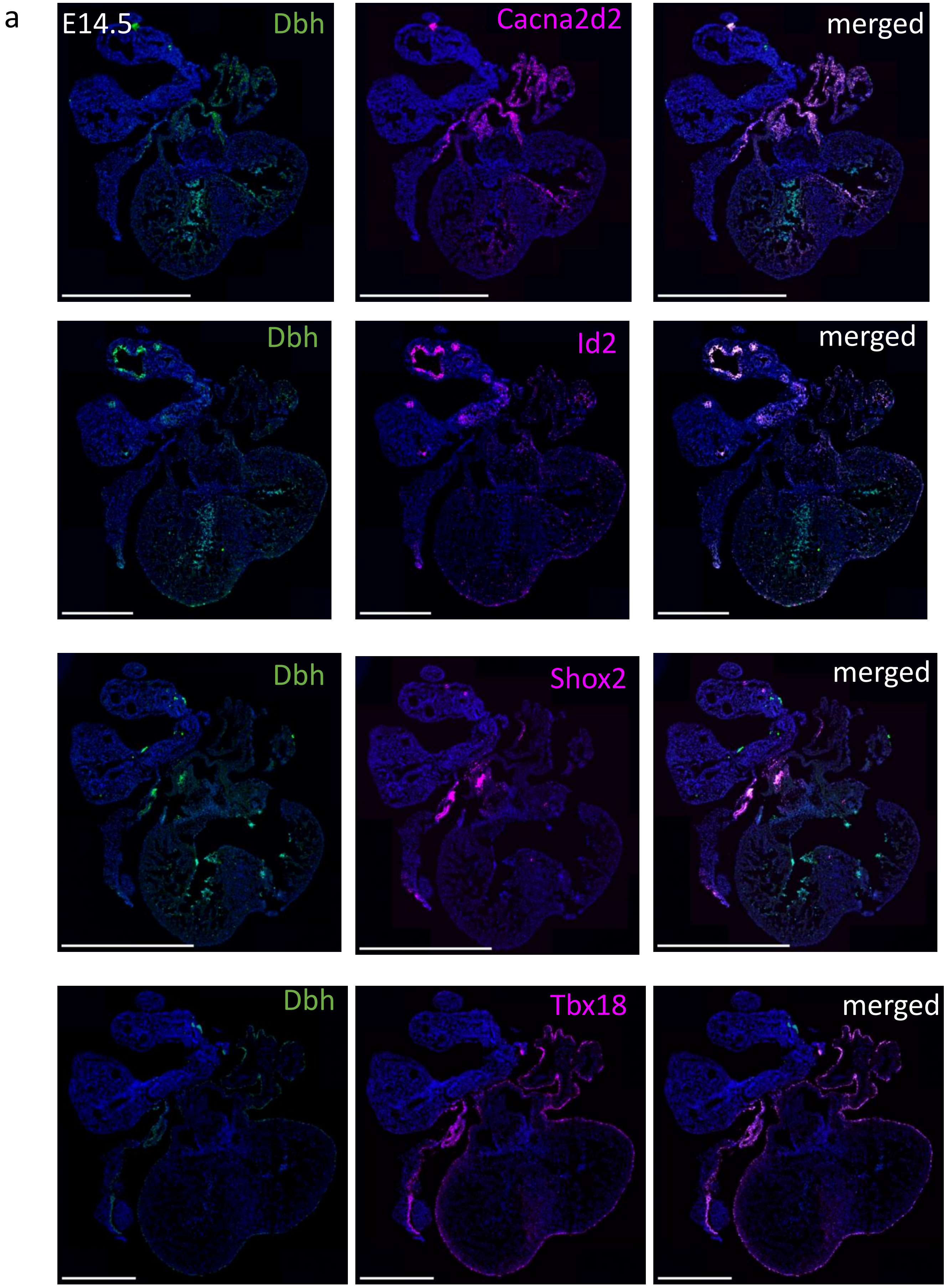
a
E14.5
Dbh
Cacna2d2
merged
Dbh
Id2
merged
Dbh
Shox2
merged
Dbh
Tbx18
merged

Fig.S17a

b

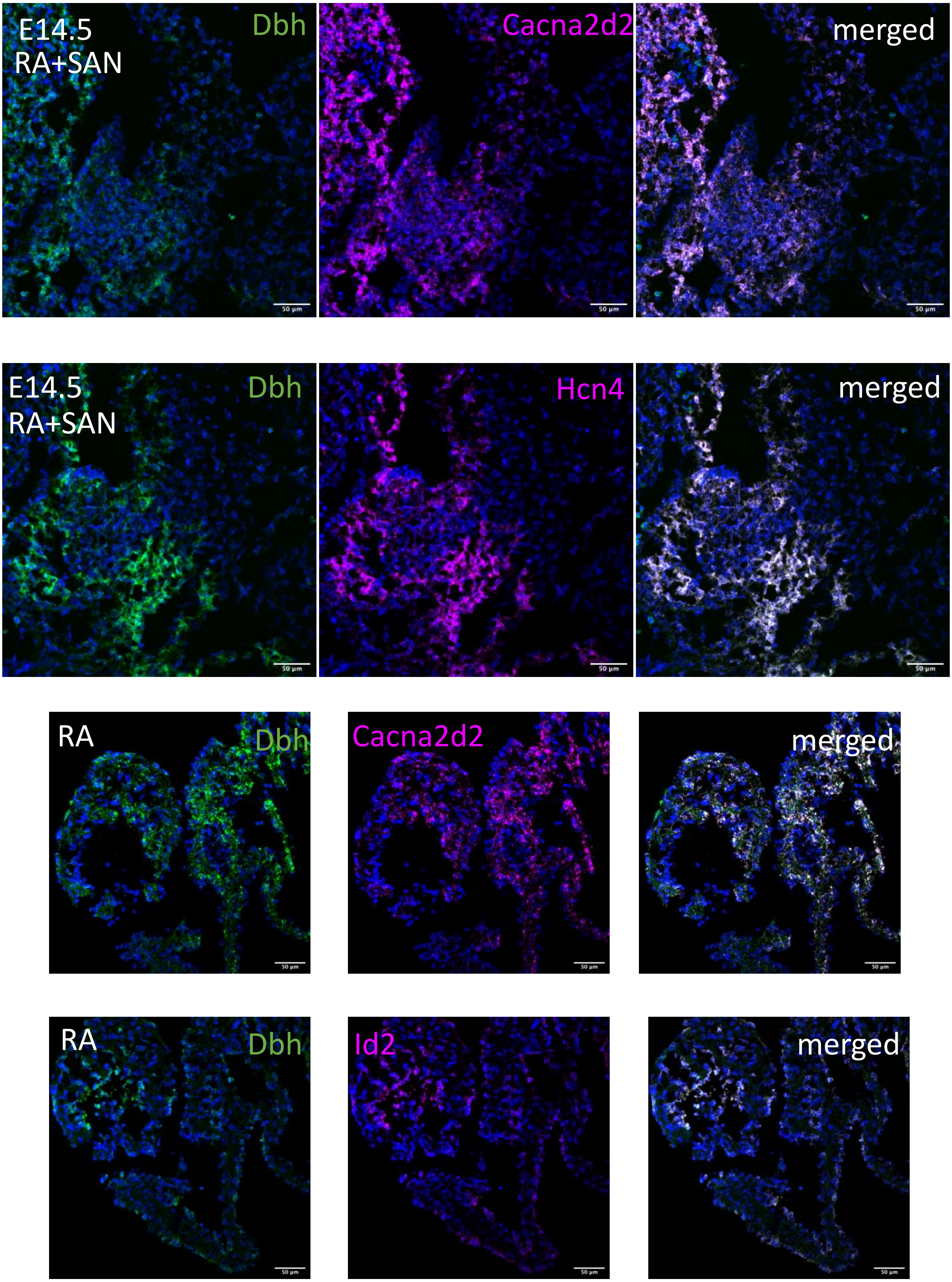
E14.5
RA+SAN
Dbh
Cacna2d2
merged
E14.5
RA+SAN
Dbh
Hcn4
merged
RA
Dbh
Cacna2d2
merged
RA
Dbh
Id2
merged

Fig.S17b

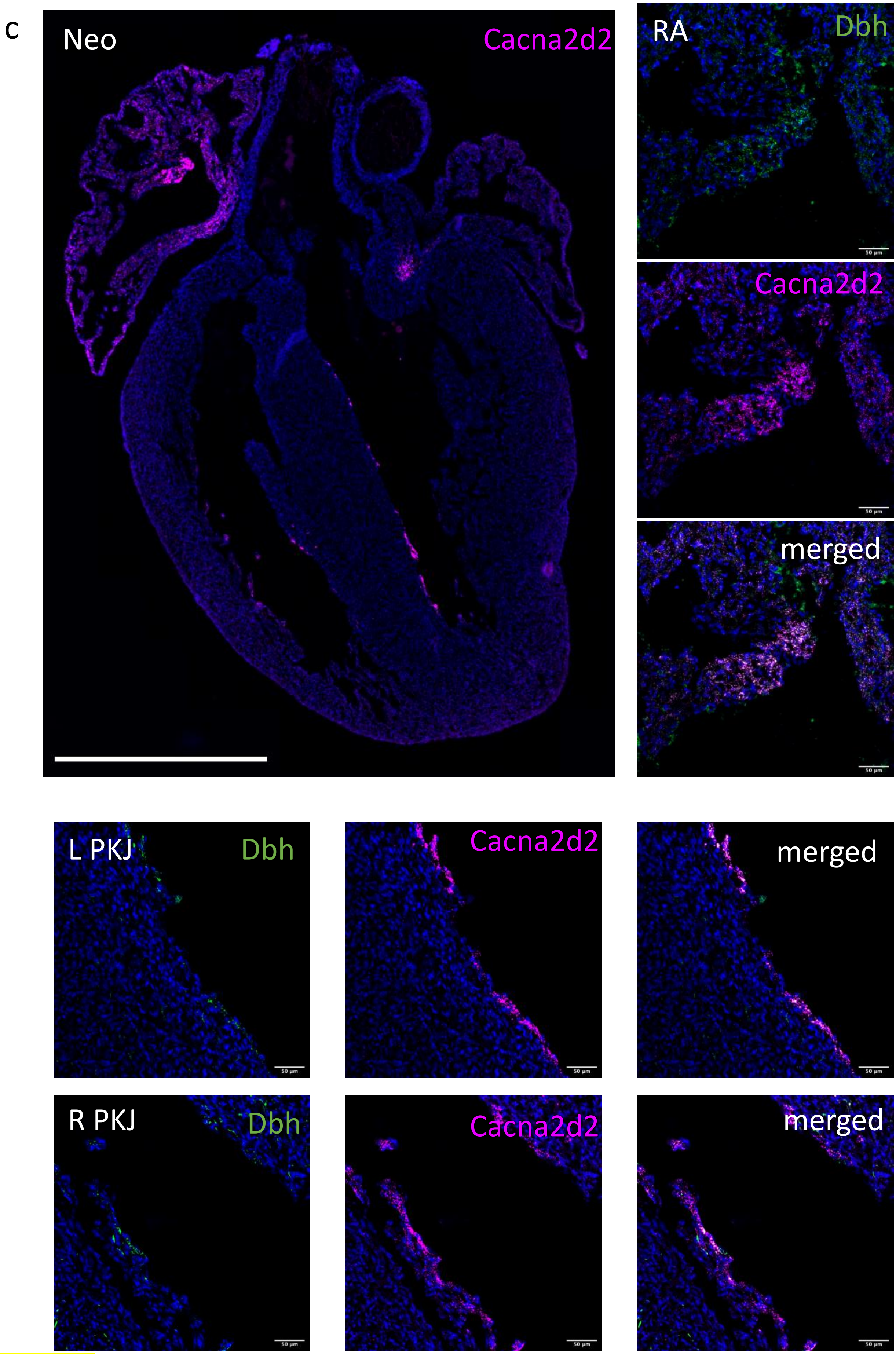
c
Neo
Cacna2d2
RA
Dbh
Cacna2d2
merged
L PKJ
Dbh
Cacna2d2
merged
R PKJ
Dbh
Cacna2d2
merged

Fig.S17c

**a**

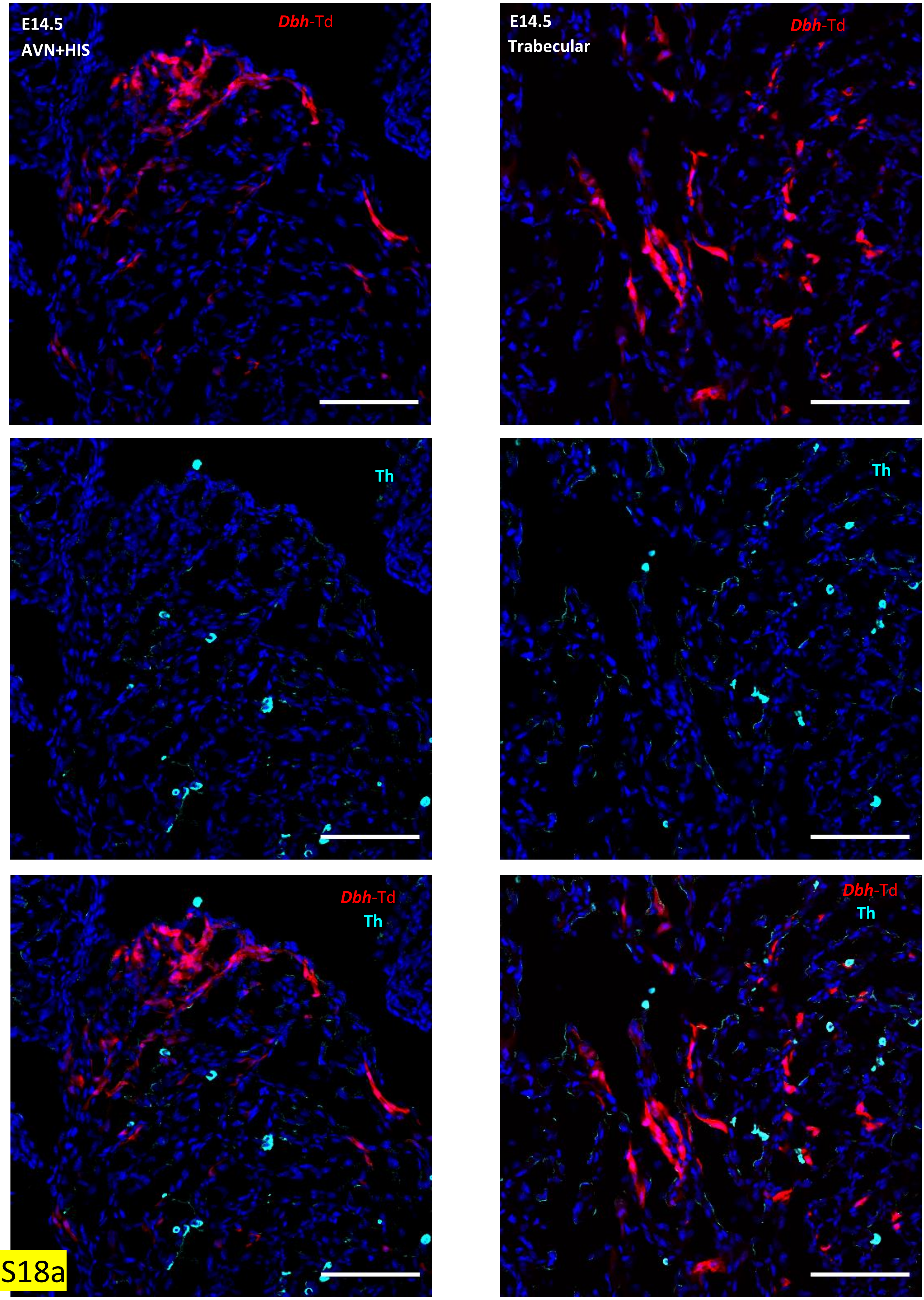

E14.5
AVN+HIS
*Dbh*-Td
E14.5
Trabecular
*Dbh*-Td
Th
Th
*Dbh*-Td
Th
*Dbh*-Td
Th


Fig.S18a

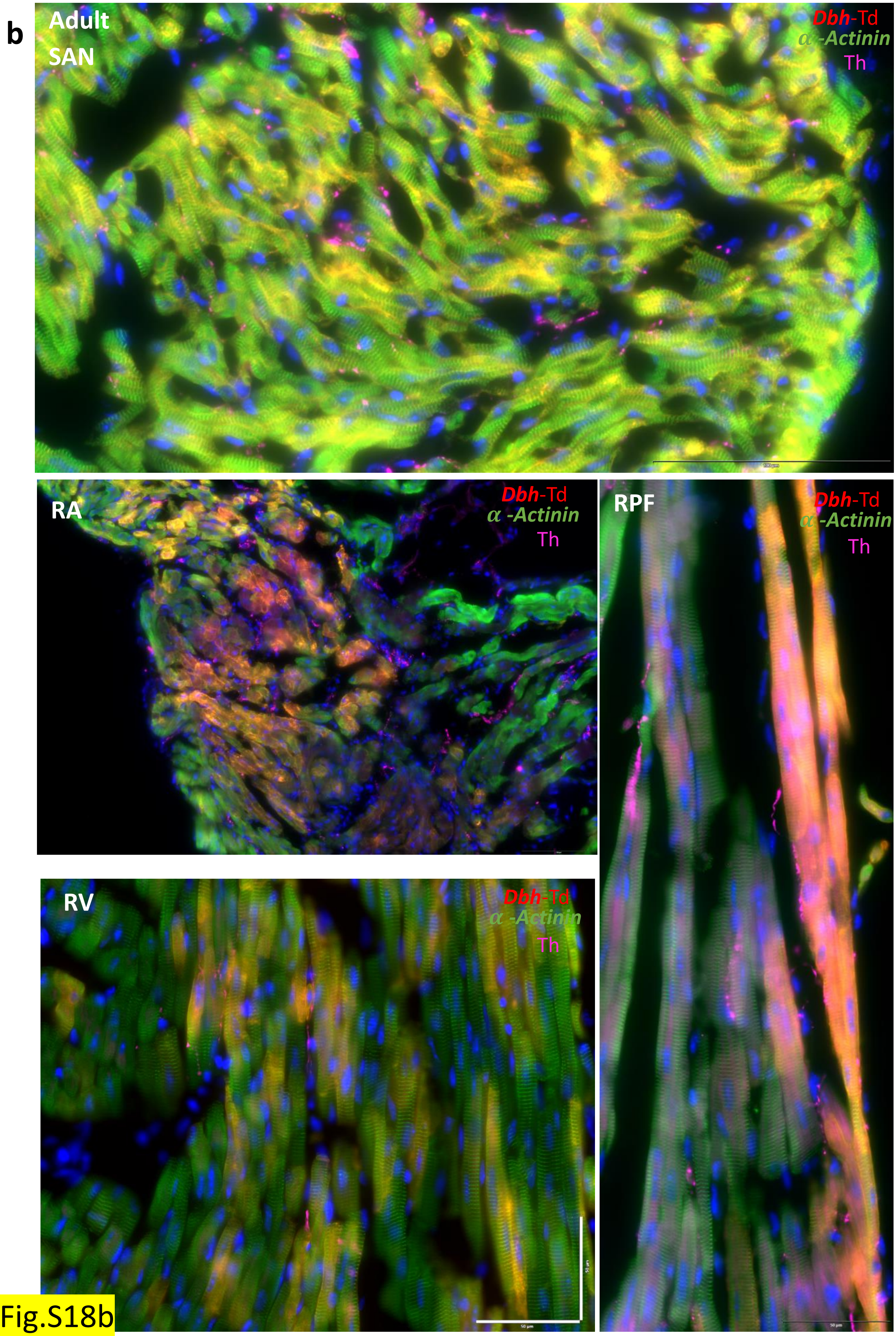
b
Adult
SAN
Dbh-Td
α -Actinin
Th
RA
Dbh-Td
α -Actinin
Th
RPF
Dbh-Td
α -Actinin
Th
RV
Dbh-Td
α -Actinin
Th
50 μm
50 μm
50 μm

Fig.S18b

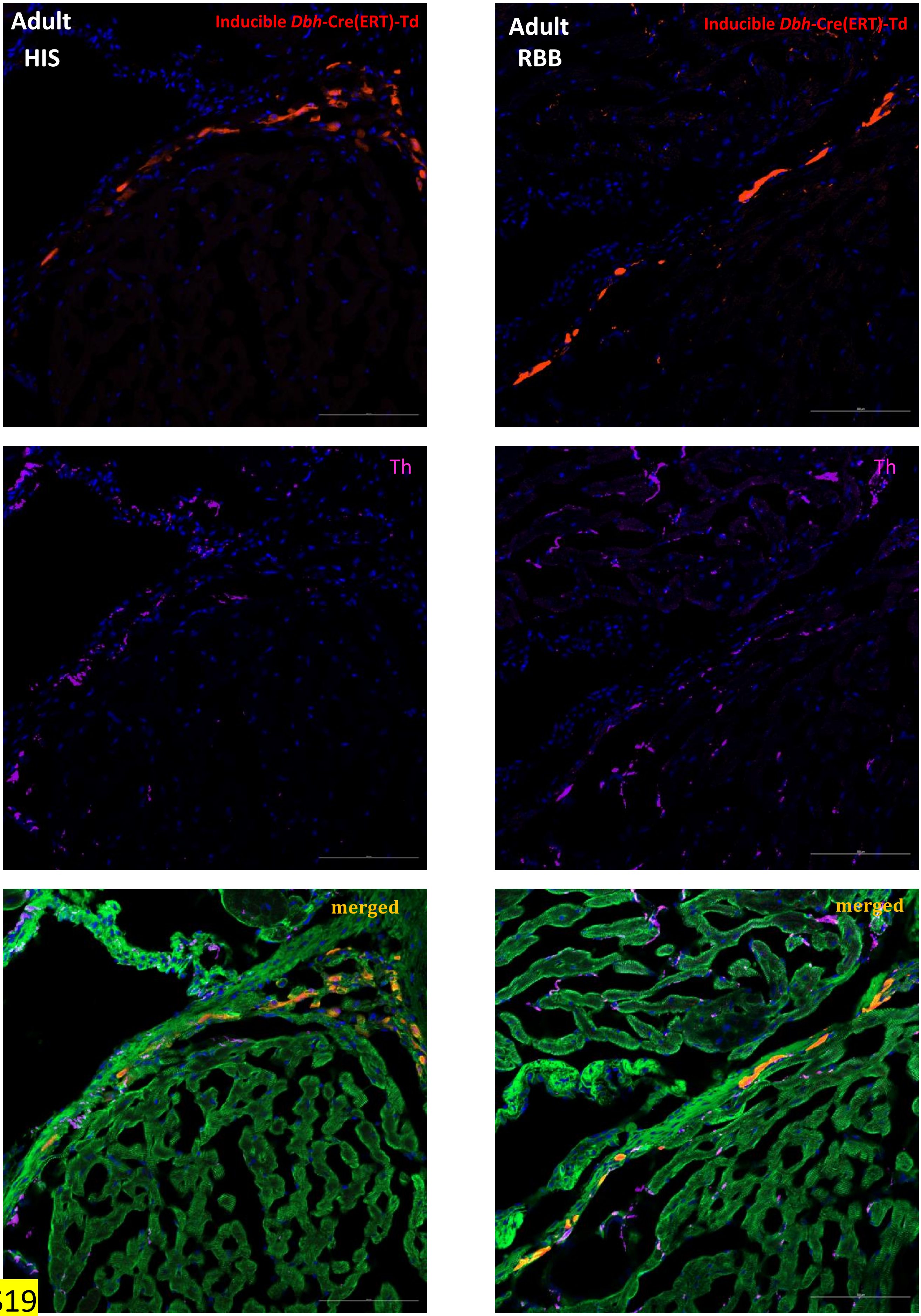

Adult
HIS
Inducible Dbh-Cre(ERT)-Td
Adult
RBB
Inducible Dbh-Cre(ERT)-Td
Th
Th
merged
merged


Fig.S19
